# Organizing neuronal ER-PM junctions is a conserved nonconducting function of Kv2 plasma membrane ion channels

**DOI:** 10.1101/296731

**Authors:** Michael Kirmiz, Stephanie Palacio, Parashar Thapa, Anna N. King, Jon T. Sack, James S. Trimmer

**Affiliations:** Departments of Neurobiology, Physiology and Behavior, University of California, Davis, CA 95616; Departments of Physiology and Membrane Biology, University of California, Davis, CA 95616; Anesthesiology and Pain Medicine, University of California, Davis, CA 95616

## Abstract

Endoplasmic reticulum (ER) and plasma membrane (PM) form junctions crucial to ion and lipid signaling and homeostasis. The Kv2.1 ion channel is unique among PM proteins in organizing ER-PM junctions. Here, we show that this organizing function is conserved between Kv2 family members that differ in their biophysical properties, modulation and cellular expression. Manipulation of actin cytoskeleton surrounding Kv2 ER-PM junctions affects their spatial organization. Kv2-containing ER-PM junctions overlap with those formed by canonical ER-PM tethers. ER-PM junction organization by Kv2 channels is unchanged by point mutations that eliminate ion conduction, but abolished by those that eliminate PM clustering without impacting ion channel function. Kv2.2 is distinct in lacking the reversible modulation of junction organization present in Kv2.1. Brain neurons in Kv2 double knockout mice have altered ER-PM junctions, demonstrating a conserved *in vivo* function for Kv2 family members distinct from their canonical role as ion-conducting channels shaping neuronal excitability.

## Introduction

Membrane contacts between the endoplasmic reticulum (ER) and plasma membrane (PM), or ER-PM junctions, are a ubiquitous feature of eukaryotic cells (1-4). These specialized sites at which ER is held in close apposition (10-30 nm) to the PM represent critical platforms for mediating ER and PM lipid metabolism and transport and as hubs for Ca^2+^ homeostasis and signaling events (5, 6). ER-PM junctions are classified according to the resident ER protein serving as the PM tether and that are members of the Extended Synaptotagmin or E-Syt (7), Junctophilin or JP (8), or the Stromal Interacting Molecule or STIM (9) families. These otherwise unrelated ER membrane proteins have a common membrane topology with a large cytoplasmic domain that mediates binding to specific classes of phospholipids in the inner leaflet of the PM (1, 10). The STIM proteins can also reversibly bind to PM Orai proteins in a process triggered by ER Ca^2+^ depletion (9). While mRNA measurements have shown that many of these ER-localized tethering proteins have high levels of expression in brain [*e.g.*, (8, 11-13)], little is known of the subcellular localization of these proteins relative to the different classes of ER-PM junctions that have been observed in ultrastructural studies of brain neurons (14-16).

Plasma membrane voltage-gated K^+^ or Kv channels play crucial yet diverse roles in shaping neuronal function (17). Among these, the Kv2 family contains two members: Kv2.1 and Kv2.2. Like other Kv channels, Kv2.1 and Kv2.2 are key determinants of action potential characteristics and intrinsic electrical excitability in distinct classes of mammalian brain neurons (18-25), and *de novo* mutations in Kv2.1 are associated with devastating neonatal encephalopathic epilepsies and neurodevelopmental delays (26-29). Kv2 channels are also prominently yet differentially expressed in pancreatic islets (30, 31), smooth muscle cells (32, 33), and other excitable and non-excitable cell types. In brain neurons, Kv2 channels are distinct from other Kv channels (17) in being specifically localized to high-density micron sized clusters prominent on the soma, proximal dendrites, and axon initial segment (34-42). Kv2 channels also form such clusters when exogenously expressed in cultured neurons and in heterologous cells (35, 37, 39, 41-47). A short proximal restriction and clustering (PRC) motif in the relatively large cytoplasmic C-terminus of Kv2.1 is necessary for its clustered localization in neurons and heterologous cells (35, 37), and is sufficient to transfer Kv2.1-like clustering to other Kv channels (37, 45). Mutations within the PRC motif in Kv2.2 result in loss of Kv2.2 clustering (39).

Immunoelectron microscopy-based studies have shown that immunoreactivity for PM Kv2.1 (38, 40, 41) and Kv2.2 (39) are associated with subsurface cisternae, a form of ER-PM junctions that are prominent in somata of brain neurons (14-16). In certain brain neurons, clusters of PM Kv2.1 channels overlay clusters of ER-localized ryanodine receptor (RyR) Ca^2+^ release channels (38, 48) which are concentrated at ER-PM junctions to mediate local Ca^2+^ signaling events in diverse cell types (49, 50). Recent studies revealed that in addition to being localized to ER-PM junctions, exogenous expression of Kv2.1 leads to recruitment and/or stabilization of ER-PM junctions in heterologous cells and cultured hippocampal neurons or CHNs (51). The ability of Kv2.1 to organize ER-PM junctions exhibits the same phosphorylation-dependent regulation as Kv2.1 clustering (47), which is regulated by numerous stimuli that impact Kv2.1 phosphorylation state (39, 41, 52-54). It is not known whether the changes in ER-PM junction structure upon heterologous expression of Kv2.1 are a result of the channel’s K^+^ conductance with a subsequent impact on membrane potential or K^+^ concentration, or whether it is through a more direct structural role. Kv2.1 and Kv2.2 share 61% overall amino acid identity (39% in their respective cytoplasmic C-termini that comprises about half of their primary structure), and have distinct biophysical properties [*e.g.*, (55, 56)] and expression patterns [*e.g.*, (18, 31, 39, 41, 57-59)]. Moreover, stimuli that trigger reversible modulation of voltage activation [*e.g.,* (38, 39, 56)] and dispersal of clustering (39) of Kv2.1 do not detectably impact Kv2.2, leading to questions as to whether Kv2.2 is also distinct from Kv2.1 in its ability to organize ER-PM junctions. Lastly, it is not known whether altering expression of endogenous Kv2 channels affects ER-PM junctions in brain neurons *in situ*. Here, we define the localization of Kv2.2 relative to ER-PM junctions in brain neurons *in situ* and in culture and determine its role in organizing ER-PM junctions. We determine the relationship of Kv2-containing ER-PM junctions to the actin cytoskeleton and to other classes of molecular-defined ER-PM junctions. We employ a strategic set of point mutations in Kv2.2 to separately determine the contributions of K^+^ conduction and clustering to the remodeling of neuronal ER-PM junctions. We also determine how the differential regulation of Kv2.1 and Kv2.2 clustering impacts the associated ER-PM junctions. Finally, we use recently generated double knockout mice lacking expression of both mammalian Kv2 channel family members to determine their *in vivo* role in organizing ER-PM junctions in brain neurons *in situ*. Our results provide compelling evidence for a conserved role for nonconducting Kv2 channels in organizing ER-PM junctions in brain neurons and other cell types in which these ion channels are abundantly expressed.

## Results

### Plasma membrane clusters of Kv2.2 associate with ER-PM junctions in mammalian brain neurons *in situ* and in culture, and in heterologous HEK293T cells

Kv2.2 is present in clusters on the soma, proximal dendrites and axon initial segments of mammalian brain neurons (39, 41, 42). To investigate the subcellular localization of these Kv2.2 clusters relative to native ER-PM junctions in brain neurons, we performed multiplex immunofluorescence labeling for PM Kv2.2 and ER-localized RyR Ca^2+^ release channels, which are concentrated at ER-PM junctions. In mouse brain sections, somatic Kv2.2 clusters were found at/near RyR clusters in specific neuron types, including hippocampal CA1 pyramidal neurons and layer 6 neocortical neurons (Figure 1). A similar juxtaposition of Kv2.2 and RyR clusters was seen in CHNs (Figure 1). In these classes of neurons, Kv2.2 was often found coclustered with Kv2.1 at these ER-PM junctions (Figure 1). Neurons in each preparation also contained RyR clusters that did not appear to colocalize with Kv2.2 or Kv2.1 (Figure 1). These findings demonstrate that Kv2.2 clusters localize to RyR-containing ER-PM junctions in intact mammalian brain neurons *in situ* and in culture.

**Figure 1.**
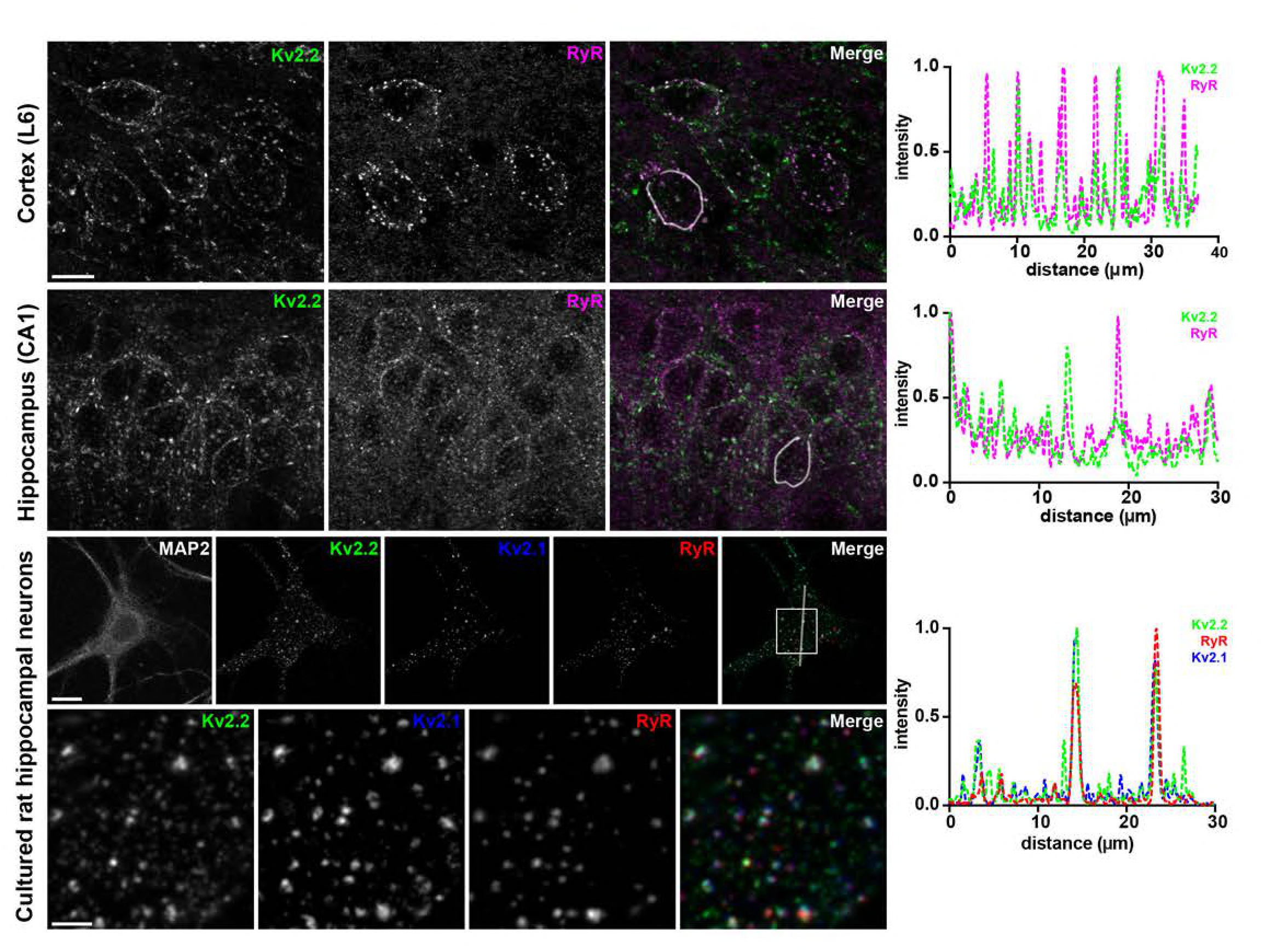
Endogenous Kv2.2 associates with RyR-containing ER-PM junctions in brain neurons *in situ* and in culture. Projected z-stack images of multiplex immunofluorescence labeling of adult mouse neocortex and hippocampal CA1 region, and CHNs, for Kv2.2 and RyR, or Kv2.2, RyR and Kv2.1, as indicated. Scale bar in Kv2.2 neocortex panel is 10 µm and holds for all brain panels. Scale bar in MAP2 CHN panel is 10 µm and holds for all CHN panels in that row. Image exposure time was optimized for the labeling of each brain region independently. Scale bar in Kv2.2 magnified inset is 2.5 µm and holds for all panels in that row. Panels to the right of each set of images are the corresponding normalized fluorescence intensity values across the individual line scans depicted by the white line in the merged images.

We next determined whether heterologously expressed and clustered Kv2.2 localizes more generally to ER-PM junctions. In HEK293T cells coexpressing GFP-tagged Kv2.2 and BFP-tagged SEC61β [a general ER marker; (60)], optical sections taken through the center of cells show fingerlike projections of SEC61β-positive ER, a subset of which were associated with PM Kv2.2 clusters, which appear as discrete PM segments (Figure 2). Three-dimensional reconstructions show that the ER projections terminating at Kv2.2-associated PM clusters were contiguous with bulk ER (Figure 2, Movie 1). Together these results suggest that Kv2.2 localizes to ER-PM junctions in mammalian brain neurons and when heterologously expressed in HEK293T cells.

**Figure 2.**
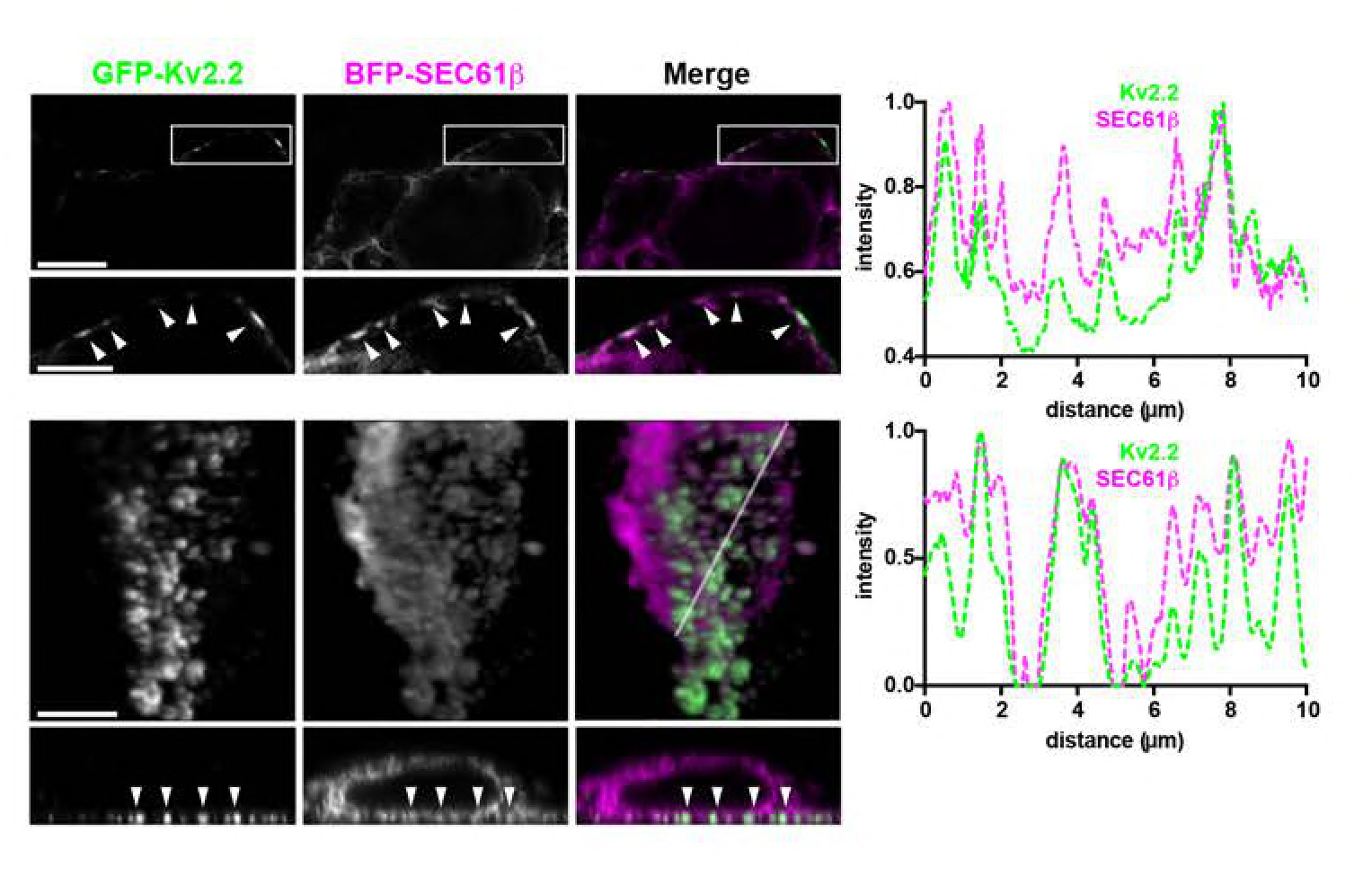
Exogenous Kv2.2 associates with ER-PM junctions in HEK293T cells. Images of fixed HEK293T cells coexpressing GFP-Kv2.2 and BFP-SEC61β. The top two rows show a single optical section taken through the center of the cell. The scale bar in the low magnification panel is 2.5 µm, and for the enlarged panel is 1.25 µm. The bottom rows show a 2D projection of a 3D reconstruction (top row), and a single orthogonal slice through the 3D reconstruction (bottom row). Scale bar in the GFP-Kv2.2 panel of the 3D reconstruction is 2.5 µm, and holds for all panels in bottom two rows. Panels to the right of each set of rows are the corresponding normalized fluorescence intensity values across the individual line scans depicted by the arrows (top) or white line (bottom) in the merged images.

### Kv2.2 expression organizes ER-PM junctions in cultured rat hippocampal neurons and heterologous cells

We next determined the impact of exogenous expression of recombinant Kv2.2 on ER-PM junctions in mammalian neurons and heterologous cells. We used Total Internal Reflection Fluorescence (TIRF) microscopy of living cells to selectively visualize fluorescence signals from ER and PM proteins localized within ≈100 nm of the coverslip (*i.e.*, at ER-PM junctions). In HEK293T cells expressing the fluorescent luminal ER marker DsRed2-ER5 [a general ER marker; (61)], the near-PM ER appeared as a highly ramified system of small reticular tubules and puncta (Figure 3), the latter representing focal structures of cortical ER coincident with the PM or ER-PM junctions (51, 62). Expression of GFP-Kv2.2 led to a reorganization of the DsRed2-ER5-positive cortical ER to form larger foci that colocalized with the clusters of PM-localized Kv2.2 (Figure 3). Cells co-expressing GFP-Kv2.2 exhibited a significant increase in both the size of ER-PM junctions (Figure 3; Figure 3-Table 1) and the percentage of basal cell surface area with associated cortical ER (Figure 3; Figure 3-Table 2). No such changes were seen in cells expressing the Kv channel Kv1.4 (Figure 3; Figure 3-Tables 1, 2). Analysis of colocalization using Pearson’s Correlation Coefficient (PCC) measurements revealed that DsRed2-ER5 was significantly more colocalized with Kv2.2 than it was with Kv1.4 (Figure 3; Figure 3-Table 3). We also found a nearly linear relationship between Kv2.2 cluster size and ER-PM junction size (Figure 4). As previously reported (51), significant increases in ER-PM junction size and ER-associated PM surface area were also observed in cells expressing Kv2.1 (Figure 3). Taken together, these data demonstrate that Kv2.2 can reorganize ER-PM junctions, and that this is a conserved function of Kv2 channels not shared with Kv1.4.

**Figure 3.**
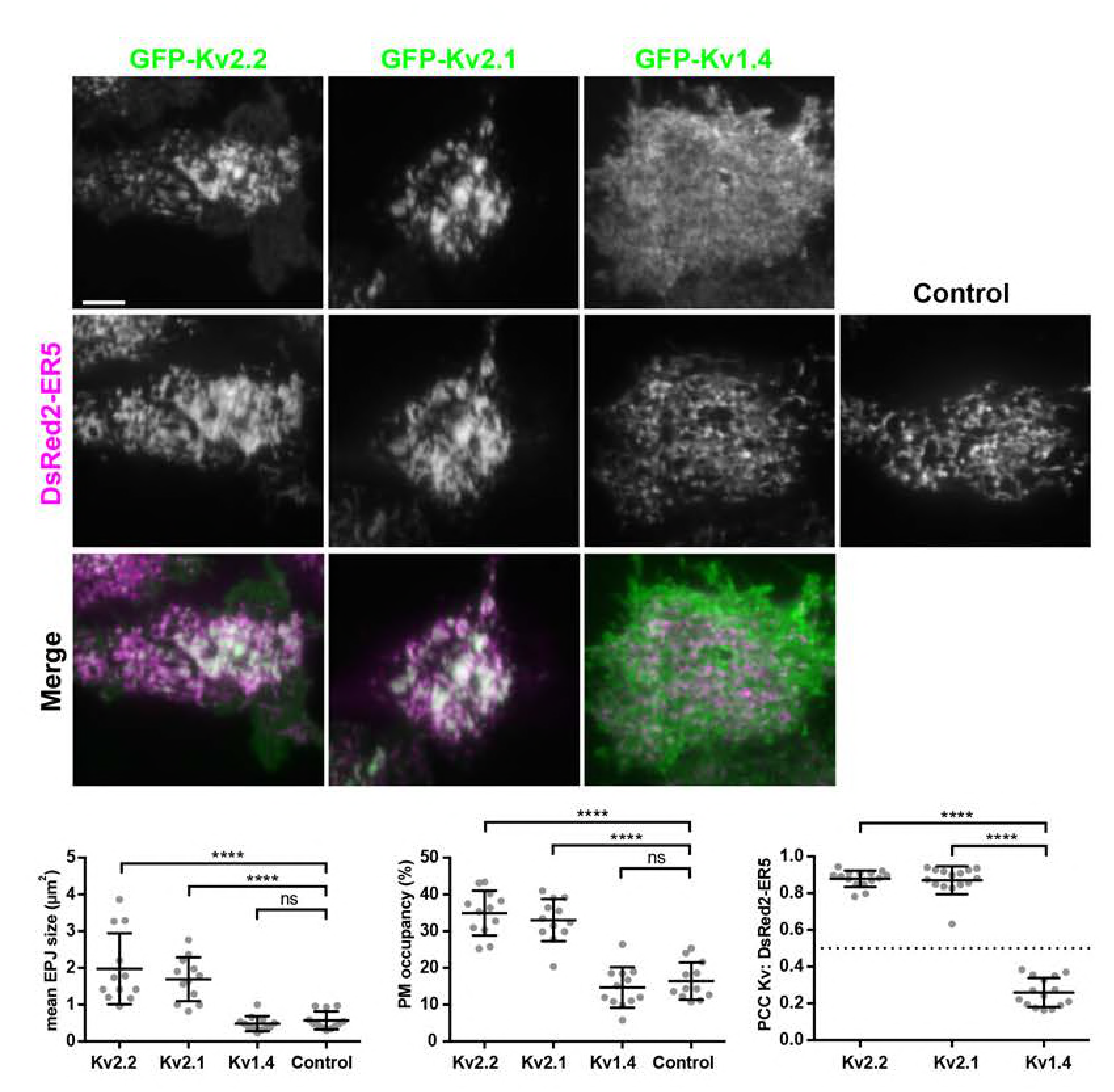
Exogenous Kv2 expression remodels ER-PM junctions in HEK293T cells. TIRF images of live HEK293T cells expressing DsRed2-ER5 either alone, or in conjunction with GFP-Kv2.2, GFP-Kv2.1, or GFP-Kv1.4, as indicated. Scale bar is 5 µm and holds for all panels. Graphs on bottom show population data. Left: Graph of mean ER-PM junction (EPJ) size per cell measured from HEK293T cells coexpressing DsRed2-ER5 and GFP-Kv2.2, GFP-Kv2.1, GFP- Kv1.4, or DsRed2-ER5 alone (control). Middle: Graph of percent of the PM area per cell occupied by cortical ER measured from HEK293T cells coexpressing DsRed2-ER5 and GFP-Kv2.2, GFP- Kv2.1, GFP-Kv1.4, or DsRed2-ER5 alone (control). Right: graph of Pearson’s Correlation Coefficient (PCC) values between DsRed2-ER5 and GFP-Kv2.2, GFP-Kv2.1, or GFP-Kv1.4 measured from HEK293T cells coexpressing DsRed2-ER5 and GFP-Kv constructs. The dashed line denotes a PCC value of 0.5. Bars on all graphs are mean ± SD. See Figure 3-Tables 1-3 for values and statistical analyses.

**Figure 4.**
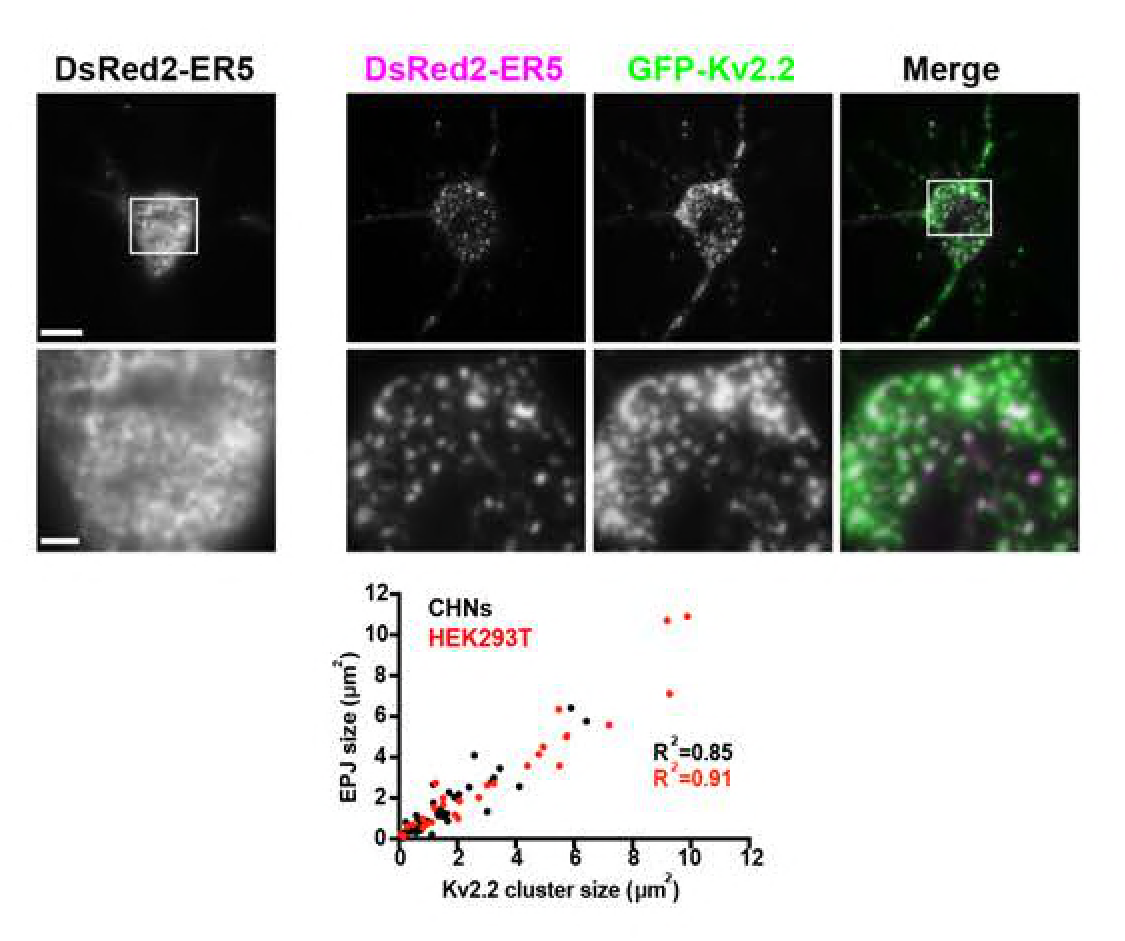
Exogenous Kv2.2 expression remodels ER-PM junctions in cultured neurons. TIRF image of a live CHN (DIV7) expressing DsRed2-ER5 alone (left panel and inset shown below) or coexpressing DsRed2-ER5 and GFP-Kv2.2 (right panels and insets shown below). Scale bar in DsRed2-ER5 panel is 10 µm and holds for all panels in that row. Scale bar in DsRed2-ER5 magnified inset panel is 2.5 µm and holds for all panels in that row. Scatter plot shows sizes of Kv2.2 clusters and associated ER-PM junctions (EPJs, as reported by DsRed2-ER5 in TIRF) in CHNs (black points) and in HEK293T cells (red points). n = 3 cells each.

We next expressed DsRed2-ER5 alone or coexpressed DsRed2-ER5 with GFP-Kv2.2 in CHNs. TIRF imaging experiments revealed that GFP-Kv2.2 expression remodeled neuronal ER-PM junctions (Figure 4). Similar to HEK293T cells, we found a nearly linear relationship between Kv2.2 cluster size and ER-PM junction size in CHNs (Figure 4). These results demonstrate that expression of Kv2.2 in both HEK293T cells and CHNs is sufficient to remodel ER-PM junctions.

### Kv2.2 channels associated with ER-PM junctions are on the cell surface

Given the extensive colocalization of Kv2.2 and these ER markers at ER-PM junctions, we further addressed whether the Kv2.2 present at these sites was in the PM. We employed live cell labeling with the Kv2-specific tarantula toxin Guangxitoxin-1E (63) conjugated to DyLight633 [GxTX-633; (64)] to label cell surface Kv2.2. We first validated this approach by coexpressing BFP-SEC61β with SEP-Kv2.1, a construct of Kv2.1 tagged with cytoplasmic mCherry and an extracellular pHluorin as a reporter of cell surface Kv2.1 (65). We observed extensive colocalization of GxTX- 633 and pHluorin signals (Figure 5-figure supplement 1), showing that GxTX-633 is a reliable reporter for cell surface Kv2 channels. No detectable GxTX-633 labeling was observed in control HEK293T cells, or those expressing DsRed2-ER5 alone (data not shown). GxTX-633 labeling of cells coexpressing GFP-Kv2.2 and DsRed2-ER5 showed a high degree of colocalization of all three signals (Figure 5). As expected, PCC measurements (Figure 5) were slightly but significantly higher for direct labeling of Kv2.2 with GxTX than for indirect labeling of ER-PM junctions with GxTX (Figure 5-Table 1). Similar results were obtained for GxTX labeling of cells coexpressing SEP-Kv2.1 and Sec61β (Figure 5-figure supplement 1, Figure 5-Table 1). Taken together, these data demonstrate that the Kv2 clusters associated with ER-PM junctions are on the cell surface.

**Figure 5.**
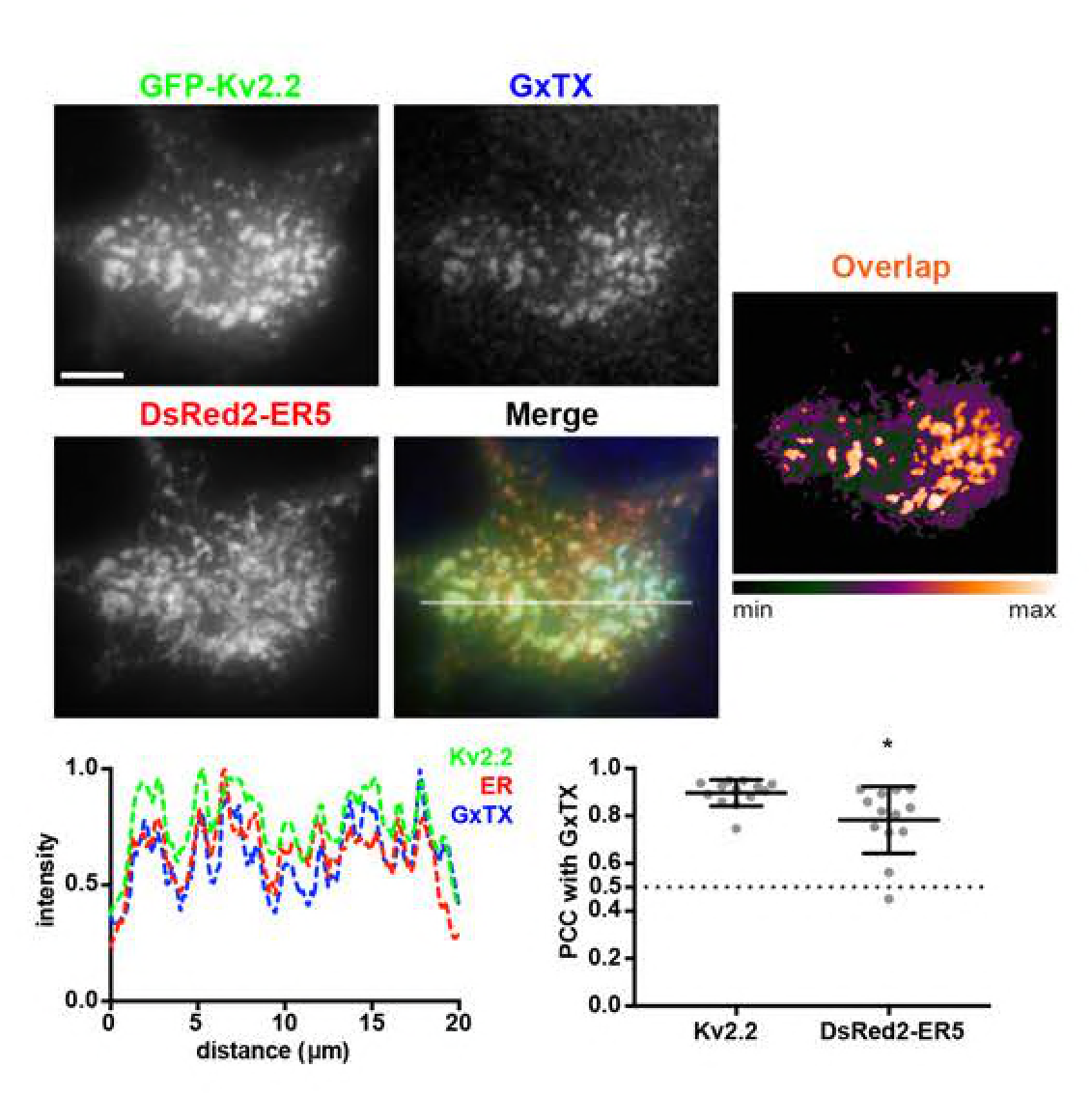
ER-PM junction-localized Kv2.2 channels are expressed on the cell surface. TIRF images of a live HEK293T cell expressing GFP-Kv2.2 and DsRed2-ER5, and surface labeled for Kv2 channels with GxTX-633. Heat map shows overlap of GFP-Kv2.2 and GxTX-633 pixels. Scale bar is 5 µm. Bottom left panel shows the fluorescence intensity values across the individual line scan depicted by the white line in the merged image. Graph on bottom right shows the PCC values between pairs of indicated signals as measured from live HEK293T cells surface labeled with GxTX-633 and coexpressing GFP-tagged Kv2.2 channels and DsRed2-ER5. Bars are mean ± SD. See Figure 5-Table 1 for values and statistical analyses.

### Kv2.2-containing ER-PM junctions are present at sites depleted in components of the cortical actin cytoskeleton

Kv2.2 is expressed in large clusters in brain neurons, including on the axon initial segment or AIS (18, 66), a subcellular compartment highly enriched for components of the actin cortical cytoskeleton including a specialized complex of spectrins and ankyrins (67). We immunolabeled brain sections for Kv2.2 and ankyrinG (ankG), which is highly expressed at the AIS. We found that in neocortical layer 5 pyramidal neurons, in addition to the somatodendritic labeling shown for Kv2.2 in Figure 1, Kv2.2 was also present in robust clusters on the AIS (Figure 6). The AIS clusters of Kv2.2 in these neurons were found at sites deficient in ankG (Figure 6). These ankG-deficient sites represent locations at which the ER present in the AIS, termed the cisternal organelle, comes into close apposition to the PM (68-70).

**Figure 6.**
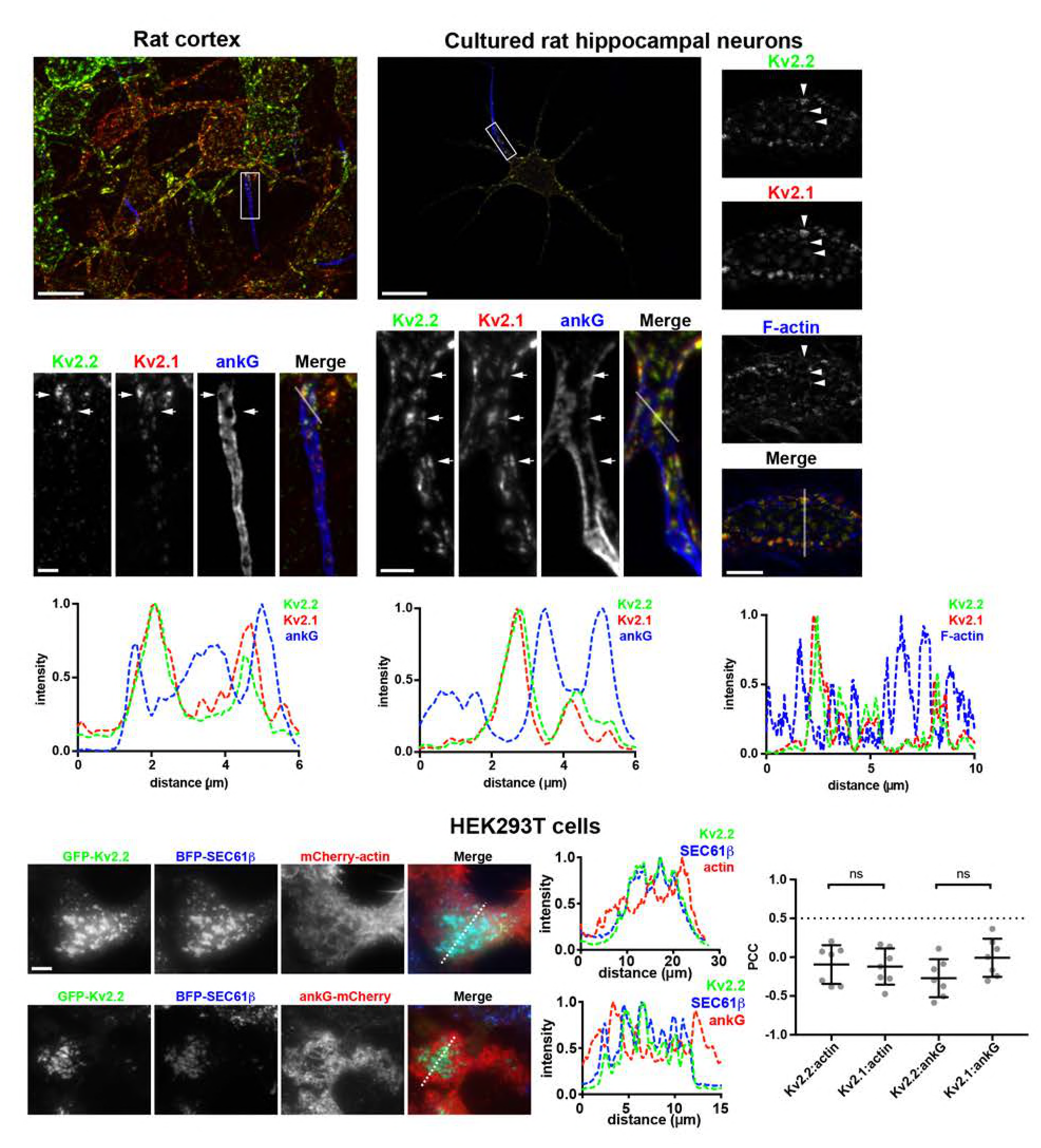
Kv2-mediated ER-PM junctions are located at sites depleted in components of the cortical actin cytoskeleton. Top left panels. Brain sections immunolabeled for Kv2.2, Kv2.1, and ankG. Scale bar for large image is 20 µm, and for Kv2.2 inset is 3 µm and holds for all inset panels. Middle panels. Projected z-stack of optical sections taken from a CHN immunolabeled for Kv2.2, Kv2.1, and ankG. Scale bar for large image is 20 µm, and for Kv2.2 inset is 3 µm and holds for all inset panels. Right panels. Single optical section taken from a CHN immunolabeled for Kv2.2, Kv2.1, and labeled for F-actin with phalloidin. Scale bar for merged panel is 10 µm holds for all panels in set. Panels below each set of images show the corresponding normalized fluorescence intensity values across the line scans indicated in the merged images in that column. Lower panels. TIRF images of live HEK293T cells coexpressing GFP-Kv2.2 and BFP-SEC61β in conjunction with mCherry-actin (top row) or ankG-mCherry (bottom row). Scale bar for GFP-Kv2.2 panel in top row is 5 µm and holds for all panels in set. Panels to the right of these rows show the corresponding normalized fluorescence intensity values across the line scan depicted by the white line in the merged images. Graph shows PCC values from cells coexpressing either GFP-Kv2.2 or GFP-Kv2.1 and mCherry-actin or ankG-mCherry. Bars on all graphs are mean ± SD. See Figure 6-Table 1 for values and statistical analyses.

We next immunolabeled for endogenous Kv2.2 and ankG in CHNs and found a similar relationship between the sites of Kv2.2 clustering on the AIS and regions deficient in both ankG and filamentous actin, the latter labeled with fluorescent phalloidin (Figure 6). This is apparent in line scan analyses, which revealed that the intensity profiles of the Kv2 immunolabeling and actin labeling were often negatively correlated (Figure 6). To determine whether this spatial relationship is also present in non-neuronal cells, we performed TIRF imaging on live HEK293T cells coexpressing GFP-Kv2.2, BFP-SEC61β and mCherry-tagged actin. We found that GFP-Kv2.2 clusters and associated ER-PM junctions displayed a negatively correlated distribution with respect to cortical mCherry-actin (Figure 6). Kv2.1 clusters and associated ER-PM junctions exhibited a similar negative relationship to the cortical actin cytoskeleton (Figure 6-figure supplement 1). The negative values of PCC measurements between either of the Kv2 channels and mCherry-actin confirmed this (Figure 6; Figure 6-Table 1). We additionally coexpressed ankG-mCherry with BFP-SEC61β and either Kv2.2, or Kv2.1, and again found a negatively correlated distribution of the Kv2 channel clusters and associated ER-PM junctions with this actin-associated protein (Figure 6, Figure 6-figure supplement 1). PCC measurements show that neither Kv2.2 nor Kv2.1 colocalized with cortical ankG-mCherry (Figure 6, Figure 6-Table 1).

### The actin cytoskeleton regulates the organization of Kv2.2 clusters and associated ER-PM junctions

Given the distinct spatial relationship between Kv2.2-associated ER-PM junctions and the cortical actin cytoskeleton, we next determined the impact of disrupting the organization of the actin cytoskeleton on characteristics of Kv2.2-mediated ER-PM junctions. We treated cells expressing Kv2.2 with Latrunculin A (LatA) which disrupts the organization of filamentous actin (71). We found LatA treatment led to a reorganization of Kv2.2 clusters and the associated ER-PM junctions (Figure 7), resulting in a significant increase in the size of both Kv2.2 clusters and ER-PM junctions (Figure 7; Figure 7-Table 1), the latter reported by the DsRed2-ER5 signal coincident with the PM. The total number of ER-PM junctions in Kv2.2-expressing cells was significantly reduced in response to LatA treatment (Figure 7; Figure 7-Table 2). Similar results were obtained upon LatA treatment of cells coexpressing GFP-Kv2.1 and DsRed2-ER5 (Figure 7-figure supplement 1; Figure 7-Table 1), as suggested in a previous study (51). These changes were not observed in untreated cells over the course of 15 minutes (data not shown). While LatA treatment significantly altered the spatial characteristics of the Kv2.2 clusters and the Kv2.2- associated ER-PM junctions, the extent of colocalization between GFP-Kv2.2 and DsRed2-ER5 was not significantly altered upon LatA treatment (Figure 7; Figure 7-Table 3). Similar results were obtained for Kv2.1 (Figure 7-figure supplement 1; Figure 7- Tables 2-3). These results show that while LatA induced an apparent fusion of Kv2 clusters and associated ER-PM junctions resulting in fewer, larger structures, it did not affect their association *per se*. These results also suggest that the distinct and mutually exclusive localization of Kv2.2 clusters and components of the cortical actin cytoskeleton seen in brain neurons likely participates in the organization and maintenance of Kv2 clusters and the associated ER-PM junctions.

**Figure 7.**
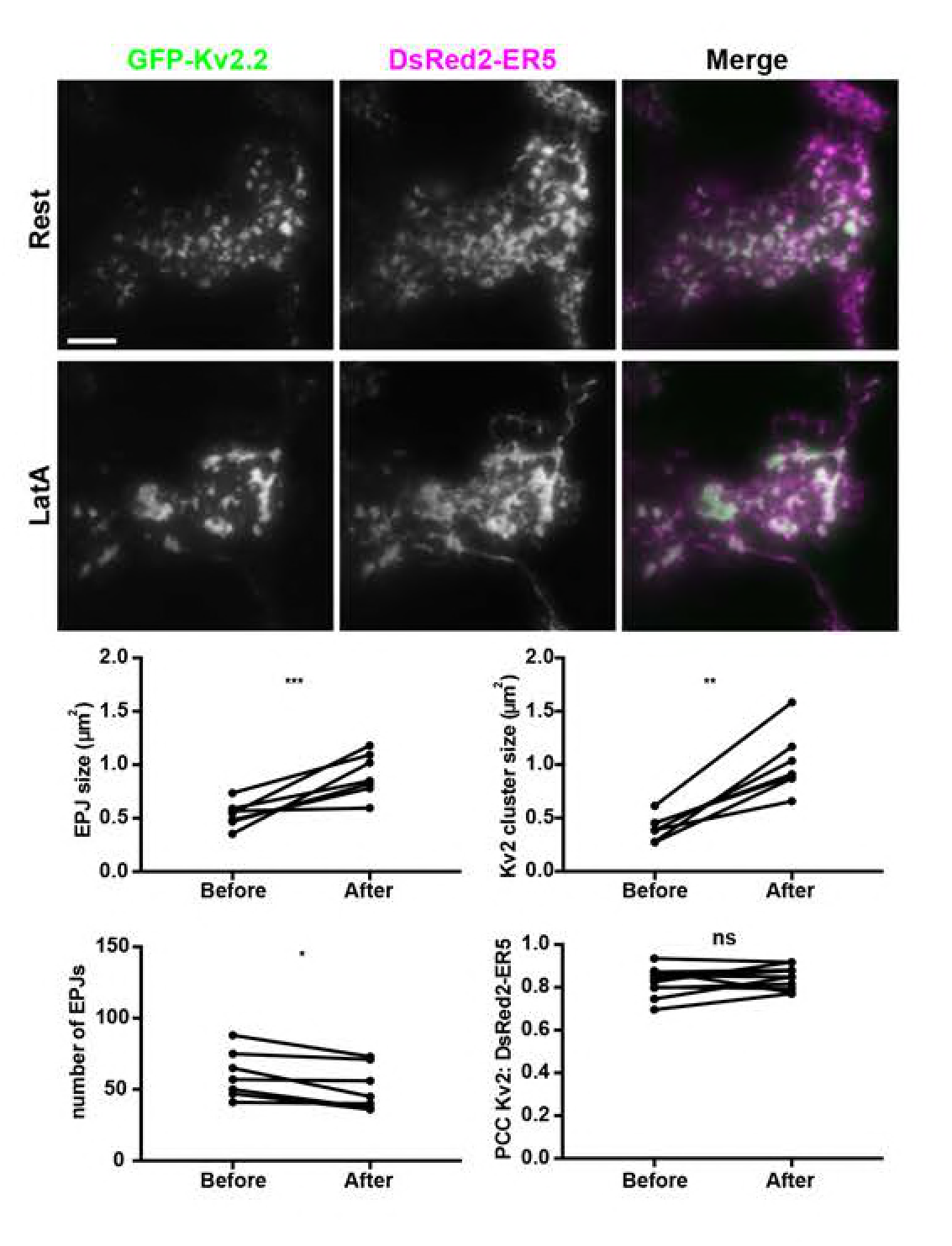
Disrupting the actin cytoskeleton impacts spatial organization of Kv2.2-mediated ER-PM junctions. TIRF images of a live HEK293T cell coexpressing GFP-Kv2.2 and DsRed2-ER5, prior to, and 15 min after, Latrunculin A (LatA) treatment. Scale bar in GFP-Kv2.2 Rest panel is 5 µm and holds for all panels. Graphs below show values measured from cells before and after a 15-minute treatment with 10 µM LatA. Top left graph. Mean ER-PM junction (EPJ) size per cell. Top right graph: Mean Kv2.2 cluster size per cell. Bottom left graph. Number of ER-PM junctions per cell. Bottom right graph. PCC values between Kv2.2 and DsRed2-ER5. Bars on all graphs are mean ± SD. See Figure 7-Tables 1-3 for values and statistical analyses.

### Kv2.2-containing ER-PM junctions associate with ER-PM junctions formed by the known classes of ER-PM tethers

We next determined the relationship of Kv2.2 clusters and associated ER-PM junctions with the three other families of mammalian ER-localized ER-PM tethers. We coexpressed FP-tagged Kv2.2 and individual members of the E-Syt, JP and STIM families in HEK293T cells. In cells coexpressing the STIMs, we also induced Ca^2+^ store depletion *via* treatment with 2 μM thapsigargin treatment for five minutes. In all cases, we observed a high degree of colocalization between clusters of Kv2.2 and these ER-PM junction tethers (Figure 8, Figure 8-figure supplements 1, 2), as demonstrated by high PCC and Mander’s overlap coefficient (MOC) values (Figure 8-Table 1). In cells coexpressing STM1, Kv2.2 and Orai1, store depletion resulted in not only a significant increase in colocalization of STIM1 and Orai1, but also of Orai1 and Kv2.2 (Figure 8-figure supplement 3; Figure 8-Table 4). The store depletion-induced increase in colocalization of Kv2.2 and Orai1 also occurred in the absence of STIM1 coexpression (Figure 8- figure supplement 3; Figure 8-Table 4), presumably due to endogenous STIM expression in HEK293T cells (72-75). Together, these results show that Kv2.2 clusters are associated with ER-PM junctions formed by the three established families of ER-PM junction tethers. Interestingly, the PCC values were significantly lower than the corresponding MOC values obtained from the same cells (Figure 8; Figure 8-Table 1), suggesting that despite the extensive overlap in signal between Kv2.2 clusters and these established classes of ER-PM junctions, there are distinctions in their fine spatial organization relative to one another. Kv2.1 also exhibited a high degree of colocalization with these diverse ER-PM junction tethers (Figure 8-figure supplement 1, Figure 8-Table 3).

**Figure 8.**
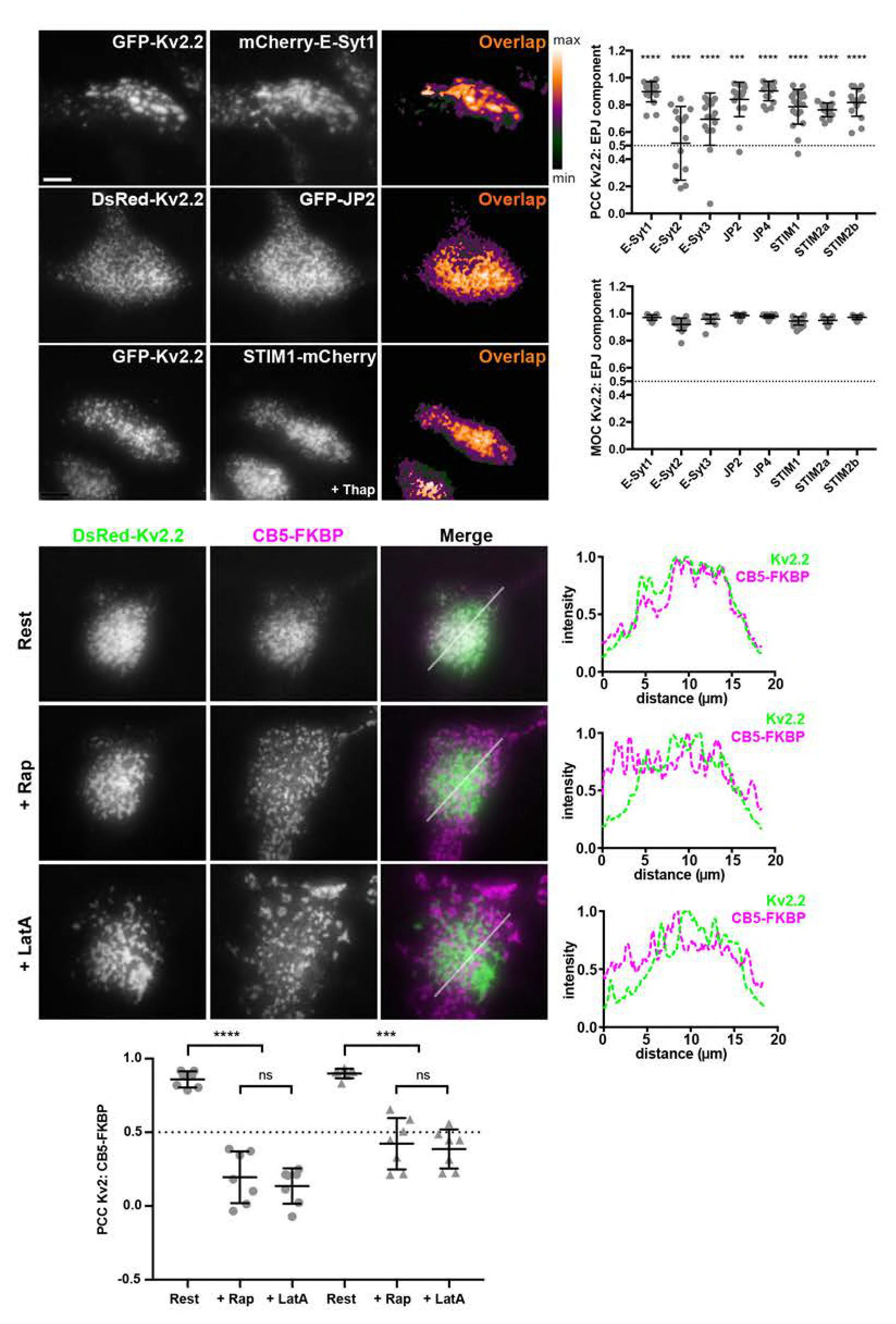
Kv2-containing ER-PM junctions colocalize with multiple components of mammalian ER-PM junctions. Upper panels. TIRF images of live HEK293T cells coexpressing GFP or DsRed-Kv2.2 and representative members of the E-Syt, JP and STIM families of ER-localized PM tethers. Scale bar in top left GFP-Kv2.2 panel is 5 µm and holds for all panels in figure. Heat maps show pixel overlap of GFP-Kv2.2 and ER-PM tether signals. The STIM1 sample was treated with 2 µM thapsigargin for 5 minutes prior to imaging. Graphs to right show PCC and MOC values of Kv2.2 and ER-PM tether signals. Bars are mean ± SD. See Figure 8-Table 1 for values and statistical analyses. Lower panels. TIRF images of live HEK293T cells coexpressing DsRed-Kv2.2, CFP- CB5-FKBP, and Lyn11-FRB. Top row. Prior to rapamycin treatment (rest). Middle row. Same cell immediately following 5 µM rapamycin treatment (+Rap). Bottom row. Same cell after subsequent 15-minute treatment with 10 µM LatA (+LatA). Panels to the right of each row are the corresponding normalized fluorescence intensity values across the individual line scans depicted by the white line in the merged images. Bottom graph shows PCC values between DsRed-Kv2.2 and CFP-CB5-FKBP signals. Bars are mean ± SD. See Figure 8-Table 2 for values and statistical analyses.

We further examined the relationship of Kv2-mediated ER-PM junctions to those previously characterized by acutely triggering ER-PM junction formation using a rapamycin-inducible system (76) employing ER-localized CB5-FKBP-CFP and PM-localized Lyn11-FRB (CB5/Lyn11). TIRF imaging reveals that acute treatment of HEK293T cells coexpressing CB5/Lyn11 with 5 μM rapamycin yields robust recruitment of ER to the cell cortex (Figure 8-figure supplement 3). HEK293T cells coexpressing Kv2.2 and CB5/Lyn11 prior to rapamycin addition exhibited CB5-FKBP-CFP fluorescence similar to other ER reporters (*e.g.*, BFP-SEC61β, DsRed2-ER5) in being throughout the ER, and also colocalized with clustered Kv2.2 at ER-PM junctions, the latter yielding a high degree of colocalization in TIRF imaging (Figure 8; Figure 8- Table 2). Surprisingly, unlike the other classes of ER-PM junctions, the rapamycin-induced CB5/Lyn11 ER-PM junctions were largely distinct and nonoverlapping from those associated with the Kv2.2 clusters (Figure 8), as shown by the significant decrease in PCC values upon rapamycin treatment (Figure 8; Figure 8-Table 2). Subsequent LatA treatment impacted the spatial organization of both the Kv2.2- and CB5/Lyn11-mediated ER-PM junctions (Figure 8). However, they remained spatially segregated such that there were no significant LatA-induced changes in PCC values between Kv2.2- and CB5 (Figure 8; Figure 8-Table 2). Similar results were obtained for Kv2.1 (Figure 8-figure supplement 4; Figure 8-Table 2). These results taken together demonstrate that despite the extensive colocalization observed between Kv2-associated ER-PM junctions and those formed by known ER-PM junction tethers, ER-PM junctions distinct from those mediated by Kv2 clustering can exist simultaneously in mammalian cells. Moreover, while the actin cytoskeleton plays a role in defining the spatial boundaries of both Kv2.2- and CB5/Lyn11-mediated ER-PM junctions, disrupting the actin cytoskeleton is not sufficient to homogenize these distinct membrane contact sites.

### Reorganization of cortical ER is a nonconducting function of Kv2.2

We next addressed whether the Kv2.2-mediated remodeling of ER-PM junctions is dependent on K^+^ flux through the channels. We generated a point mutation (P412W) in the S6 transmembrane helix of Kv2.2 that is at the same relative position as a point mutation (P404W) that eliminates conductance through Kv2.1 channels heterologously expressed in *Xenopus* oocytes (77). We first expressed GFP-Kv2.2 P412W in HEK293T cells and evaluated conductance relative to wild-type GFP-Kv2.2 using voltage-clamp electrophysiology. HEK293T cells expressing GFP-Kv2 channels or GFP alone as a control were whole-cell patch clamped and held at a resting membrane potential of −80 mV. In response to positive voltage steps, delayed rectifier outward currents emerged from cells expressing GFP-Kv2.2, but not from cells expressing either GFP-Kv2.2 P412W or GFP (Figure 9, Figure 9-Table 1). As expected from previous analyses in oocytes, GFP-Kv2.1 P404W was nonconducting when expressed in HEK293T cells (Figure 9-figure supplement 1, Figure 9-Table 1).

**Figure 9.**
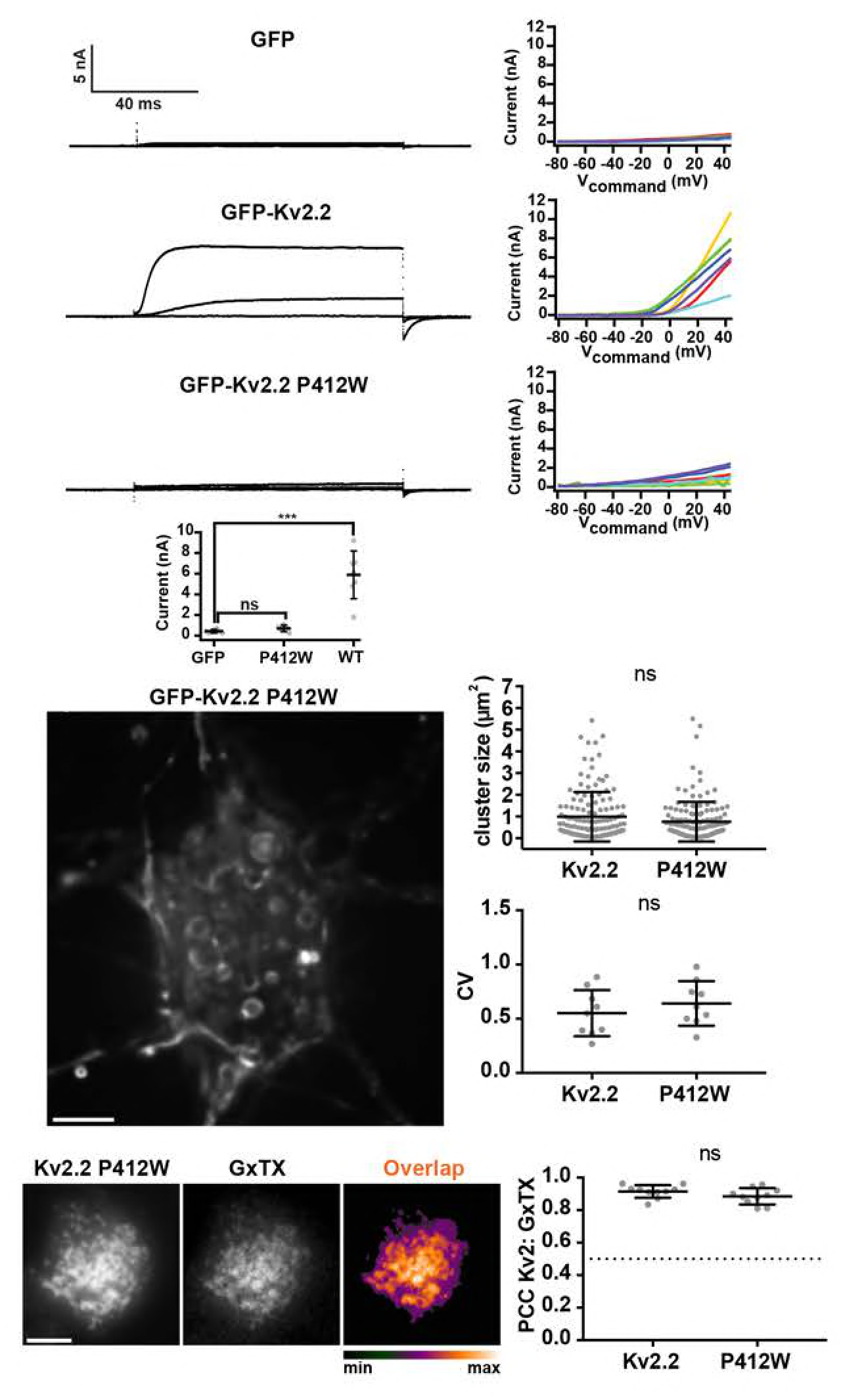
Mutations that eliminate K^+^ conductance do not impact Kv2.2 channel clustering. Top panels show exemplar whole-cell voltage clamp recordings (left) and corresponding graphs of current levels versus command voltage (right) of HEK293T cells expressing GFP (control), GFP-Kv2.2, or GFP-Kv2.2 P412W. Recordings shown are representative responses to 100 ms steps from −100 mV to −40, 0 and +40 mV. Note the lack of outward currents in control and GFP- Kv2.2 P412W recordings. Summary graph shows whole cell current at +40 mV. See Figure 9- Table 1 for values and statistical analyses. Middle panel shows a deconvolved widefield image of a live DIV 7-10 CHN expressing GFP-Kv2.2 P412W. Scale bar is 5 µm. Graphs to the right are measurements of mean cluster size per cell and CV values measured from CHNs expressing GFP-Kv2.2 or GFP-Kv2.2 P412W. Bars are mean ± SD. See Figure 9-Tables 2-3 for values and statistical analyses. Bottom panels show TIRF images of live HEK293T cells expressing GFP- Kv2.2 P412W and surface labeled with GxTX-633. Scale bar in the Kv2.2 P412W panel is 5 µm and hold for all panels in row. The graph to the right shows comparisons of PCC measurements of Kv2 and GxTX fluorescence from HEK293T cells expressing GFP-Kv2.2 and/or GFP-Kv2.2 P412W. Dashed line denotes a PCC value of 0.5. Bars are mean ± SD. See Figure 9-Table 4 for values and statistical analyses.

We next expressed GFP-Kv2.2 P412W in CHNs and found that it was localized in clusters indistinguishable from GFP-Kv2.2 (Figure 9). The size of GFP-Kv2.2 P412W clusters was not significantly different than those of GFP-Kv2.2 (Figure 9, Figure 9-Table 2). We used Coefficient of Variation (CV) of pixel intensity as a quantitative measure of nonuniformity imparted by clustering (39, 41, 47, 65). We found that CV values for GFP-Kv2.2 P412W expressed in CHNs were not significantly different than those for GFP-Kv2.2 (Figure 9; Figure 9-Table 3). We also found a lack of any significant differences in clustering of conducting GFP-Kv2.1 and nonconducting GFP-Kv2.1 P404W (Figure 9-figure supplement 1, Figure 9-Tables 2-3).

We next surface labeled live HEK293T cells with GxTX-633 and found no significant differences in colocalization between GxTX-633 and GFP-Kv2.2 *versus* GFP-Kv2.2 P412W (Figure 9; Figure 9-Table 4). A similar lack of significant differences was seen for GxTX labeling of GFP-Kv2.1 versus nonconducting GFP-Kv2.1 P404W (Figure 9-figure supplement 1, Figure 9- Table 4). These data taken together demonstrate that these Kv2 mutants lack ionic conductance but exhibits cell surface expression and clustering indistinguishable from their wild-type counterparts.

We next addressed whether the clustered but nonconducting GFP-Kv2.2 P412W mutant retained its ability to recruit/stabilize cortical ER at ER-PM junctions. Live cell TIRF imaging showed that GFP-Kv2.2 P412W reorganized the DsRed2-ER5-labeled cortical ER into ER-PM junctions (Figure 10). We found no significant difference between cells expressing GFP-Kv2.2 P412W versus GFP-Kv2.2 in either the size of ER-PM junctions (Figure 10; Figure 10-Table 1), or the surface area of the PM occupied by the cortical ER (Figure 10; Figure 10-Table 2). The extent of colocalization of DsRed2-ER5 with GFP-Kv2.2 P412W was also not significantly different than for GFP-Kv2.2 (Figure 10; Figure 10-Table 3). We next evaluated the lateral mobility of DsRed2-ER5-labeled cortical ER as an additional measure of its recruitment into ER-PM junctions (51, 78). The mobility of PM-associated ER was significantly reduced in Kv2.2- expressing cells compared to control cells expressing DsRed2-ER5 alone (Figure 10-figure supplement 2; Figure 10-Table 6). Cortical ER mobility was not significantly different in cells expressing the nonconducting Kv2.2 P412W mutant versus those expressing WT Kv2.2 (Figure 10-figure supplement 2; Figure 10-Table 6). These parameters of cortical ER recruitment/stabilization were also not significantly different between WT Kv2.1 and the nonconducting Kv2.1 P404W mutant (Figure 10-figure supplements 1, 2, Figure 10-Tables1-4). These data taken together demonstrate that the function of Kv2 channels to localize to and organize ER-PM junctions is independent of their canonical ion conducting function and is instead a distinct nonconducting function.

**Figure 10.**
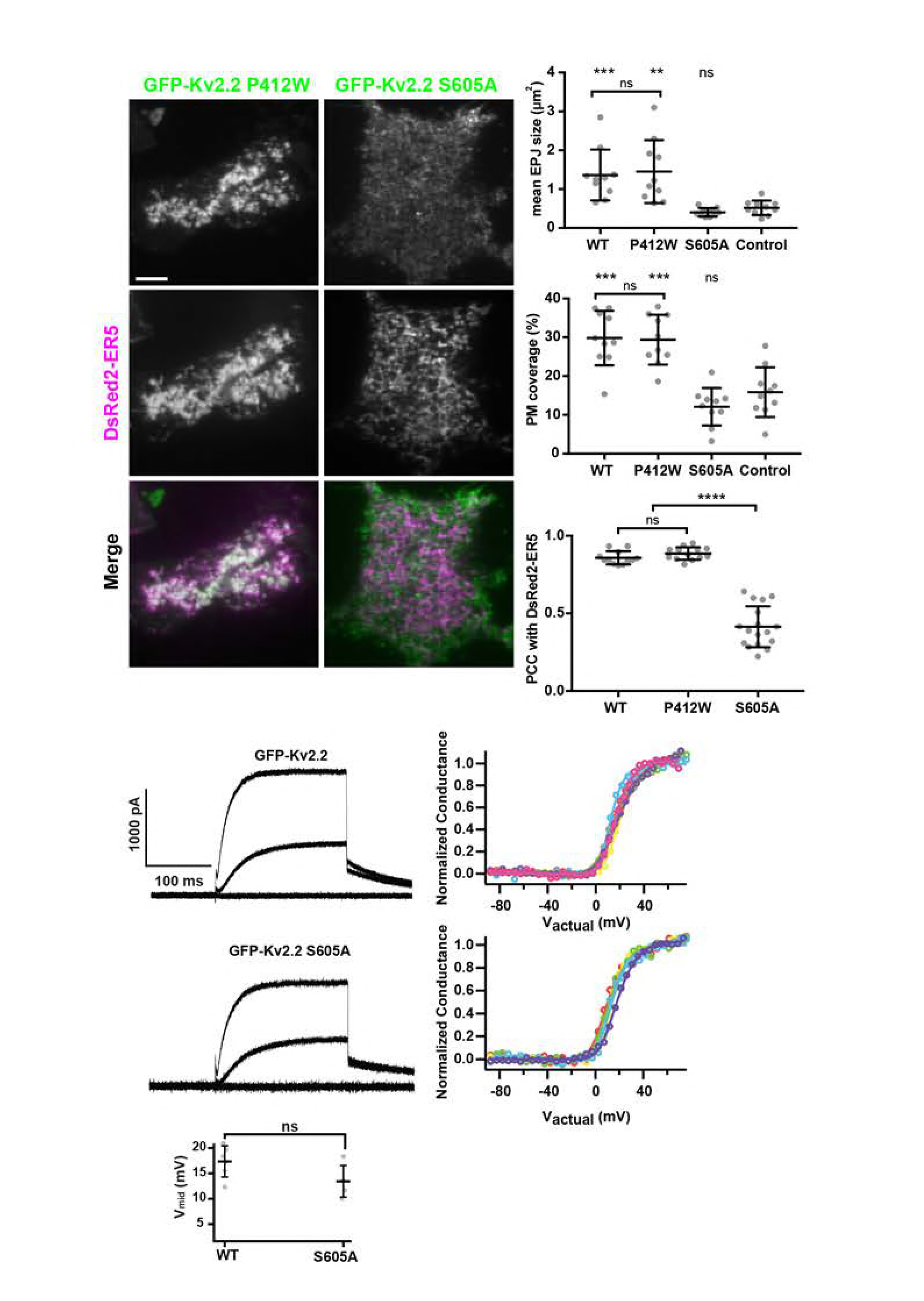
Separation of function point mutations show that clustering, but not conduction, is necessary for Kv2.2-mediated remodeling of ER-PM junctions. Left panels show TIRF images of live HEK293T cells expressing GFP-tagged Kv2.2 mutants (nonconducting P412W and nonclustering S605A) and DsRed2-ER5. Scale bar is 5 µm and holds for all panels. Graphs to right show comparisons from cells expressing wild-type and mutant Kv2.2 isoforms (P412W or S605A); control refers to cells expressing DsRed2-ER5 alone. Top right graph. Mean ER-PM junction (EPJ) size per cell. Middle right graph. Percent PM per cell occupied by cortical ER. Lower right graph. PCC values between DsRed2-ER5 and wild-type (WT) and mutant Kv2.2 isoforms. Bars on all graphs are mean ± SD. See Figure 10-Tables 1-3 for values and statistical analyses. Bottom panels show exemplar whole-cell voltage clamp recordings (left) and graphs of the corresponding normalized conductance-voltage relationship from HEK293T cells expressing GFP-Kv2.2, or GFP-Kv2.2 S605A (right). Different colors represent data from distinct cells. Recordings shown are representative responses to 200 ms steps from −100 mV to −40, 0 and +40 mV. Bottom graph shows V_mid_ values. Note the lack of effect of the declustering point mutation on the properties of the whole cell currents. See Figure 10-Tables 4-5 for values and statistical analyses.

We next determined whether Kv2.2 clustering is necessary for remodeling of ER-PM junctions. We used a point mutant in the cytoplasmic C-terminus of Kv2.2 (S605A) that abolishes its clustering (39). Based on analyses of C-terminal truncation mutants in Kv2.1 [*e.g.*, (35, 79)], we expected that this point mutant would not impact the ability of Kv2.2 to conduct K^+^. To verify this, we used whole cell patch clamp recordings to compare currents from wild-type and nonclustered Kv2.2 channels in voltage clamped cells. We found that expression of GFP-Kv2.2 S605A in HEK293T cells resulted in expression of voltage-activated outward currents (Figure 10). The conductance-voltage relationships of cells expressing GFP-Kv2.2 versus GFP-Kv2.2 S605A were not significantly different (Figure 10; Figure 10-Table 4), nor were those from GFP-Kv2.1 versus GFP-Kv2.1 S586A (Figure 10-figure supplement 1; Figure 10-Tables 4). The K^+^ current density was also not significantly altered by the point mutations that disrupt Kv2.2 clustering and association with ER-PM junctions. The whole cell K^+^ current density from cells expressing GFP- Kv2.2 *versus* GFP-Kv2.2 S605A were not significantly different (Figure 10; Figure 10-Table 5), nor were those from cells expressing GFP-Kv2.1 versus GFP-Kv2.1 S586A (Figure 10-figure supplement 1; Figure 10-Table 5). Thus, these measurements of current density and the conductance-voltage relationship supports that Kv2 channels with these cytoplasmic point mutations that disrupt clustering do not affect the density of conducting channels on the cell surface or their gating.

Finally, we determined the function of the nonclustering but conducting Kv2.2 S605A point mutant in organizing ER-PM junctions. TIRF imaging revealed a diffuse localization of GFP-Kv2.2 S605A (Figure 10). The ER-PM junction size (Figure 10; Figure 10-Table 1) and percentage of PM surface area occupied by cortical ER (Figure 10; Figure 10-Table 2) were not significantly different between cells coexpressing GFP-Kv2.2 S605A and cells expressing DsRed2-ER5 alone. This nonclustered GFP-Kv2.2 S605A mutant also had a significantly reduced colocalization with coexpressed DsRed2-ER5 relative to GFP-Kv2.2 (Figure 10; Figure 10-Table 3). We obtained similar results for Kv2.1 in that the ability to organize ER-PM junctions was significantly reduced in the nonclustering but conducting GFP-Kv2.1 S586A point mutant (Figure 10-figure supplement 1, Figure 10-Tables 1-3). Taken together, these results using this set of separation-of-function point mutants demonstrate that Kv2 channel clustering, but not conduction, is necessary for the unique ability of PM Kv2 channels to localize to and organize ER-PM junctions, and that the functions of Kv2 channels in conducting ions and organizing ER-PM junctions are separable and distinct.

### Kv2.2- and Kv2.1-mediated ER-PM junctions exhibit distinct cell cycle-dependent regulation in COS-1 cells

Kv2.1 exhibits conditional phosphorylation-dependent clustering that can be regulated by neuronal activity and other stimuli (52-54, 80-82). In certain mammalian cell lines such as COS- 1 cells, Kv2.1 exhibits reversible cell-cycle dependent clustering and recruitment/stabilization of ER-PM junctions, presumably due to increased phosphorylation of Kv2.1 observed at the onset of M-phase (47). To determine whether Kv2.2 exhibited similar cell cycle-dependent regulation, we expressed mCherry-SEC61β with untagged Kv2.2 or Kv2.1 and performed imaging after immunolabeling for the expressed Kv2 channel and Hoechst 33258 staining of chromatin to define cell cycle stage (47). As previously reported, Kv2.1 has an overall diffuse localization in interphase cells and prominent clustering in M-phase cells (Figure 11; Figure 11-figure supplement 1). In contrast, Kv2.2 clusters were present in both interphase and M-phase cells (Figure 11; Figure 11- figure supplement 1), such that in interphase cells Kv2.2 exhibited significantly higher CV values compared to Kv2.1 (Figure 11; Figure 11-Table 1). In contrast, we found no significant difference between CV values for Kv2.2 versus Kv2.1 in M-phase cells (Figure 11; Figure 11-Table 1).

**Figure 11.**
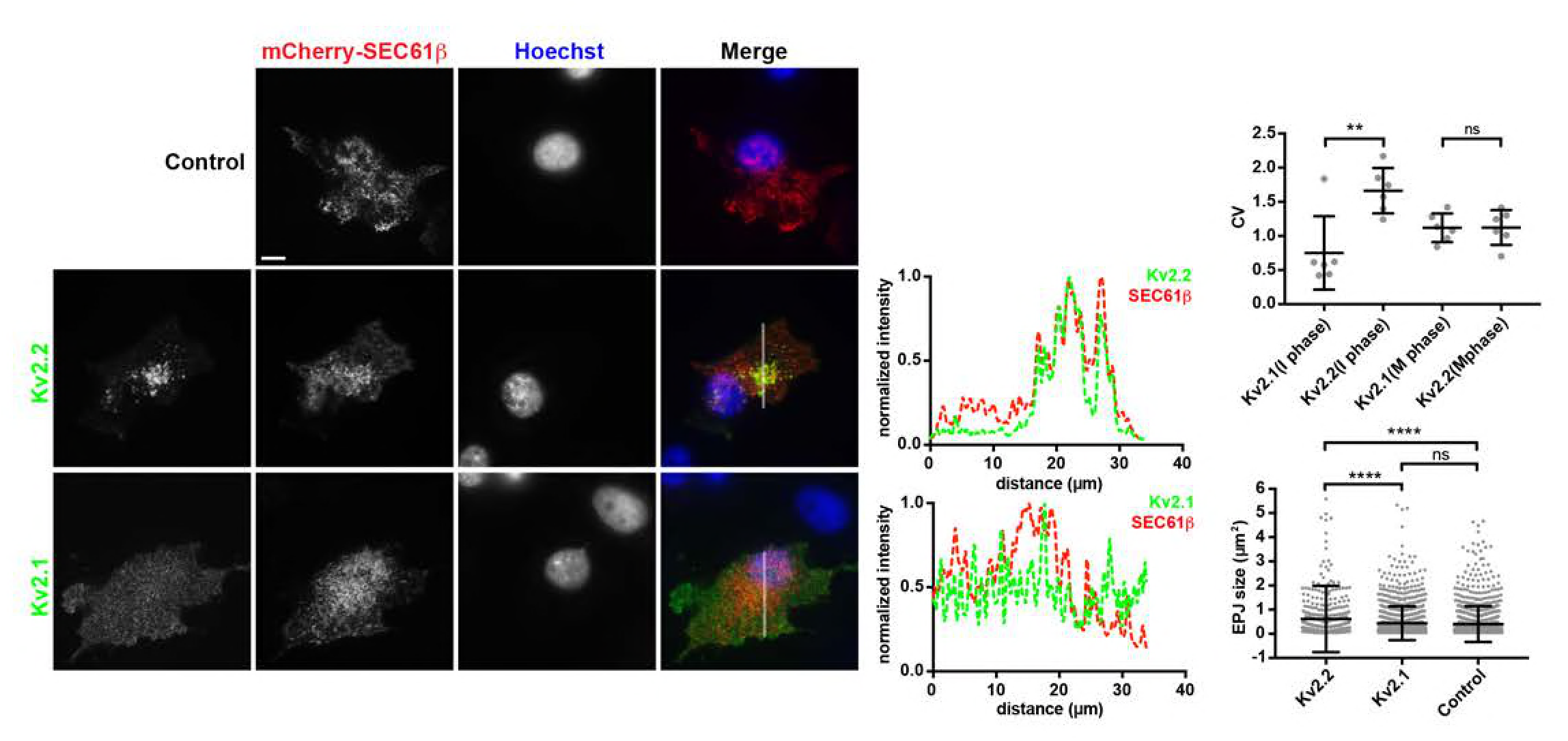
Kv2.2 but not Kv2.1 can organize ER-PM junctions in interphase COS-1 cells. TIRF images of fixed interphase COS-1 cells stained with Hoechst 33258 and expressing mCherry-SEC61β alone, or coexpressing Kv2.2 (immunolabeled with mAb N372B/60) or Kv2.1 (immunolabeling with mAb K89/34). Scale bar is 10 µm and is for all panels. Panels to the right are the normalized fluorescence intensity values across the individual line scans depicted by the white line in the merged images. Top right graph is CV values of Kv2.2 or Kv2.1 measured from interphase (I phase) or M phase cells. Bottom right graph is ER-PM junction (EPJ) size measured from interphase cells coexpressing mCherry-SEC61β and either Kv2.2 or Kv2.1. Bars are mean ± SD. See Figure 11-Tables 1-2 for values and statistical analyses.

In interphase COS-1 cells lacking Kv2.2 or Kv2.1 expression, mCherry-SEC61β was present as reticular tubules and puncta (Figure 11), and expression of Kv2.2 but not Kv2.1 reorganized ER-PM junctions, such that interphase cells expressing Kv2.2 had a ER-PM junctions significantly larger than cells without Kv2 channel expression or cells expressing Kv2.1 (Figure 11; Figure 11-Table 2). In contrast, mean ER-PM junction cluster size was not significantly different in cells without or with Kv2.1 expression (Figure 11; Figure 11-Table 2). Taken together, these data demonstrate that the ability of Kv2.2 to impact ER-PM junctions does not exhibit the reversible, cell cycle-dependent modulation as seen for Kv2.1, a distinction that could impact Kv2-associated ER-PM junctions in cells primarily expressing one or the other mammalian Kv2 channel paralog.

### Eliminating Kv2 channel expression *in vivo* impacts RyR-containing ER-PM junctions in brain neurons

As detailed above, clustered endogenous Kv2 channels colocalize with RyR-containing ER-PM junctions in brain neurons *in situ* and in culture, and exogenously expressing either Kv2.2 or Kv2.1 can remodel ER-PM junctions in CHNs and heterologous cells. We next tested whether eliminating Kv2 channel expression in knockout mice impacts the spatial organization of RyR-containing ER-PM junctions in brain neurons, taking advantage of the availability of Kv2.1 (30, 83) and Kv2.2 (84) knockout mice, and double knockout mice (41). We immunolabeled brain sections from these mice and from wild-type controls for Kv2.2, Kv2.1 and RyR, and analyzed RyR clusters in hippocampal CA1 pyramidal neurons, which express both Kv2.2 and Kv2.1 (23, 39, 41, 83). As shown in Figure 12, while there were no significant changes in the spatial characteristics of RyR clusters in the samples from the single Kv2 knockout mice when compared to those from wild-type mice, the size of RyR clusters in CA1 pyramidal neurons was significantly reduced in the samples from the double Kv2 knockout mice (Figure 12-Table 1). This supports an *in vivo* role for Kv2 channels in organizing RyR-containing ER-PM junctions in brain neurons.

**Figure 12.**
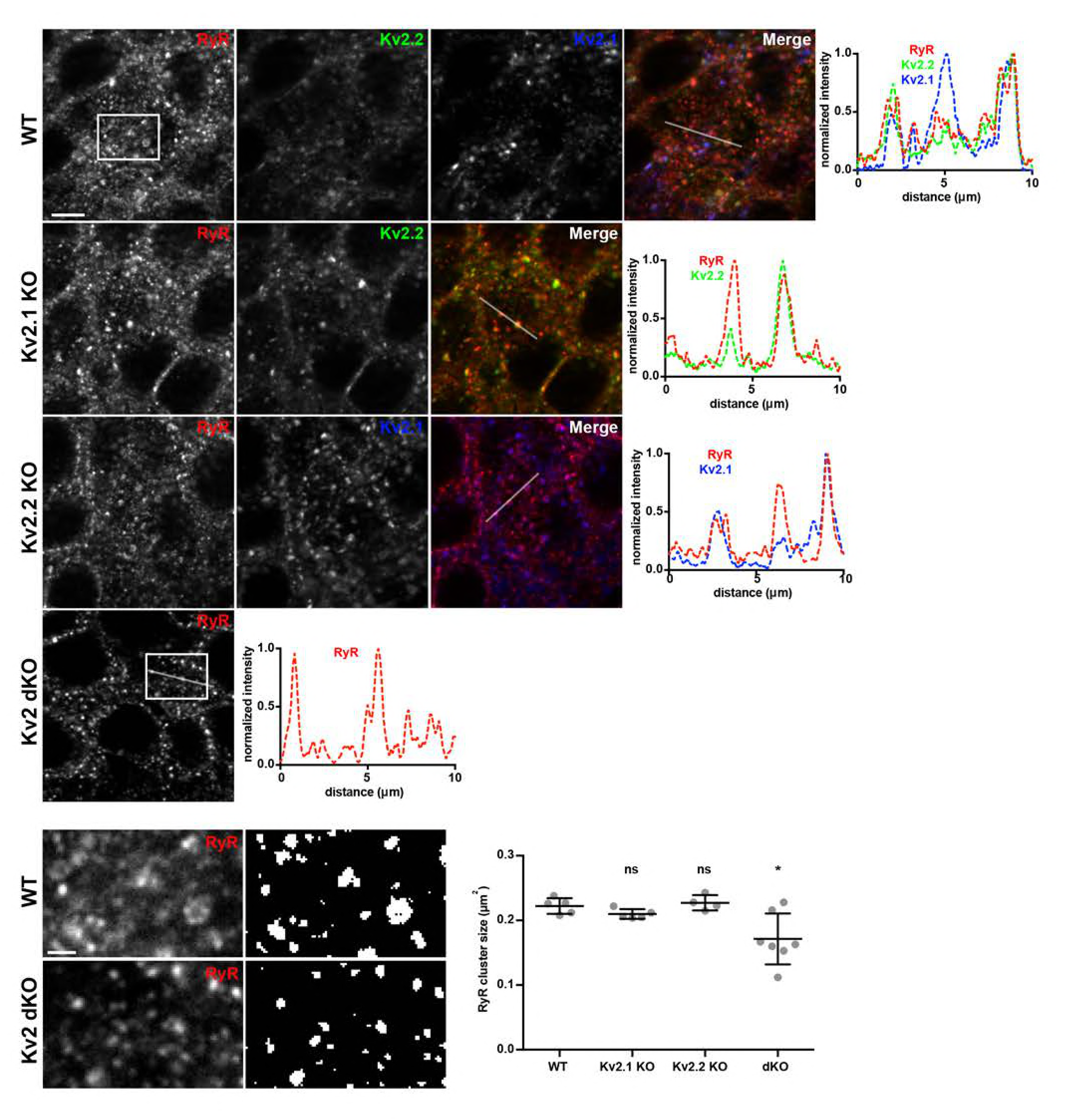
Genetic ablation of Kv2.2 and Kv2.1 alters RyR localization in mouse brain neurons. Projected z-stack images of CA1 hippocampus from brain sections of wild-type (WT), Kv2.1 knockout (Kv2.1 KO), Kv2.2 knockout (Kv2.2 KO), or Kv2.1 and Kv2.2 double knockout (Kv2.1/Kv2.2 dKO) mice immunolabeled for RyR, Kv2.2, and Kv2.1. Top row shows RyR, Kv2.2, and Kv2.1 immunolabeling from WT mouse. Second row shows immunolabeling RyR and Kv2.2 immunolabeling from Kv2.1 KO mouse. Third row shows RyR and Kv2.1 immunolabeling from Kv2.2 KO mouse. Fourth row shows RyR immunolabeling from Kv2.1/Kv2.2 dKO mouse. Scale bar in WT RyR panel is 10 µm and holds for all panels in set. Panels to the right of each row are the corresponding normalized fluorescence intensity values across the individual line scans depicted by the white line in the merged images. Bottom panels are enlarged selections of RyR- labeling of WT and Kv2.1/Kv2.2 dKO images as indicated by boxes. Scale bar in WT RyR inset panel is 1.25 µm and holds for all panels in set. Panels to the right are binary/thresholded masks generated during analysis of RyR cluster size. Graph to the right are measurements of individual RyR cluster sizes. Bars are mean ± SD. See Figure 12-Table 1 for values and statistical analyses.

## Discussion

Our results presented here demonstrate that members of the Kv2 channel family have as a conserved function the ability to organize ER-PM junctions, which is unique among all PM proteins studied to date. We show that Kv2.2 ion channels localize to ER-PM junctions on somata, proximal dendrites and the AIS in brain neurons. Experiments in CHNs, and in heterologous HEK293T and COS-1 cells show that Kv2.2 channels function in themselves to organize ER-PM junctions. We show that the ability to organize ER-PM junctions is a nonconducting function of mammalian Kv2 ion channels that requires their PM clustering. Moreover, elimination of Kv2 expression in knockout mice leads to altered ER-PM junctions in brain neurons. The conserved ER-PM junction-organizing function of Kv2.2 and Kv2.1 makes the Kv2 family of mammalian ion channels the first family of PM proteins whose expression is sufficient to reorganize ER-PM junctions. Separation-of-function mutants in Kv2.2 and Kv2.1 reveal that this conserved function is independent of their well-established canonical function as ion conducting channels regulating electrical signaling in neurons and non-neuronal cells, but entirely dependent on their clustering in the PM as mediated by a conserved motif in their respective cytoplasmic C-termini whose mutation does not impact their function as ion channels. Kv2-containing ER-PM junctions are found at sites deficient in components of the cortical actin cytoskeleton, which contributes to but is not the sole determinant of the overall spatial organization of Kv2 channel-containing ER-PM junctions. Kv2-containing ER-PM junctions are found associated with those containing diverse ER tethers that mediate ER and PM contacts, suggesting that ER-PM junctions formed by Kv2 channels and these ER tethers may structurally and functionally overlap in cells in which they are coexpressed. While Kv2.2 and Kv2.1 share a conserved function in organizing ER-PM junctions, they are distinct in that Kv2.2 ER-PM junctions are stable in the face of changes in cellular environment that result in reversible disruption and subsequent reformation of Kv2.1 ER-PM junctions. That Kv2.2 and Kv2.1 have distinct patterns of cellular expression suggests that the highly similar yet distinct functions of these mammalian Kv2 channel paralogs in organizing ER-PM junctions would distinctly impact the structure, function and regulation of ER-PM junctions in the types of neurons and non-neuronal cells in which they are differentially expressed.

Endogenous Kv2 channels are present in large PM clusters in diverse classes of brain neurons [Kv2.1: (39, 41, 69); Kv2.2: (18, 39, 41, 42)]. That in certain brain neurons and in neurons in culture we found clusters of Kv2.2 at sites containing high densities of associated ER-localized RyRs supports that these clusters represent native Kv2.2-containing ER-PM junctions, and that these sites are associated with neuronal Ca^2+^ signaling. Moreover, that elimination of expression of both Kv2 channels leads to changes in the spatial organization of RyR-containing ER-PM junctions in brain neurons suggests that Kv2 channels play a role in the structural organization of these Ca^2+^ signaling microdomains. Although both Kv2.2 and Kv2.1 are unique among mammalian PM proteins in being capable of organizing ER-PM junctions, their distinct cellular expression patterns in brain and in other mammalian tissues, together with their distinct phospho-dependent regulation, may contribute to the unique phenotypes seen in mice upon knockout of either Kv2.2 [altered sleep wake cycles (85)] or Kv2.1 [neuronal and behavioral hyperexcitability (83)]. The relative contribution of the separate functions of Kv2 channels as ion conducting channels shaping membrane excitability, and as structural organizers of ER-PM junctions, to the behavioral phenotypes of these mice is as of yet unknown.

Our data using a strategically selected set of separation-of-function point mutants support that recruitment/stabilization of ER-PM junctions is a nonconducting and physical function of Kv2 channels that relies on their clustering. Both Kv2.2 and Kv2.1 are *bona fide* PM voltage-gated K^+^ channels whose ion conducting function underlies the bulk of the delayed rectifier K^+^ current in various classes of neurons (20-22, 36). Moreover, acute pharmacological inhibition of Kv2 channels impacts neuronal excitability and the characteristics of action potentials (22-25, 86, 87). Our findings that the ability to organize ER-PM junctions is a nonconducting function of Kv2 channels is intriguing given previous findings that the bulk of exogenous Kv2.1 expressed in either heterologous cells or neurons may be present in a nonconducting state (88-90). That ion channels can have diverse nonconducting functions distinct from their canonical ion conducting roles is an emerging theme in biology, with nonconducting roles as cell adhesion molecules, as enzymes or as scaffolds for enzymes, as voltage sensors for intracellular events through conformational coupling, etc. [reviewed in (91)]. Studies in pancreatic beta cells support such a nonconducting function for Kv2.1 in regulating insulin secretion (92). As this nonconducting role is dependent on Kv2.1 clustering (93) suggests a potential function for Kv2.1 in organizing ER-PM junctions in beta cells, which have been proposed to play an important role in glucose-stimulated insulin secretion (94, 95). Recent studies employing whole exome sequencing have led to identification of encephalopathic epilepsy patients with *de novo* mutations in the KCNB1 gene that encodes Kv2.1. While the bulk of these disease-associated mutations are in the voltage-sensing and pore domains that are crucial to the canonical function of Kv2.1 as a *bona fide* Kv channel [*e.g.*, (26- 28)], a subset are nonsense mutations that result in a truncated cytoplasmic C-terminus (29, 96). While the cytoplasmic C-terminus plays a modulatory role in regulating activation gating of Kv2.1 channels (97-99), the most obvious effect of these nonsense mutations that eliminate the PRC domain is to disrupt the clustering of Kv2.1 (35, 37, 39, 41, 51, 65) and presumably organization of ER-PM junctions. Generating mouse models that express the separation-of-function mutations used here to selectively disrupt Kv2.1 conduction and clustering may lead to insights into these distinct classes of disease-associated mutations, as well as the relative contributions of the separable electrical and structural roles of Kv2 channels in normal physiology.

Our results show that both members of the Kv2 family of ion channels can in themselves organize ER-PM junctions. As these are the first mammalian PM proteins with this function suggests Kv2 channels use a molecular mechanism distinct from all other known classes of ER-PM junction organizers (*i.e.*, members of the E-Syt, JP and STIM families), which are ER tethers that bind specific lipids present in the inner leaflet of the PM, although STIM family members also exhibit conditional interaction with PM Orai proteins (1, 10). That both Kv2.2 and Kv2.1 expression are sufficient to remodel ER-PM junctions in the absence of their ion conducting functions, and *via* a mechanism that requires an intact PRC motif, suggests that both Kv2 family members act through the same mechanism. One plausible mechanism is that the Kv2-specific cytoplasmic C-termini, and the PRC motif, in particular, interact directly with an ER-localized protein or lipid binding partner. That these Kv2 channels are capable of forming clusters localized at ER-PM junctions in diverse cell types including brain neurons of diverse mammalian species *in situ* and in culture [*e.g.*, (34-36, 39, 41, 42, 48, 52, 80, 81, 90, 100), etc.], spinal motor neurons (101) and in non-neuronal heterologous cells such as human HEK293 (39, 41), monkey COS-1 (47) and canine MDCK (35) kidney cells, rat PC12 pheochromocytoma cells (102), and hamster CHO ovary cells (47) suggests that the underlying mechanism involves components highly conserved across diverse mammalian species and cell types. Moreover, should the mechanism involve binding to a specific ER protein, the protein should also be highly expressed across these diverse cell types, as the formation of Kv2 clusters and recruitment of ER-PM junctions is not obviously saturable, such that the higher the level of Kv2.2 or Kv2.1 expression, the larger the clusters and associated ER-PM junctions (47, 48). We note that the clustering of Kv2.1 is conditional in many if not all of these cell backgrounds, such that it occurs in M-phase but not interphase COS-1 cells (47), polarized MDCK cells in confluent epithelial monolayers but not in nonpolarized MDCK cells in low density culture (35), and in PC12 cells before but not after differentiation with nerve growth factor (102). Moreover, in neurons and HEK293 cells, changes in protein kinase (54, 99) and protein phosphatase (39, 52, 53, 80, 103, 104) activity leads to changes in Kv2.1 phosphorylation state and clustering, and presumably its association with ER-PM junctions. As such, the mechanism whereby Kv2.1 organizes ER-PM junctions may involve regulation *via* dynamic changes in phosphorylation state, including in critical serine residues within the PRC domain itself (37, 47). That the Kv2.2 PRC domain contains these same serine residues suggests that should phosphorylation at these sites be required for Kv2.2 clustering and ER-PM junction reorganization, that this phosphorylation is more constitutive than the dynamically-regulated phosphorylation of Kv2.1.

That Kv2-containing ER-PM junctions can colocalize with all known members of the E-Syt and STIM families, as well as JP2 and JP4, suggests potential overlap with these distinct classes of ER-PM junctions in coexpressing mammalian cells. One explanation of these findings is that these ER-localized PM tethers, by virtue of their ER localization are passively recruited along with other ER proteins such as Sec61β to Kv2-containing ER-PM junctions. However, the lack of association of Kv2-containing ER-PM junctions and those generated *via* the rapamycin-triggered coupling of Lyn11-FRB and CB5-FKBP would argue against a promiscuous presence of Kv2 channels at any ER-PM junction. As the tethering of E-Syts, JPs and STIMs to the PM occurs at least in part on their binding to lipids on the PM inner leaflet (1), another possible explanation for the robust colocalization between Kv2-containing ER-PM junctions and these ER tethers is that Kv2 clustering results in a distinct lipid microenvironment in the PM inner leaflet at or near these clusters. Changes in the local lipid environment at/near Kv2 clusters could also underlie generation of ER-PM junctions at these sites, *via* recruitment of one or more lipid-binding ER-PM tethers. As noted above, these tethers in aggregate would need to have sufficiently robust expression across the numerous species and cell types in which endogenous and exogenous Kv2 channels are clustered. We note that our quantitative analyses of colocalization between Kv2-containing ER-PM junctions and these ER tethers suggests that despite the extensive overlap, as reported by high (≈1.0) MOC values, the intensity profiles of these proteins do not uniformly coincide, as reported by significantly lower paired PCC measurements (105). That there is heterogeneity in ER-PM junctions within the same cell is consistent with the variable co-occurrence of Kv2.2 and Kv2.1 clusters with RyR clusters between and within different classes of mammalian brain neurons (38, 48, 106). This concept is further supported by the lack of colocalization between Kv2-containing ER-PM junctions and those formed *via* triggered coupling of Lyn11/CB5. That little is known of the subcellular localization of the different members of the E-Syt, JP and STIM families endogenously expressed in mammalian brain neurons makes it difficult to understand the relationship between the native ER-PM junctions formed by these ER tethers and those containing Kv2 channels.

That LatA treatment impacted the characteristics of both Kv2-and Lyn11/CB5-containing ER-PM junctions but did not lead to their fusion suggests that the actin cytoskeleton is not the only determinant of their distinct spatial organization. The effects of actin disruption on Kv2-containing ER-PM junctions, and that they are localized to zones at the cell cortex depleted in actin and actin-interacting proteins, suggests a role for the actin cytoskeleton in shaping their spatial characteristics. This is consistent with previous studies demonstrating that Kv2.1 clusters on the axon initial segment of brain neurons are specifically localized to ankyrinG-deficient “holes” (69), and that disruption of the actin cytoskeleton impacts clustering of Kv2.1 (44, 46). Recent studies reveal that the STIM1:Orai1 complex at the immune synapse (107) and HeLa cell ER-PM junctions labeled with the reporter MAPPER (108) are also present in actin-poor zones, and that disruption of the actin cytoskeleton altered the distribution and dynamics of these HeLa cell ER-PM junctions (108). Depletion of ER Ca^2+^ stores triggers a conditional association of the STIM1: Orai1 complex with Kv2-containing ER-PM junctions (51). We also found that in the absence of exogenously expressed STIM1, store depletion triggered a significant increase in colocalization between Kv2.2 and Orai1, presumably due to endogenous STIM1 expression in HEK293T cells (75). That both ER (RyR) and PM (Orai1) Ca^2+^ channels colocalize with Kv2-containing ER-PM junctions suggests a structural role for Kv2 channels in regulating sites important in neuronal Ca^2+^ homeostasis above and beyond their established role in shaping membrane excitability. Future studies will define the respective contributions of the separate yet highly conserved conducting and nonconducting roles of Kv2 channels in impacting cellular physiology, and how this is disrupted in pathological conditions that may exert their effects through distinct impacts on these broadly and highly expressed ion channels.

## Materials and methods

### Preparation of mouse brain sections for immunohistochemistry

All procedures involving mice were approve by the University of California Davis Institutional Animal Care and Use Committee and were performed in strict accordance with the Guide for the Care and Use of Laboratory Animals of the NIH. All mice were maintained under standard light-dark cycles and allowed to feed and drink ad libitum. Kv2.1-KO mice (RRID:IMSR_MGI:3806050) have been described previously (30, 83), and were generated from breeding of Kv2.1^+/-^ mice that had been backcrossed on the C57/BL6J background (RRID:IMSR_JAX:000664). Kv2.2-KO mice (84, 85) were obtained from Drs. Tracey Hermanstyne and Jeanne Nerbonne. All Kv2.2-KO mice used here were obtained from heterozygotic crosses in the C57/BL6J background (RRID:IMSR_JAX:000664). Double knockout mice for Kv2.1/Kv2.2 (Kv2 dKO) were generated by crossing Kv2.1^+/-^ and Kv2.2^-/-^ mice. Both male and female mice were used, over 12 weeks old. Littermates were used when available. Mice were deeply anesthetized with 90 mg/kg Na-pentobarbital salt (Sigma Cat# P3761) in 0.9% NaCl solution through intraperitoneal injections, followed by boosts as needed. Once mice were completely anesthetized, they were transcardially perfused with a brief prefix wash with 4.5 ml of ice cold PBS [150 mM NaCl, 10 mM sodium phosphate buffer (PB), pH 7.4] containing 10 U/ml heparin, followed by an ice-cold fixative solution of 4% formaldehyde (freshly prepared from paraformaldehyde, Sigma Cat# 158127) in 0.1 M sodium PB, pH 7.4, using a volume of 1 ml fixative solution per gram of mouse weight.

Following perfusions, brains were removed from the skull and cryoprotected in 10% sucrose, 0.1 M PB overnight at 4°C, then transferred to a solution of 30% sucrose, 0.1 M PB until they sank to the bottom of the tube (24–48 h). Following cryoprotection, all brains were frozen, and cut on a freezing stage sliding microtome (Richard Allen Scientific) to obtain 30-µm-thick sagittal sections. Sections were collected in 0.1 M PB and processed for immunohistochemistry (IHC) as free-floating sections.

### Multiplexed fluorescence immunohistochemistry

Multiplex immunofluorescence labeling of mouse brain sections was performed essentially as previously described (109). Briefly, free-floating sections were washed 3× in 0.1 M PB and 10 mM sodium azide at room temperature with slow agitation. All subsequent incubations and washes were at room temperature with slow agitation, unless stated otherwise. Sections were incubated in blocking buffer (10% goat serum in 0.1 M PB, 0.3% Triton X-100, and 10 mM sodium azide) for 1 h. Immediately after blocking, sections were incubated with primary antibody combinations (diluted in blocking buffer) overnight at 4°C in shaker. Following incubation, sections were washed 3 x 10 min each in 0.1 M PB and incubated for 1 h with affinity-purified goat anti-rabbit and/or goat anti-mouse IgG-subclass-specific Alexa fluor-conjugated secondary antibodies and diluted in blocking buffer. Sections were labeled with the DNA-specific dye Hoechst 33258 during the secondary antibody step. After 3 x 10 min washes in 0.1 M PB, sections were mounted and dried onto gelatin-coated slides, treated with 0.05% Sudan Black Sudan Black (EM Sciences Cat# 21610) in 70% ethanol for 1.5 min, extensively washed in water, and mounted with Prolong Gold (ThermoFisher Cat# P36930). All immunolabeling reported for quantification purposes are representative of three animals (biological replicates) per genotype, except for Kv2.2 KO that included brain sections from two animals. Brain sections from all biological replicates within each experiment were labeled, treated, and mounted in parallel.

All images were acquired on a Zeiss AxioObserver Z1 microscope with an X-Cite 120 lamp as the fluorescent light source and equipped with an AxioCam MRm digital camera. High-magnification optical sections were acquired using an ApoTome structured illumination system (Carl Zeiss MicroImaging) with a 63X/1.40 NA plan-Apochromat oil immersion objective. ApoTome z-stacks were acquired and processed with Axiovision 4.8.2 acquisition software (Carl Zeiss MicroImaging, RRID: SciRes_000111). All brain sections within a given experiment and immunolabeled with the same antibody cocktail were imaged under the same conditions (objective, exposure time, lamp settings, etc.). Image processing was performed in Axiovision (Carl Zeiss MicroImaging) and Fiji v2.0.0-rc-43/1.51 (NIH). All panels in a given figure were imaged and processed identically, unless otherwise noted. High-magnification ApoTome z-stacks were opened for analysis as raw image files in Fiji (NIH) using the Bio-Formats library importing plugin (Linkert et al., 2010). Quantification was done using single optical z-sections. All statistical analyses of immunolabeling were performed in Prism (GraphPad).

Quantification of RyR immunolabeling was performed in FIJI. Images were first background subtracted; background levels were determined from “no primary antibody” immunolabeling controls for each animal, and mathematically subtracted from paired images of RyR labeling, and images were converted to 8-bit. An ROI selection was made to include cell bodies of neurons in the pyramidal cell layer of CA1, and the image was automatically converted into a binary mask using auto local thresholding (110). RyR cluster size was quantified automatically using the “analyze particles” function in FIJI. Particles smaller than 0.06 μm^2^ were excluded from this analysis.

### Culture and transfection of rat hippocampal neurons

All procedures involving rats were approved by the University of California Davis Institutional Animal Care and Use Committee and were performed in strict accordance with the Guide for the Care and Use of Laboratory Animals of the NIH. All rats were maintained under standard light-dark cycles and allowed to feed and drink *ad libitum*. Hippocampi were dissected from embryonic day 18 rat embryos and dissociated enzymatically for 20 min at 37 °C in 0.25% (w/v) trypsin (ThermoFisher Cat# 15050065) in HBSS and dissociated mechanically by triturating with glass polished Pasteur pipettes. Dissociated cells were suspended in Neurobasal (Invitrogen Cat# 21103-049) supplemented with 10% FBS (Invitrogen Cat# 16140071), 2% B27 (Invitrogen Cat# 17504044), 1% GlutaMAX (Invitrogen Cat# 35050061), and 0.001% gentamycin (Gibco Cat #1570-064) and plated at 60,000 cells per dish in glass bottom dishes (MatTek Cat# P35G-1.5- 14-C), or number 1.5 glass coverslips, coated with poly-L-lysine (Sigma Cat# P2636). At 4-7 DIV, cytosine-D-arabinofuranoside (Millipore Cat# 251010) was added to inhibit non-neuronal cell growth. CHNs were transiently transfected at DIV 5-10 using Lipofectamine 2000 (Invitrogen Cat# 11668019) for 1.5 hours as previously described (37). CHNs were imaged 40-48 hours post transfection.

### Heterologous cell culture, reagents, and transfection

HEK293T cells were maintained in Dulbecco’s modified Eagle’s medium supplemented with 10% Fetal Clone III (HyClone Cat# SH30109.03), 1% penicillin/streptomycin, and 1X GlutaMAX (ThermoFisher Cat# 35050061) in a humidified incubator at 37 °C and 5% CO_2_. COS-1 cells were maintained in Dulbecco’s modified Eagle’s medium supplemented with 10% Bovine Calf Serum (HyClone Cat# 16777-206), 1% penicillin/streptomycin, and 1X GlutaMAX in a humidified incubator at 37 °C and 5% CO_2_. HEK293T cells were transfected as previously described (39). Briefly, HEK293T cells were split to 15% confluence on glass bottom dishes (MatTek Cat# P35G- 1.5-14-C) coated with poly-L lysine then transiently transfected using Lipofectamine 2000 (Invitrogen) transfection reagent following the manufacturer’s protocol. HEK293T cells were transiently transfected in DMEM without supplements, then returned to regular growth media 4 hours after transfection. HEK293T cells were imaged 40-48 hours post-transfection. COS-1 cells were transiently transfected as previously described (47). Briefly, COS-1 cells were split to 30% confluence on glass bottom dishes (MatTek Cat# P35G-1.5-14-C) coated with poly-L lysine and immediately transfected with Kv2.2, Kv2.1, and mCherry-SEC61β in DMEM with supplements. COS-1 cells were fixed and immunolabeled 40-48 hours post-transfection.

### Cell fixation, immunolabeling, and fixed-cell imaging

For experiments involving imaging of fixed cells, fixation and immunolabeling, fixation was performed as previously described (111). Briefly, HEK293T and COS-1 cells were fixed in 3.2% formaldehyde (freshly prepared from paraformaldehyde, Sigma Cat# 158127) and 0.1% glutaraldehyde (Ted Pella, Inc., Cat # 18426) for 30 minutes and room temperature, washed 3 x 5 minutes in PBS and quenched with 1% sodium borohydride in PBS for 15 minutes at room temperature. Cells were blocked and permeabilized in 4% non-fat milk powder in PBS containing 0.5 % Triton-X 100. Neurons were fixed in 4% formaldehyde in PBS for 15 minutes, washed 3 x 5 minutes in PBS and blocked and permeabilized in 4% non-fat milk powder in PBS containing 0.1 % Triton-X 100. All antibodies used in this study have been previously described (see Table 1 for a description of primary antibodies). Primary antibody incubation was performed in blocking solution for 1 hour at room temperature. Following primary antibody incubation, and 3 x 5 minute washes in blocking solution at room temperature, coverslips were immunolabeled with species- and or mouse IgG subclass-specific Alexa Fluor-conjugated goat anti-mouse IgG subclass-specific (109) or goat anti-rabbit IgG secondary antibodies (all secondary antibodies from ThermoFisher) at 1–1500 and Hoechst 33258 (ThermoFisher Cat# H1399) for one hour in blocking solution, washed 3 x 5 min in PBS, and mounted onto microscope slides using Fluoromount G (Southern Biotech Cat# 0100-01), or for samples prepared for TIRF, imaged in PBS containing ascorbate.

**Table 1.**
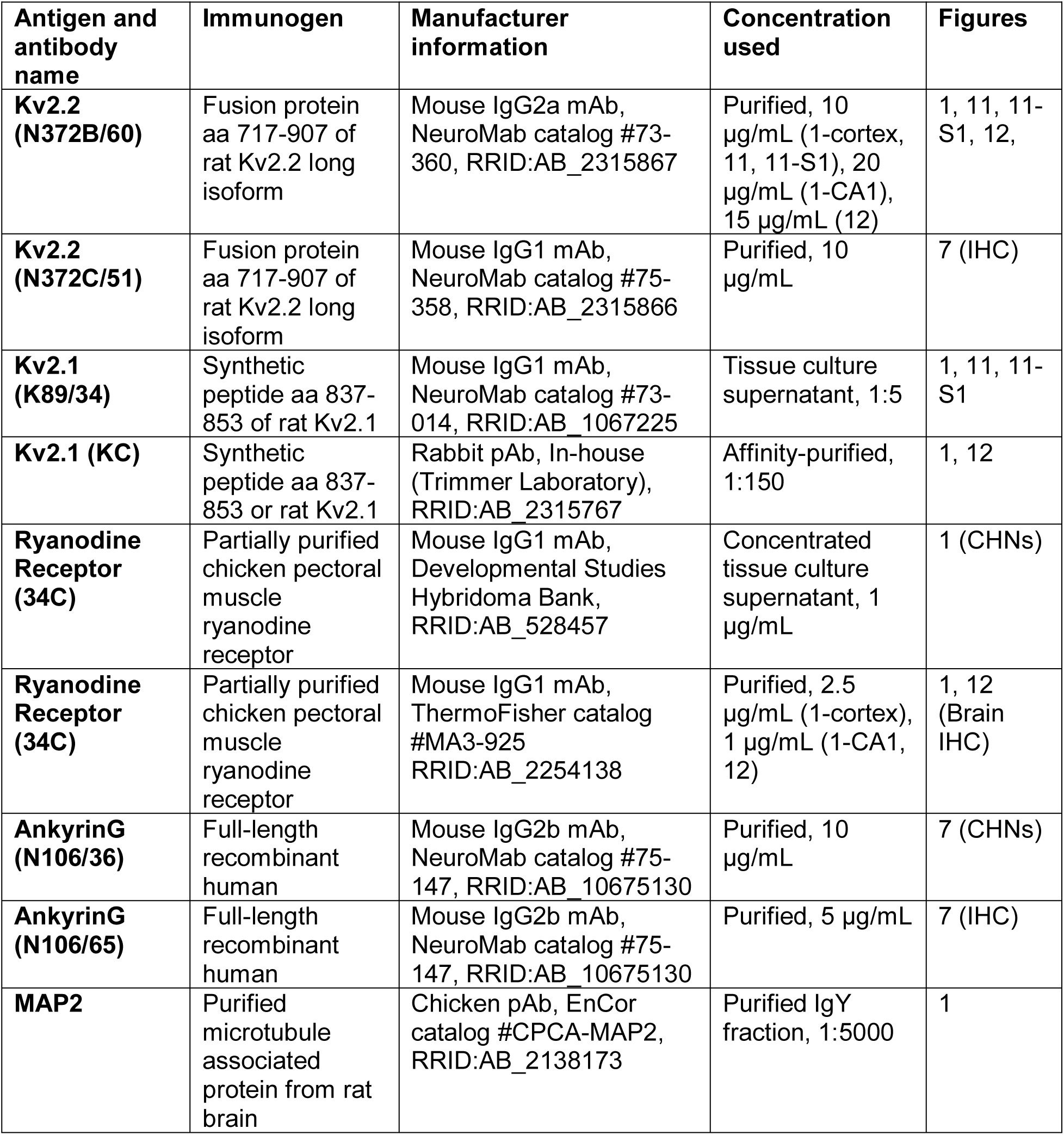
Antibody information.

For conventional fluorescence imaging (used in Figure 1E and 1G; 7A; 10F; and Figure 10-figure supplement 1) images were acquired with an AxioCam MRm digital camera installed on a Zeiss AxioImager M2 microscope or with an AxioCam HRm digital camera installed on a Zeiss AxioObserver Z1 microscope with a 63X/1.40 NA plan-Apochromat oil immersion objective or a 20X/0.8 NA plan-Apochromat objective and an ApoTome coupled to Axiovision software (Zeiss, Oberkochen, Germany). For TIRF imaging of fixed cells, imaging was identical to that used in live-cell TIRF experiments but in the absence of a heated stage/objective heater. Images were obtained with an Andor iXon EMCCD camera installed on a TIRF/widefield equipped Nikon Eclipse Ti microscope using a Nikon LUA4 laser launch with 405, 488, 561, and 647 nm lasers and a 100X PlanApo TIRF/1.49 NA objective run with NIS Elements software (Nikon). Images were collected within NIS Elements as ND2 images. For N-SIM imaging of fixed cells, images were acquired using a Hamamatsu ORCA-ER CCD camera installed on a SIM/widefield equipped Nikon Eclipse Ti microscope using an EXFO X-Cite metal halide light source and a 100X PlanApo TIRF/1.49 objective, run with NIS Elements software (Nikon). Images were collected within NIS Elements as ND2 images. SIM analysis was performed in NIS Elements. Airyscan imaging was performed with a Zeiss LSM 880 confocal laser scanning microscope (Carl Zeiss), equipped with an Airyscan detection unit, with a Plan-Apochromat 63X/1.40 Oil DIC M27 objective.

### Plasmid constructs

All novel constructs used in this study (GFP-Kv2.2, GFP-Kv2.2 P412W, DsRed-Kv2.2, GFP-Kv2.2 S605A, GFP-Kv2.1 S586A, GFP-Kv2.1 P404W) were generated using standard molecular biology approaches and confirmed by sequencing. GFP-Kv2.2 and DsRed-Kv2.2 were generated using Gibson assembly to insert full-length rat Kv2.2, also termed Kv2.2_long_ (42) into the GFP-C1 or DsRed-C1 vector (ClonTech) resulting in fusion of GFP or DsRed to the N-terminus of full-length rat Kv2.2. GFP-Kv2.1 S586A, GFP-Kv2.1 P404W, and GFP-Kv2.2 S605A were generated *via* site directed point mutagenesis utilizing a quick change PCR reaction of GFP-Kv2.1 (48) or GFP-Kv2.2, respectively, or *via* Gibson assembly. GFP-Kv2.2 P412W was generated at Mutagenex. Plasmids encoding DsRed2-ER5 and mCherry-actin were a generous gift from Dr. Michael Davidson (Addgene plasmids # 55836 and 54965). The plasmid encoding ankG-mCherry was a generous gift from Dr. Benedicte Dargent (Addgene plasmid #42566). The plasmids encoding BFP-SEC61β, mCherry-SEC61β, and BFP-STIM1 were a generous gift from Dr. Jodi Nunnari (University of California, Davis). The plasmid encoding GFP-JP2 was a generous gift from Dr. Fernando Santana (University of California, Davis). The plasmid encoding mCherry-E- Syt1-3 was a generous gift from Dr. Pietro De Camilli (Yale University School of Medicine). The plasmid encoding mCherry-JP4 was a generous gift from Dr. Yousang Gwack (University of California, Los Angeles). The plasmids encoding mCherry STIM1, 2, and 2 and GFP-Orai1 were a generous gift from Dr. Richard Lewis (Stanford University). The plasmids encoding CFP- CB5-FKBP and Lynn11-FRB (76) were a generous gift from Dr. Eamonn Dickson.

### Live cell Guangxitoxin labeling

The GxTX peptide used in surface labeling was synthesized at the Molecular Foundry of the Lawrence Berkeley National Laboratory under US Department of Energy contract no. DE-AC02- 05CH11231. HEK293T cells were surface labeled with 1 μM GxTX as previously described (64) and imaged in TIRF as described below but in physiological saline solution (4.7 mM KCl, 146 mM NaCl, 2.5 mM CaCl_2_, 0.6 mM MgSO_4_, 1.6 mM NaHCO_3_. 0.15 mM NaH_2_PO_4_, 20 mM HEPES, pH 7.4) containing 8 mM glucose and 0.1 mM ascorbic acid) containing 0.1% BSA.

### Live cell TIRF imaging

Total internal reflection fluorescence (TIRF) imaging was performed at the UC Davis MCB Imaging Facility. Live transfected HEK293T cells cultured on glass bottom dishes were imaged in a physiological saline solution (4.7 mM KCl, 146 mM NaCl, 2.5 mM CaCl_2_, 0.6 mM MgSO_4_, 1.6 mM NaHCO_3_. 0.15 mM NaH_2_PO_4_, 20 mM HEPES, pH 7.4) containing 8 mM glucose and 0.1 mM ascorbic acid). Cells were maintained at 37°C during the course of imaging with a heated stage and objective heater. For experiments involving Latrunculin A (ThermoFisher Scientific, Cat# 428021100UG) treatment, Latrunculin A was diluted to 20 μM in imaging saline and added by pipette, to glass bottom dishes already containing imaging saline, to a final concentration of 10 μM. For experiments involving thapsigargin (Millipore, Cat# 586005-1MG) treatment, thapsigargin was diluted to 4 μM in imaging saline and added by pipette, to GLASS BOTTOM dishes already containing imaging saline, to a final concentration of 2 μM. For experiments involving rapamycin (Sigma, Cat# R8781-200UL) treatment, rapamycin was diluted to 10 μM in imaging saline and added by pipette to glass bottom dishes already containing imaging saline, to a final concentration of 5 μM. Images were obtained with an Andor iXon EMCCD camera installed on a TIRF/widefield equipped Nikon Eclipse Ti microscope using a Nikon LUA4 laser launch with 405, 488, 561, and 647 nm lasers and a 100X PlanApo TIRF, 1.49 NA objective run with NIS Elements software (Nikon). Images were collected within NIS Elements as ND2 images.

### Cell culture and transfection for electrophysiology

All cell lines were grown in a humidified incubator at 37°C and 5% CO_2_. HEK293T cells were maintained in Dulbecco’s modified Eagle’s medium supplemented with 10% fetal bovine serum (HyClone Cat # SH30109.02) and 1% penicillin/streptomycin. Transfections were performed with Lipofectamine 2000 (Life Technologies Cat #11668-027). Cells were plated overnight prior to transfection and allowed to grow to ≈40% confluency. Lipofectamine was diluted, mixed, and incubated in Opti-MEM (Gibco Cat #31965-062) in a 1:100 ratio for 5 minutes. Concurrently, 1 μg of plasmid DNA and Opti-MEM were mixed in the same fashion. After incubation, the DNA and Lipofectamine 2000 mixtures were combined, triturated, and allowed to incubate for 20 minutes. The transfection cocktail was added to cells for 5 hours before the media was replaced. For experiments in Figure 9, 1 μg of GFP-Kv2 or a peGFP-C1 plasmid were used. For experiments in Figure 10, 0.2 μg of GFP-Kv2 plasmids were diluted with 0.8 μg pcDNA3 plasmid.

### Electrophysiology

Whole cell voltage clamp was used to measure currents from HEK293T cells expressing GFP- Kv2.2, GFP-Kv2.2 P412W, GFP-Kv2.1, GFP-Kv2.1 P404W, or GFP as a control. On the day of the experiment (two days after transfection), transiently transfected cells were detached with trypsin and plated onto cell culture-treated polystyrene dishes for electrophysiological measurements. The external (bath) solution contained (in mM): 3.5 KCl, 155 NaCl, 10 HEPES, 1.5 CaCl_2_, 1 MgCl_2_, adjusted to pH 7.41 with NaOH. The internal (pipet) solution contained (in mM): 35 KOH, 70 KCl, 50 KF, 50 HEPES, 5 EGTA adjusted to pH 7.2 with KOH. Liquid junction potential (calculated to be 7.8 mV) was not corrected for. Borosilicate glass pipettes (Sutter Instruments, Cat #BF150-110-10HP) with resistance less that 3 M were used to patch the cells. Recordings were at room temperature (22–24 °C). Voltage clamp was achieved with an Axon Axopatch 200B amplifier (MDS Analytical Technologies, Sunnyvale, CA) run by PATCHMASTER software, v2×90.2 (HEKA, Bellmore, NY). Holding potential was −80 mV. Capacitance and Ohmic leak were subtracted using a P/5 protocol. Recordings were low pass filtered at 10 kHz and digitized at 100 kHz. Voltage clamp data were analyzed and plotted with IGORPRO software, version 7 (Wavemetrics, Lake Oswego, OR). Current amplitudes at each voltage were the average from 0.19-0.20 s after voltage step. In the experiments plotted in Figure 9, series resistance compensation was not used. The estimated series resistance in these experiments ranged from3-8 MΩ, which is predicted to result in substantial cell voltage errors for conducting channels. For quantitative comparison of current levels and voltage activation (Figure 10), we improved control of intracellular voltage by reducing the amount of DNA transfected (described above), partially blocking the K^+^ currents with tetraethylammonium (TEA) and using series resistance compensation. For experiments shown in Figure 10 on HEK293T cells expressing GFP-Kv2.2, GFP-Kv2.2 S605A, GFP-Kv2.1, or GFP-Kv2.1 S586A, the following modifications were made. The internal (pipet) solution contained (in mM): 140 KCl, 13.5 NaCl, 1.8 MgCl2, 0.09 EGTA, 4 Na-ATP, 0.3 Na-GTP, and 9 HEPES, adjusted to pH 7.2 with KOH. The external (bath) solution contained (mM): 3.5 KCl, 155 TEA-Cl, 1.5 CaCl_2_, 1 MgCl_2_, 10 HEPES, and 10 glucose adjusted to pH 7.42 with NaOH. 155 mM extracellular TEA is predicted to inhibit at least 97% of Kv2.1 current at 0 mV [see (112-114)]. A calculated liquid junction potential of 7.6 mV was corrected. Pipette tips were coated with Sylgard 184 (Dow Corning Cat #2010518, Midland, MI) and fire polished. Series resistance compensation with lag set to 10 µs was used to constrain calculated voltage error to ≤ 10 mV. Conductance was measured from the amplitude of outward tail currents averaged from the end of any capacitance transient until 2 ms after stepping to 0 mV from the indicated voltage. Fits with the fourth power of a Boltzmann distribution are described previously, where *V*_*mid*_ is the voltage where the function reaches half maximal conductance, and *z* is valence in units of elementary charge (*e*^+^) of each of the four independent voltage sensors (115). Conductance data shown are normalized to the maximal conductance of the Boltzmann fit.

### Image analysis and statistics

All colocalization analyses were performed within Nikon NIS Elements using ND2 files. An ROI was drawn within a cell of interest and PCC and MOC values were collected. Measurements of structure sizes were quantified automatically within FIJI essentially as previously described (111). ND2 files of DsRed2-ER5 or BFP-SEC61β collected in TIRF were imported directly into FIJI, background subtracted, converted into an 8-bit image, and automatically converted into a binary mask using auto local thresholding (110). An ROI with identical dimensions and containing an area of 60.6 μm^2^ was drawn within each cell analyzed. The number of individual ER-PM junctions, average ER-PM junction size, and percent PM occupancy were quantified automatically using the “analyze particles” function in FIJI. Signals smaller than 0.04 μm^2^ were excluded from this analysis. An identical approach was taken in whole cell analysis.

Quantification of Kv2 cluster sizes was performed similarly. ND2 files of GFP-Kv2.1, GFP- Kv2.1 P404W, GFP-Kv2.2, or GFP-Kv2.2 P412W collected in widefield and deconvolved in NIS elements were imported directly into FIJI, converted into an 8-bit image, and automatically converted into a binary mask using auto local thresholding (110). Kv2 cluster size was quantified automatically using the “analyze particles” function in FIJI. For scatterplot generation of ER-PM junction and Kv2 cluster sizes (Figure 3J), ND2 files were imported directly into FIJI, background subtracted using a rolling ball radius of 10 pixels and converted into an 8-bit image. Images were converted into binary masks and manually subjected to erosion operations designed to separate objects as previously described (111). Care was taken to ensure that the resulting binary image was comparable to the original image. The areas of these structures were quantified automatically using the “analyze particles” function in FIJI. Areas from 10-20 overlapping structures from each cell were paired as coordinates. In cases were more than one structure overlapped, the areas of the overlapping structures were summed as a single coordinate.

Coefficient of variation is defined as standard deviation of intensity divided by mean intensity as previously described (39, 65). Quantification of coefficient of variation and intensity measurements were collected in FIJI. An ROI was drawn around a cell and standard deviation of intensity, and mean intensity values were collected.

For line scan analysis of fluorescence intensity, raw intensity values were collected within FIJI and normalized to the maximum value collected.

Analysis of DsRed2-ER5 velocity was performed in MATLAB (MathWorks) using the PIVlab toolkit (116) as previously described (51). Briefly, successive frames (captured at 31.25 Hz) of DsRed2-ER5 expression in HEK293T cells transfected with DsRed2-ER5 alone or cotransfected with GFP-Kv2.1, GFP-Kv2.2, GFP-Kv2.1 P404W, or GFP-Kv2.2 P412W, were collected in TIRF. Images were converted into BMP file format and 1 out of every 10 frames (creating a time lapse of 320 ms) were imported into PIVlab. Contrast limited adaptive histogram equalization (contrast enhancement) was engaged, and frame pairs were analyzed with 3 successive passes, utilizing interrogation areas of 64, 32, and 16 pixels. From an ROI drawn within the center of each cell analyzed, average velocity magnitude values (reported as pixels per frame) were collected.

For all analysis, values were imported into GraphPad Prism for presentation and statistical analysis as noted. For IHC experiments, we define biological replicates as individual animals. The datasets in this manuscript involving IHC contain biological replicates. For experiments performed with cells in culture, we define biological replicates as experiments performed on different days, and technical replicates as experiments performed on the same day. The datasets in this manuscript involving cells in culture contain biological and/or technical replicates.

## Figure legends

**Figure 5-figure supplement 1.**
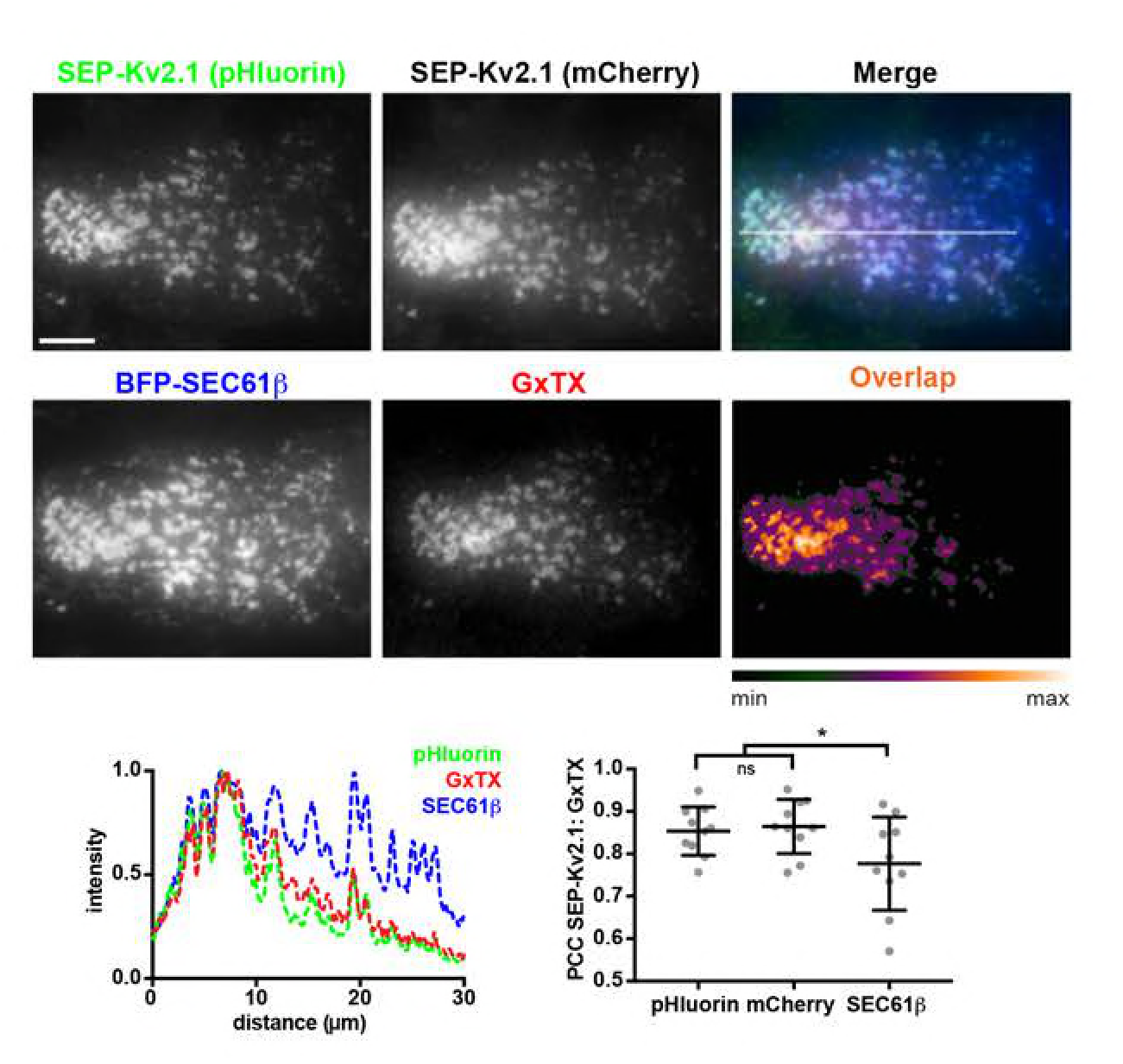
ER-PM junction-localized Kv2.1 channels are expressed on the cell surface. Top panels. TIRF images of live HEK293T cells coexpressing SEP-Kv2.1 and BFP-SEC61β and surface labeled with GxTX-633. The merged image shows SEP-Kv2.1 (pHluorin), BFP-SEC61β, and GxTX-633. Scale bar is 5 µm and holds for all panels. Heat map shows overlap of SEP-Kv2.1 (pHluorin) and GxTX-633 pixels. Bottom left panel shows the normalized fluorescence intensity values across the line scan depicted by the white line in the merged image. Graph on bottom right shows PCC values between GxTX and SEP-Kv2.1 or BFP-SEC61β. Bars are mean ± SD. See Figure 5-Table 1 for values and statistical analyses.

**Figure 6-Figure supplement 1.**
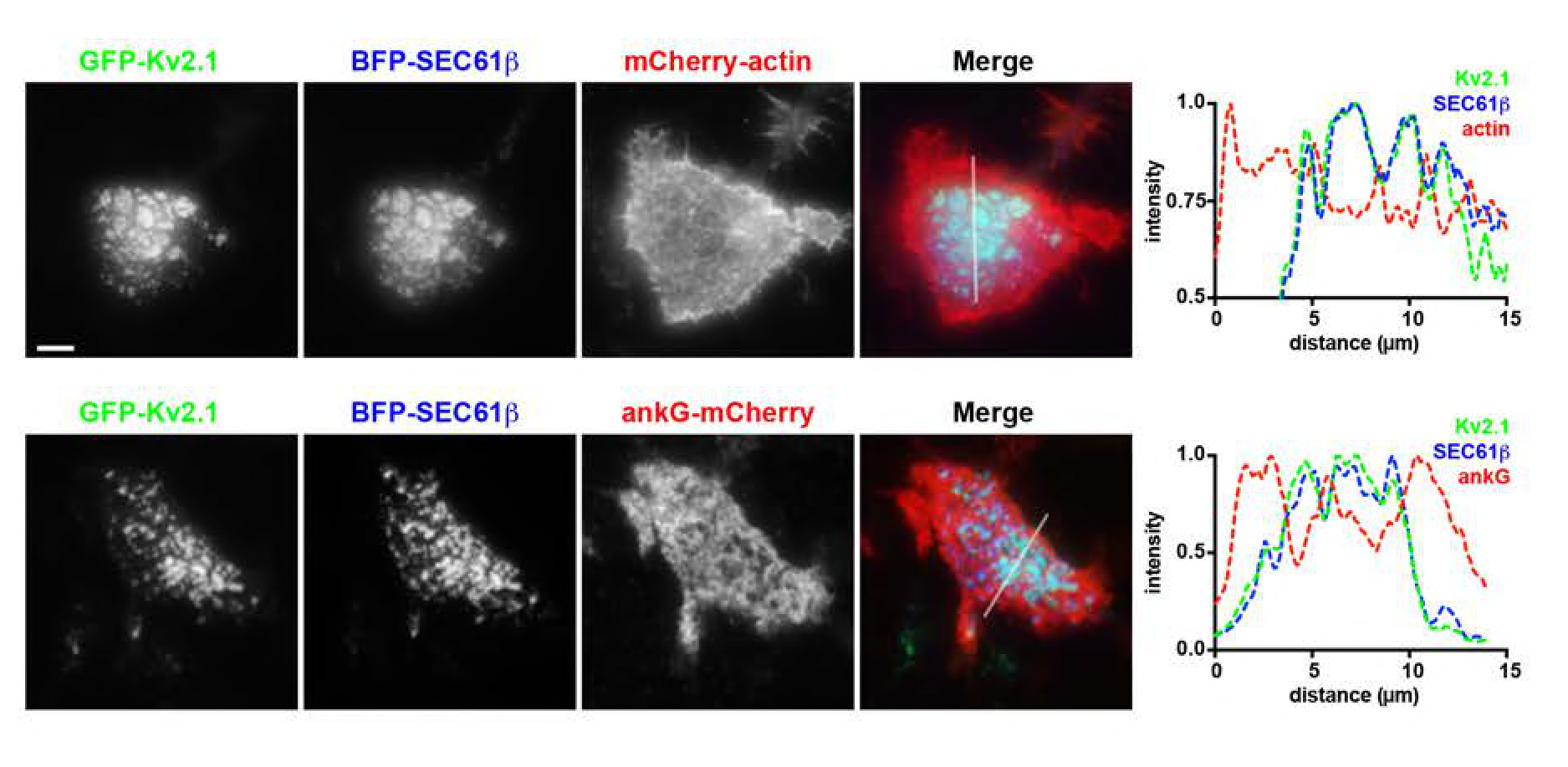
Kv2.1-mediated ER-PM junctions are located at sites depleted in components of the cortical actin cytoskeleton. Top panels. TIRF image of a live HEK293T cell expressing GFP-Kv2.1, BFP-SEC61β, and mCherry-actin. Bottom panels. TIRF image of a live HEK293T cell expressing GFP-Kv2.1, BFP- SEC61β, and ankG-mCherry. Scale bar for GFP-Kv2.1 panel in top row is 5 µm and holds for all panels in set. Panels to the right of each row are the corresponding normalized fluorescence intensity values across the individual line scans depicted by the white line in the merged images.

**Figure 7-figure supplement 1.**
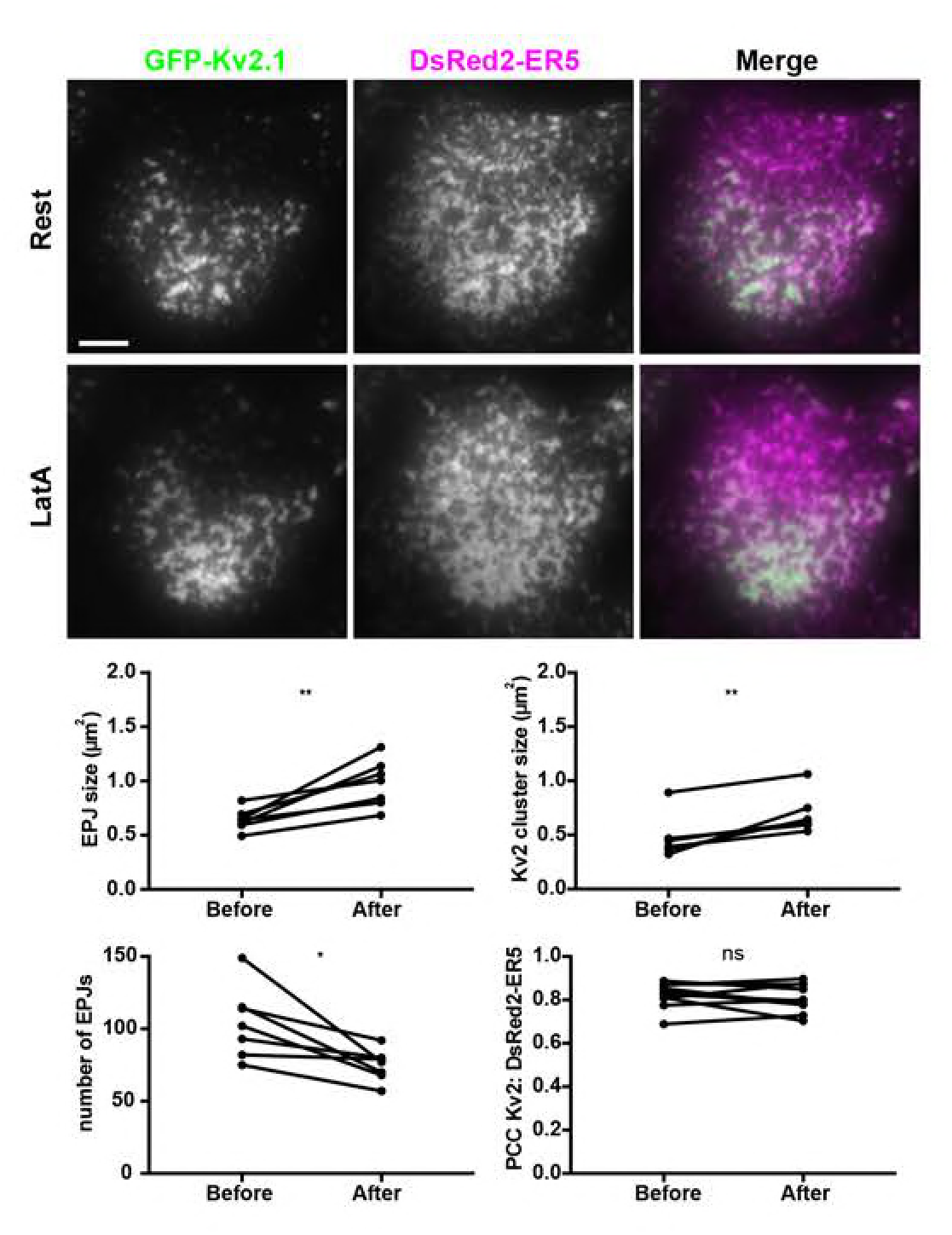
Disrupting the actin cytoskeleton impacts spatial organization of Kv2.1-mediated ER-PM junctions. TIRF image of a live HEK293T cell coexpressing GFP-Kv2.1 and DsRed2-ER5, prior to, and 15 min after, Latrunculin A (LatA) treatment. Scale bar in GFP-Kv2.1 Rest panel is 5 µm and holds for all panels. Graphs show values measured from cells before and after a 15-minute treatment with 10 µM LatA. Top left graph. Mean ER-PM junction (EPJ) size. Top right graph: Mean Kv2.1 cluster size per cell. Bottom left graph. Number of ER-PM junctions (EPJs) per cell. Bottom right graph. PCC values between Kv2.1 and DsRed2-ER5. Bars on all graphs are mean ± SD. See Figure 7-Tables 1-3 for values and statistical analyses.

**Figure 8-figure supplement 1.**
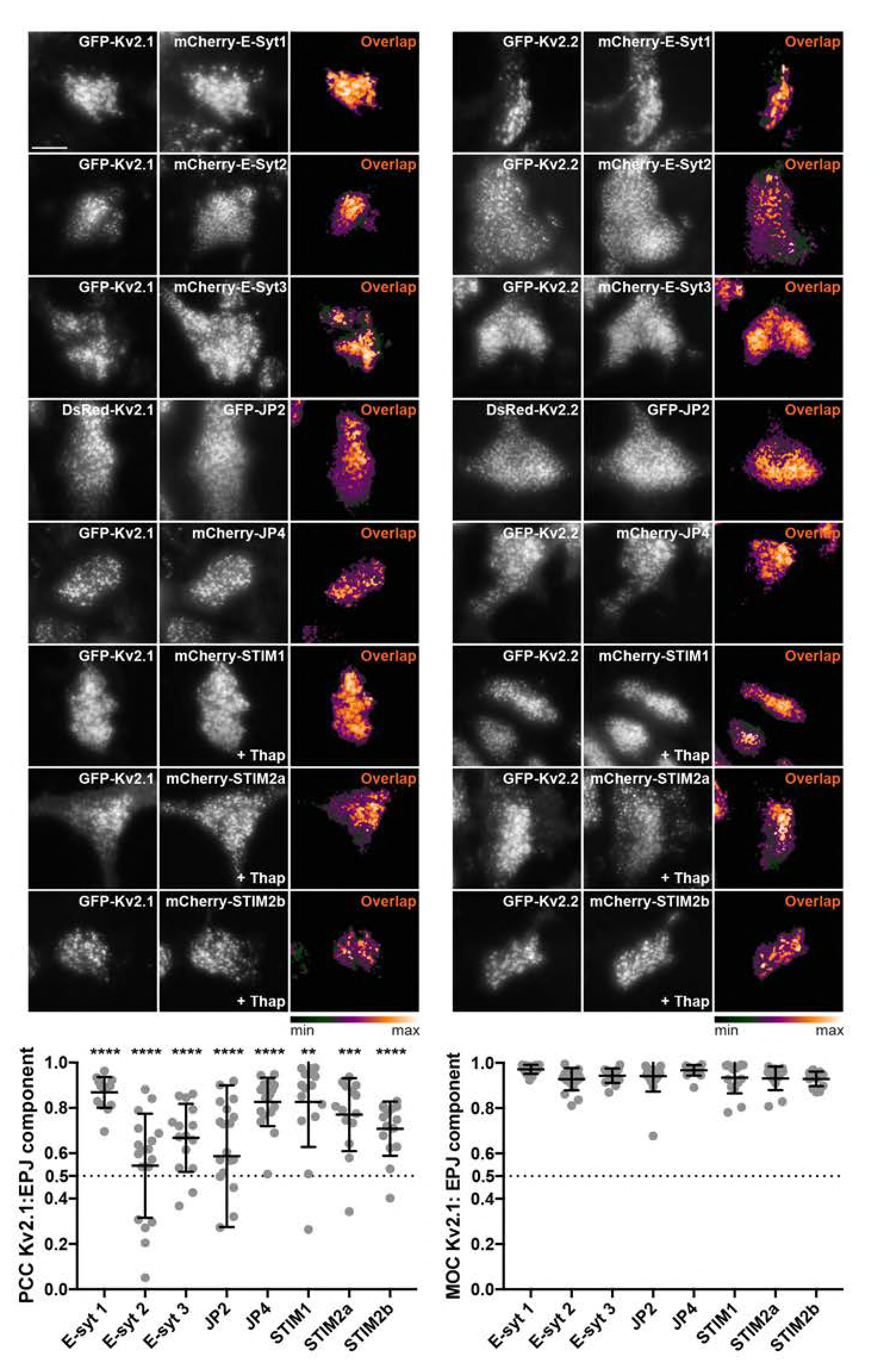
Kv2s colocalize with multiple native components of ER-PM junctions from the E-Syt, JP, and STIM families. TIRF images of live HEK293T cells coexpressing GFP-Kv2.2 or DsRed-Kv2.2 (left panels) or GFP-Kv2.1 or DsRed-Kv2.1 (right panels) and members of the E-Syt, JP and STIM families of ER-localized PM tethers. Heat maps show pixel overlap of Kv2 and ER-PM tether signals. The STIM samples were treated with 2 µM thapsigargin for 5 minutes prior to imaging. Scale bar in top left GFP-Kv2.2 panel is 10 µm and holds for all panels in figure. Graphs show PCC and MOC values of Kv2.1 and ER-PM tether signals. Bars are mean ± SD. See Figure 8-Table 3 for values and statistical analyses.

**Figure 8-figure supplement 2.**
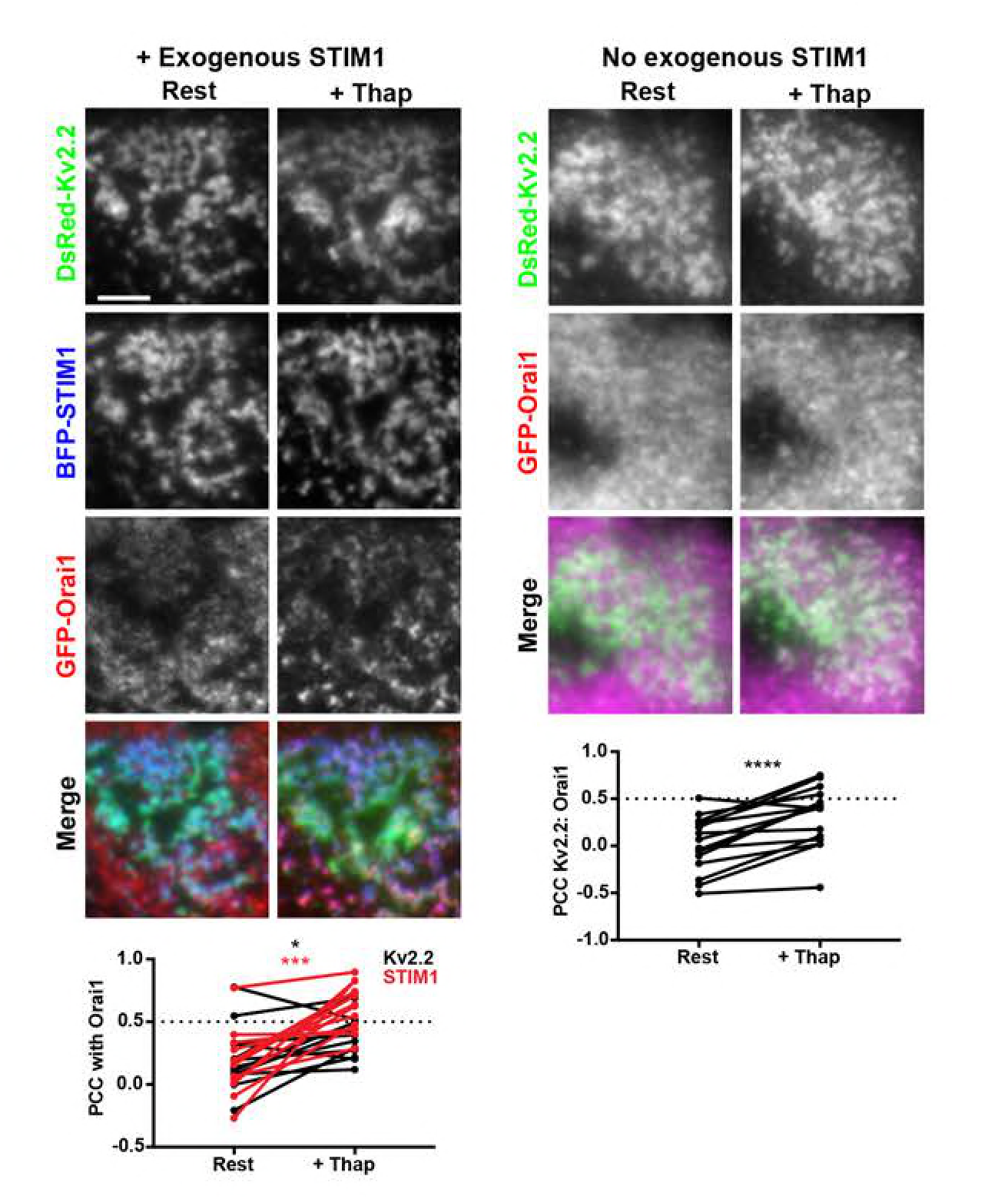
Orai1 translocates to Kv2.2-containing ER-PM junctions in response to store depletion independent of exogenous STIM1 expression. TIRF images of live HEK293T cells coexpressing DsRed-Kv2.2 and GFP-Orai1 with (left panels) and without (right panels) BFP-STIM1 coexpression. For each set the same cell is shown prior to and immediately after 5 min of treatment with 2 µM Thapsigargin. Scale bar in top left DsRed-Kv2.2 panel is 5 µm and holds for all panels in figure. Bottom left graph. PCC values between Orai1 and Kv2.2 (black) or STIM1 (red) measured from cells with BFP-STIM1 coexpression before (Rest) and after (+Thap) Thapsigargin treatment. Bottom right graph. PCC values between Orai1 and Kv2.2 measured from cells without BFP-STIM1 coexpression before (Rest) and after (+Thap) Thapsigargin treatment. Bars on all graphs are mean ± SD. See Figure 8-Table 4 for values and statistical analyses.

**Figure 8-figure supplement 3.**
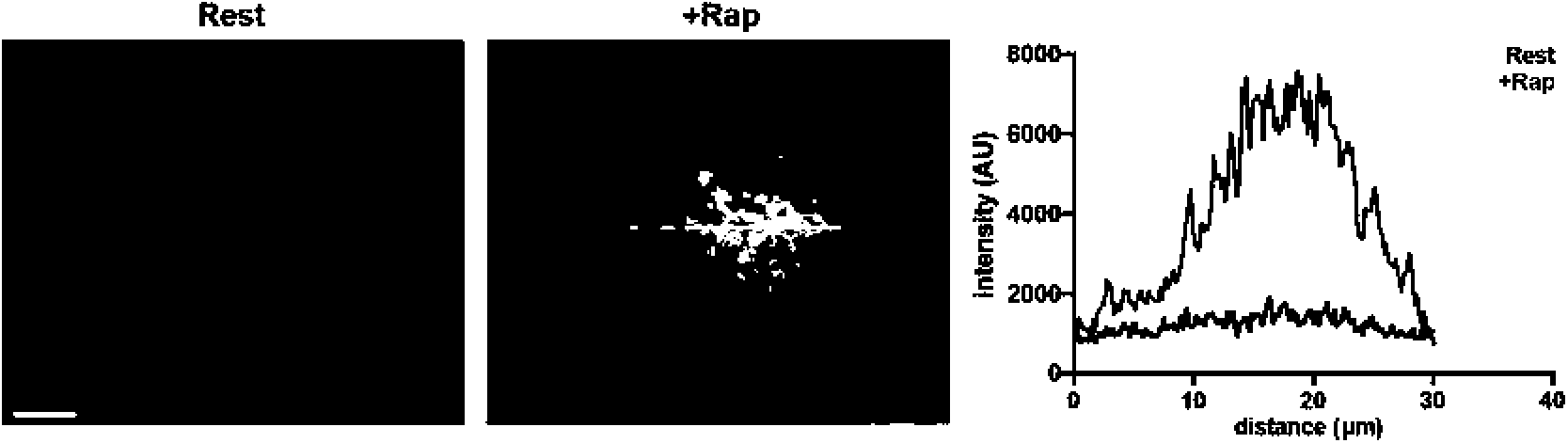
Formation of enhanced ER-PM junctions in HEK293T cells triggered by a rapamycin based heterodimerization strategy. TIRF images of CFP fluorescence in a HEK293T cell coexpressing CFP-CB5-FKBP and lyn11- FRB before (rest) and immediately after treatment with 5 µM rapamycin. Scale bar is 5 µm and holds for all panels. Left graph shows fluorescence intensity of CFP-CB5-FKBP across the individual line scan depicted by the white lines at rest and immediately following treatment with 5 µM rapamycin.

**Figure 8-figure supplement 4.**
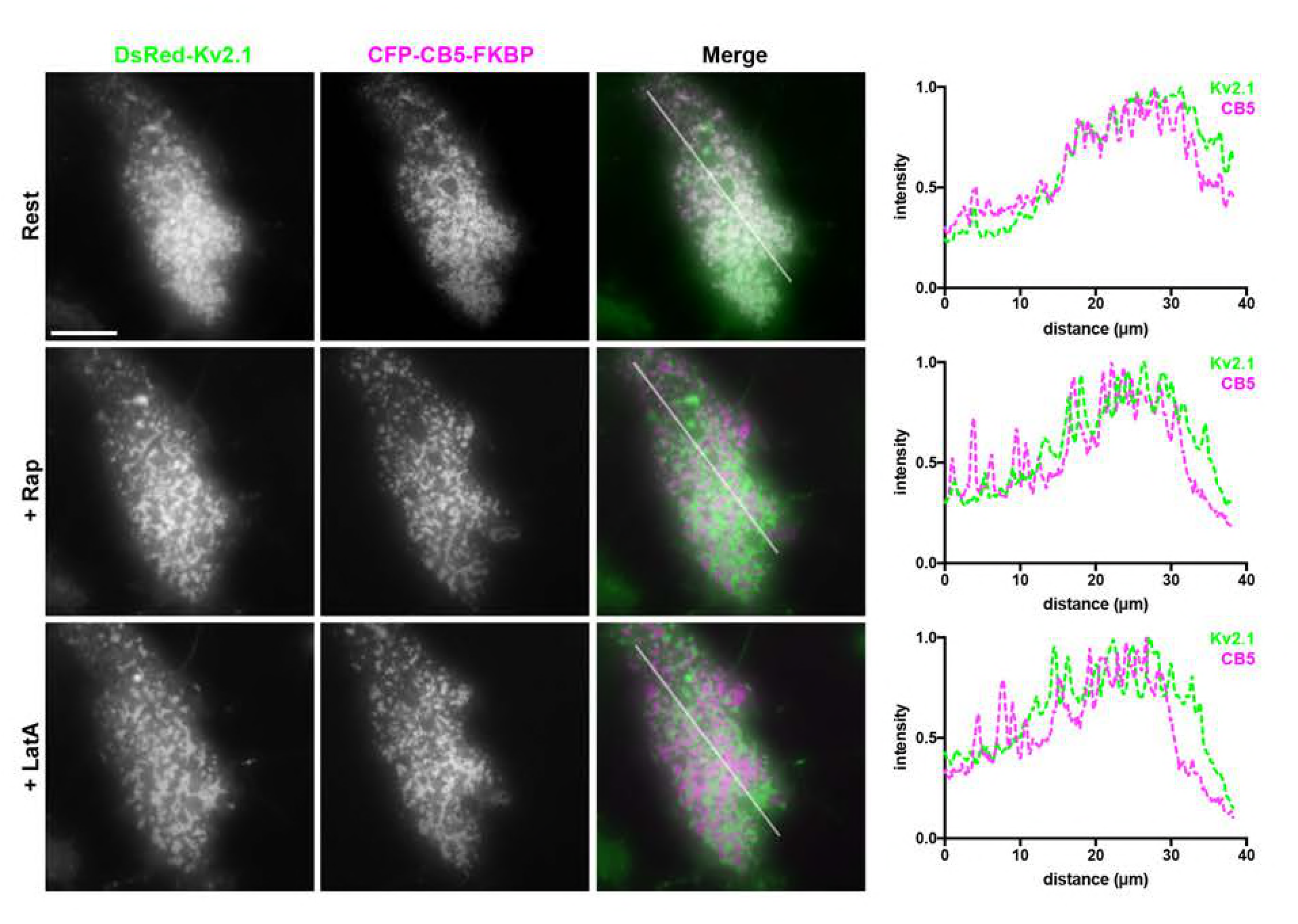
Enhanced ER-PM junctions triggered by a rapamycin based heterodimerization strategy are mutually exclusive with Kv2.1 clusters in HEK293T cells. Left panels. TIRF images of a live HEK293T cell coexpressing DsRed-Kv2.1, CFP-CB5-FKBP, and Lyn11-FRB. Scale bar is 5 µm and holds for all panels. Top row. Prior to rapamycin treatment (rest). Middle row. Same cell immediately following 5 µM rapamycin treatment (+Rap). Bottom row. Same cell after subsequent 15-minute treatment with 10 µM LatA (+LatA). Panels to the right of each row shows the corresponding normalized fluorescence intensity values across the individual line scans depicted by the white line in the merged images.

**Figure 9-figure supplement 1.**
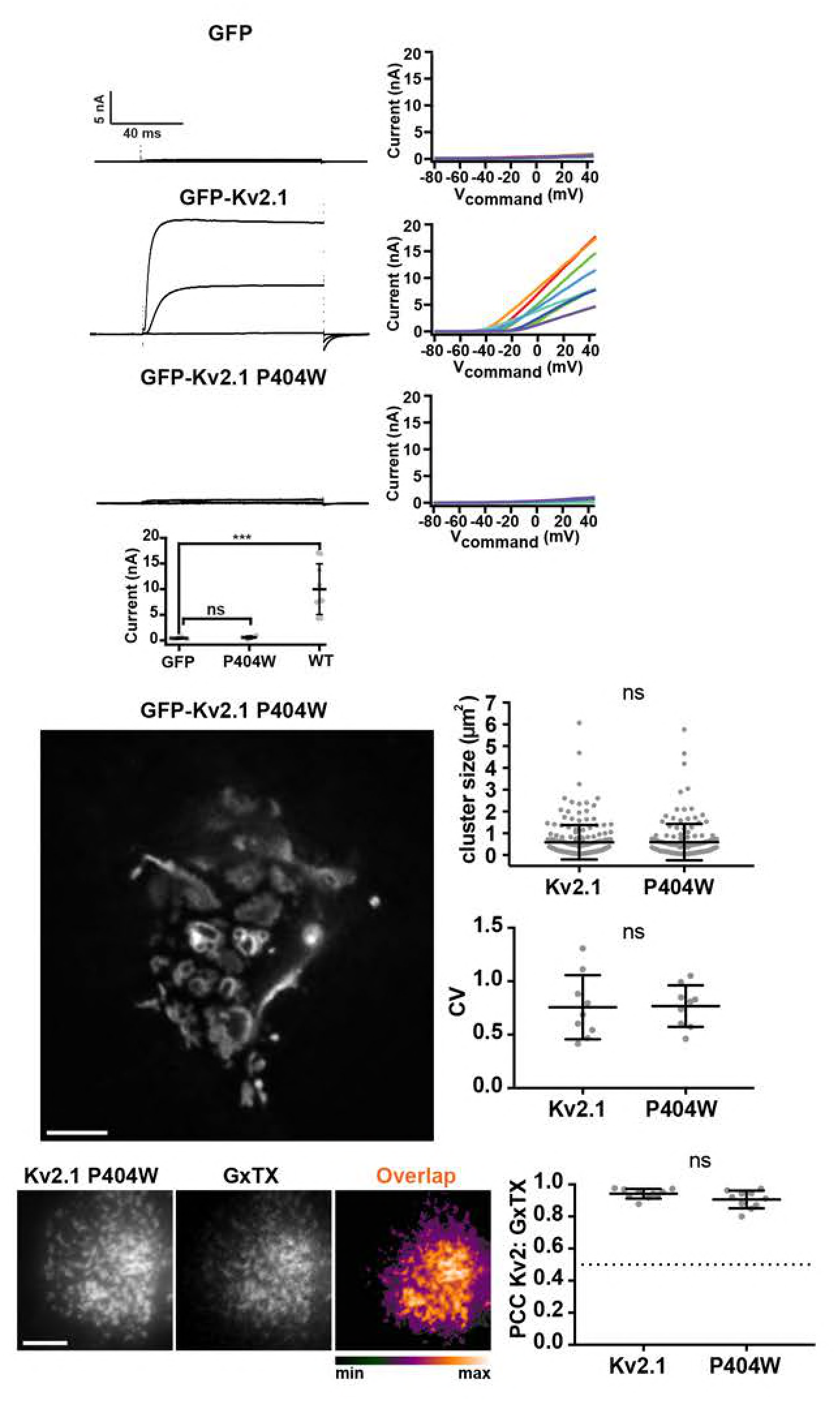
Mutations that eliminate K^+^ conductance do not impact Kv2.1 channel clustering. Top panels show exemplar whole-cell voltage clamp recordings (left) and corresponding graphs of current levels versus command voltage (right) of HEK293T cells expressing GFP-Kv2.1 or GFP-Kv2.1 P404W. Recordings shown are representative responses to 100 ms steps from −100 mV to −40, 0 and +40 mV are shown on left. Note the lack of outward currents in GFP-Kv2.1 P404W recordings. Summary graph shows whole cell current at +40 mV. See Figure 9-Table 1 for values and statistical analyses. Middle panel shows a deconvolved widefield image of a live DIV 7-10 CHN expressing GFP-Kv2.1 P404W. Scale bar is 5 µm. Graphs to the right are measurements of mean cluster size per cell or CV values. Bars are mean ± SD; measured from CHNs transfected with GFP-Kv2.1 or GFP-Kv2.1 P404W. See Figure 9-Tables 2-3 for values and statistical analyses. Bottom panels show TIRF images of live HEK293T cells expressing GFP- Kv2.1 P404W and surface labeled with GxTX-633. Scale bar in Kv2.1 P404W panel is 5 µm and holds for all panels in row. The graph to the right shows comparisons of PCC measurements of Kv2 and GxTX fluorescence from cells expressing GFP-Kv2.1 and/or GFP-Kv2.1 P404W. Dashed line denotes a PCC value of 0.5. Bars are mean ± SD. See Figure 9-Table 4 for values and statistical analyses.

**Figure 10-figure supplement 1.**
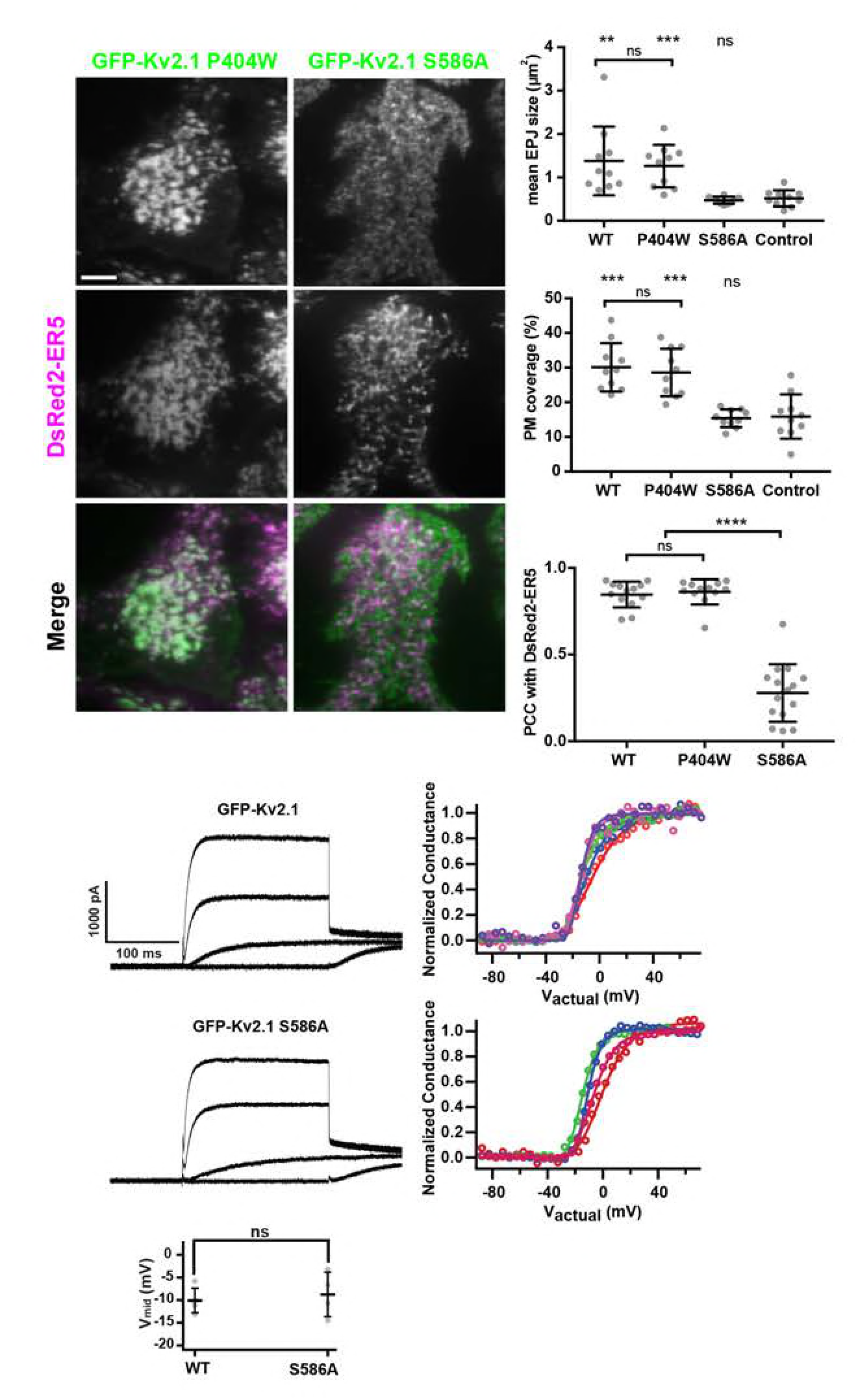
Separation of function point mutations show that clustering, but not conduction, is necessary for Kv2.1-mediated remodeling of ER-PM junctions. Left panels show TIRF images of live HEK293T cells expressing GFP-tagged Kv2.1 mutants and DsRed2-ER5. Scale bar is 5 µm and holds for all panels. Graphs show comparisons from cells expressing wild-type and mutant Kv2.1 isoforms. Top right graph. Mean ER-PM junction (EPJ) size per cell. Middle right graph. Percent PM per cell occupied by cortical ER. Lower right graph. PCC values between DsRed2-ER5 and wild-type and mutant Kv2.1 isoforms. Bars are mean ± SD. See Figure 10-Tables 1-3 for values and statistical analyses. Bottom panels show exemplar whole-cell voltage clamp recordings (left) and graphs of the corresponding normalized conductance-voltage relationship from HEK293T cells expressing GFP-Kv2.1, or GFP-Kv2.1 S586A. Different colors represent data from distinct cells. Recordings shown are representative responses to 200 ms steps from −100 mV to −40, 0 and +40 mV. Bottom graph shows V_mid_ values. Note the lack of effects of the declustering point mutations on the properties of the whole cell currents. Bars are mean ± SD. See Figure 10-Tables 4-5 for values and statistical analyses.

**Figure 10-figure supplement 2.**
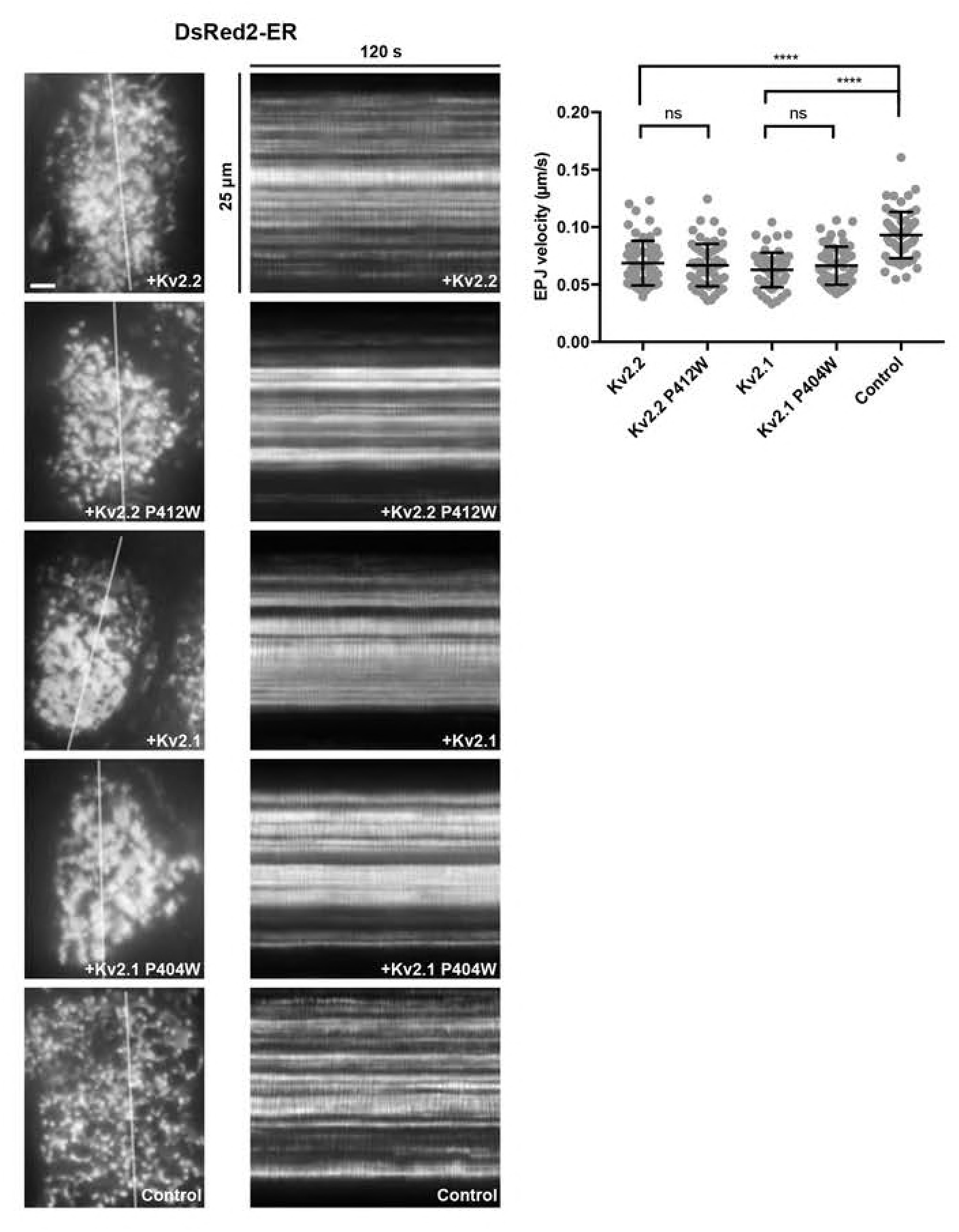
Both wild-type and nonconducting Kv2 channel mutants stabilize ER-PM junctions in HEK293T cells. Left panels are TIRF images of DsRed2-ER5 expressed in live HEK293T cells with and without coexpression of wild-type and mutant Kv2 channel isoforms as labeled. Scale bar is 2.5 µm and holds for all panels. Right panels are kymographs of DsRed2-ER5 mobility from regions indicated by the lines in the adjacent panels. Graph to right shows ER-PM junction (EPJ) velocity (as reported by DsRed2-ER5 in TIRF) as measured from kymographs. Bars are mean ± SD. See Figure 10-Table 6 for values and statistical analyses.

**Figure 11-figure supplement 1.**
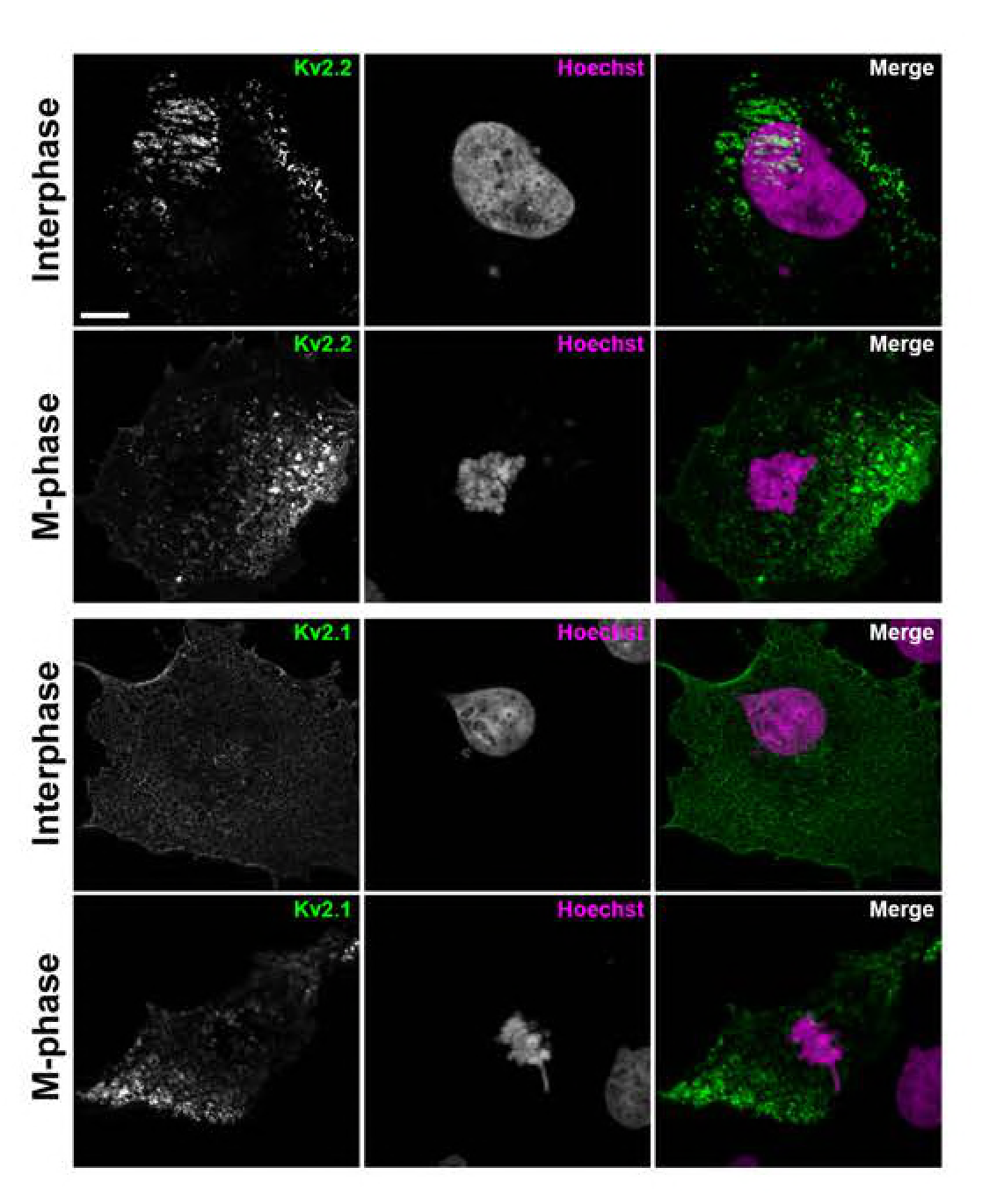
Kv2.2 clusters in COS-1 cells during interphase and M- phase. Single optical sections of fixed interphase (top rows) or M phase (bottom rows) COS-1 cells stained with Hoechst 33258 and expressing Kv2.2 (immunolabeled with mAb N372B/60) or Kv2.1 (immunolabeled with mAb K89/34). Scale bar in Kv2.1 interphase panel is 10 µm and is for all panels. Note chromatin morphologies characteristic of interphase or M phase nuclei as revealed by Hoechst 33258 labeling.

**Movie 1.** Rotating 3D reconstruction of a fixed HEK293T cell expressing GFP-Kv2.2 (left panel, green) and BFP-SEC61β (middle panel, magenta). Merged image is shown in right panel.

**Figure 3-Table 1.**
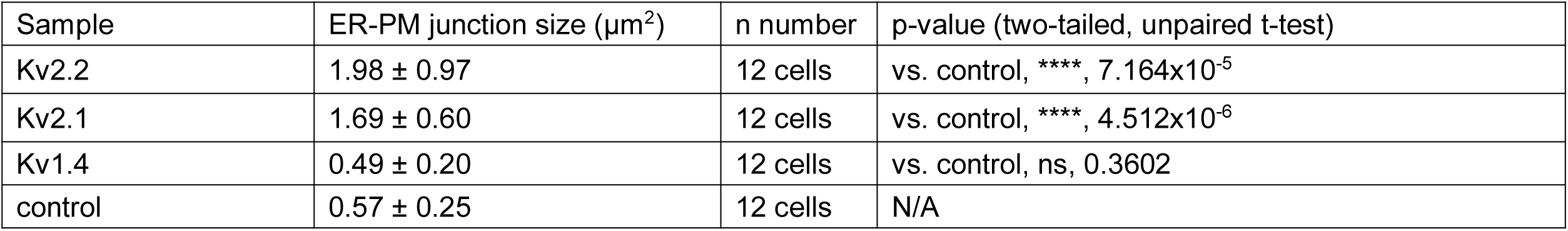
Kv2 channels impact ER-PM junction size.

**Figure 3-Table 2.**
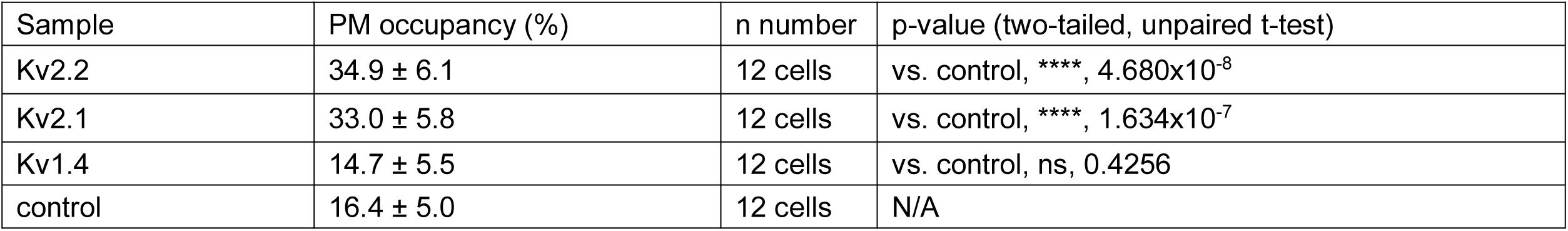
Kv2 channels impact PM occupancy by ER-PM junctions.

**Figure 3-Table 3.**
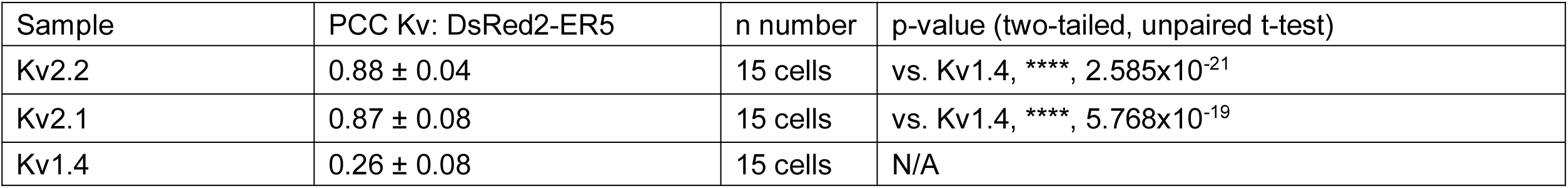
Kv2 channels colocalize with near-PM ER.

**Figure 5-Table 1.**
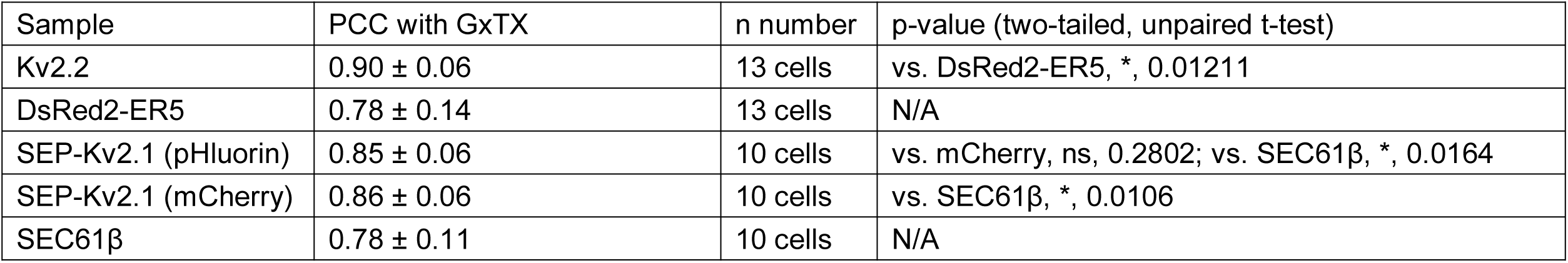
GxTX labeling of cell surface Kv2 channels.

**Figure 6-Table 1.**
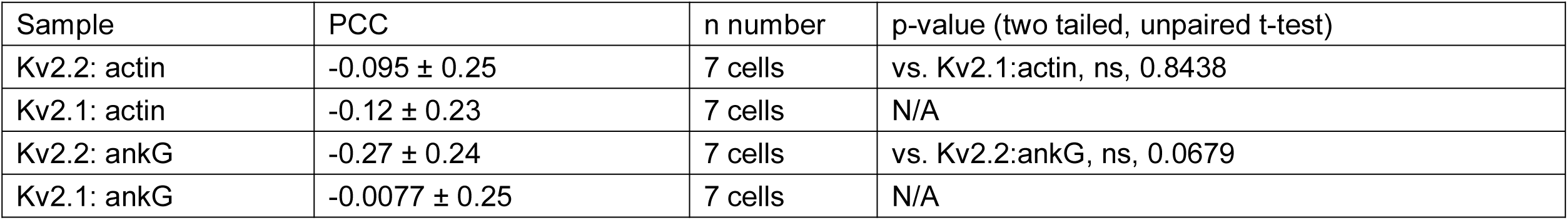
Lack of colocalization of Kv2 channel isoforms with the cortical actin cytoskeleton.

**Figure 7-Table 1.**
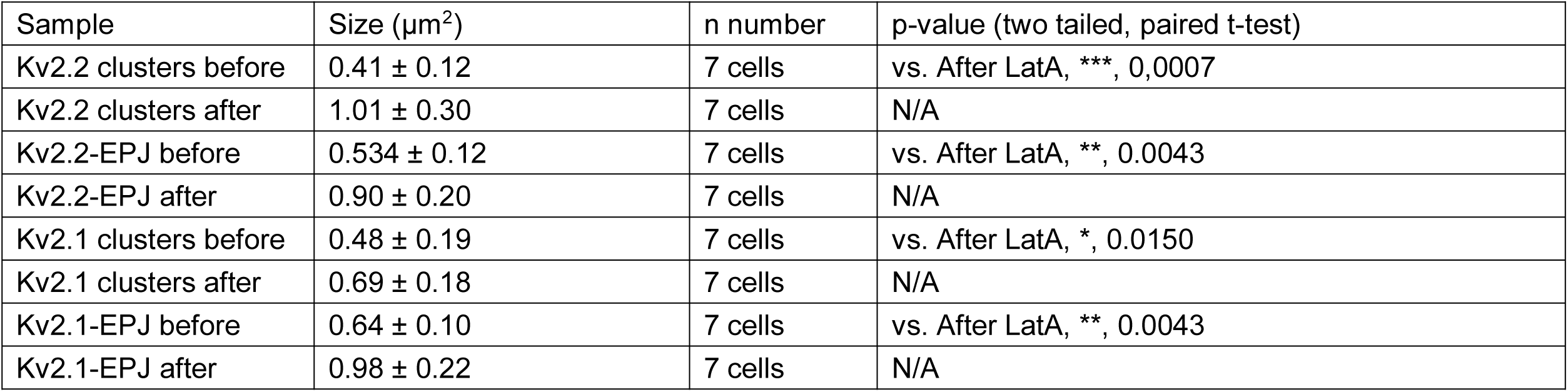
Effects of LatA treatment on Kv2 and ER-PM junction (EPJ) cluster size.

**Figure 7-Table 2.**
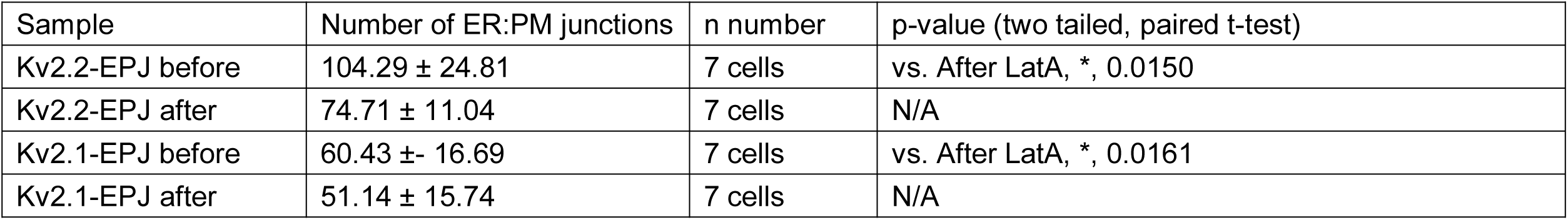
Effects of LatA treatment on number of ER-PM junctions.

**Figure 7-Table 3.**
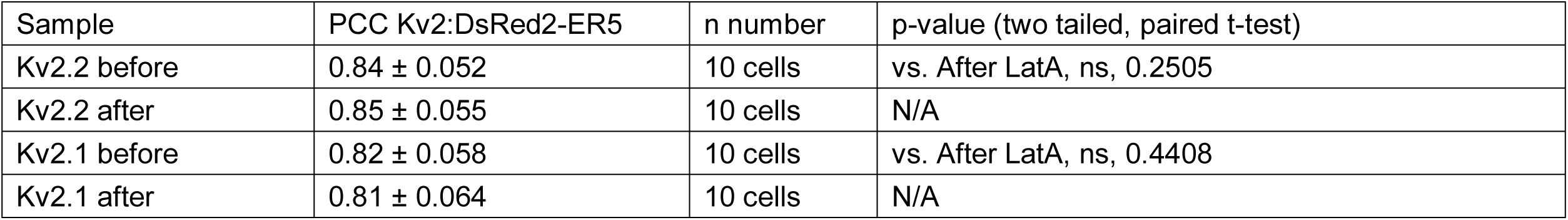
Effects of LatA treatment on Kv2 colocalization with DsRed2-ER5.

**Figure 8-Table 1.**
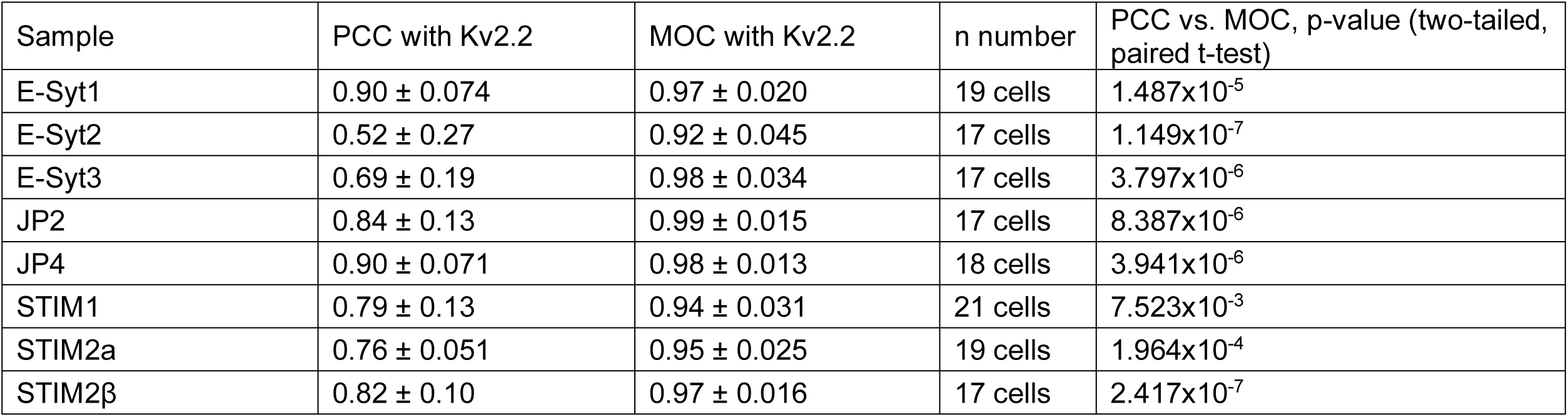
Kv2.2 colocalization with coexpressed ER tethers.

**Figure 8-Table 2.**
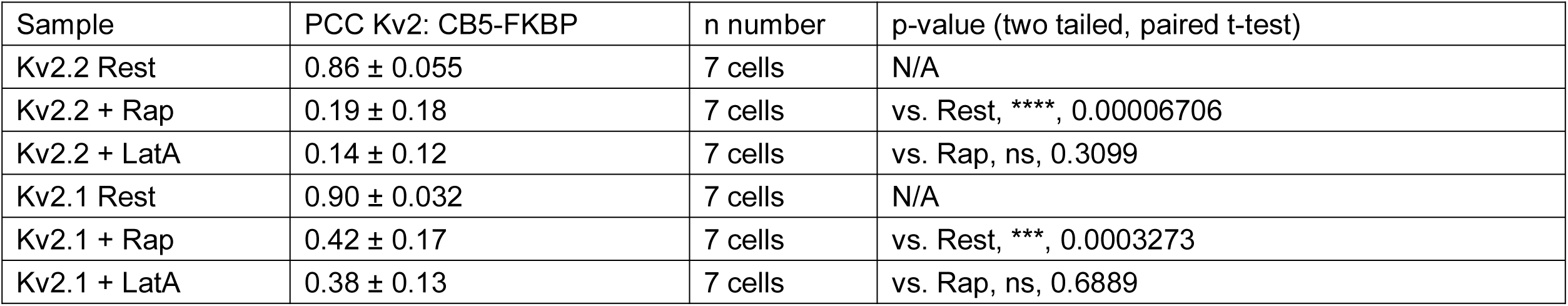
Kv2.2 colocalization with induced ER-PM junctions.

**Figure 8-Table 3.**
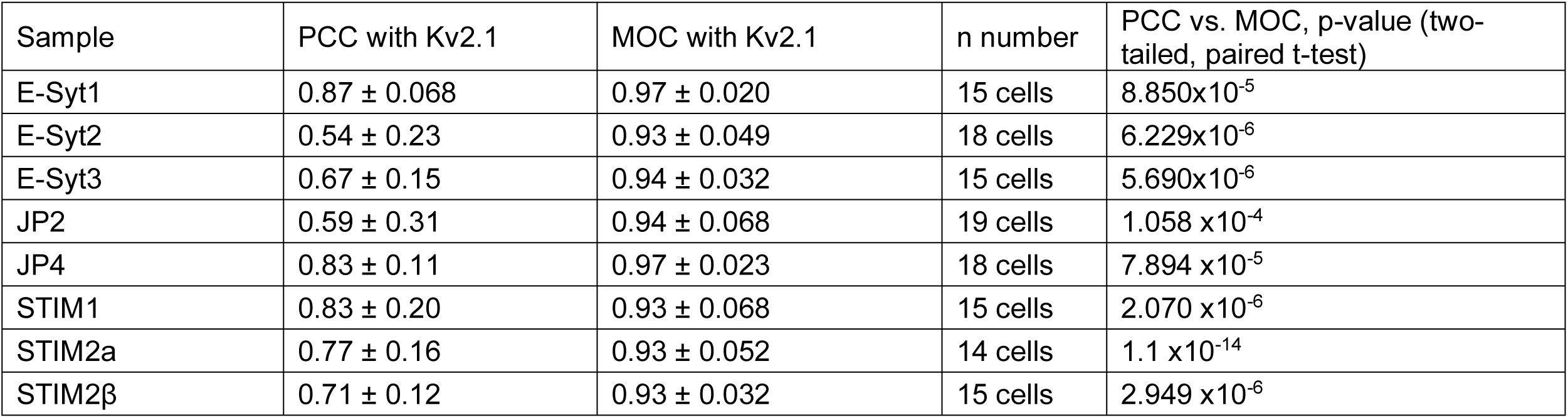
Kv2.1 colocalization with coexpressed ER tethers.

**Figure 8-Table 4.**
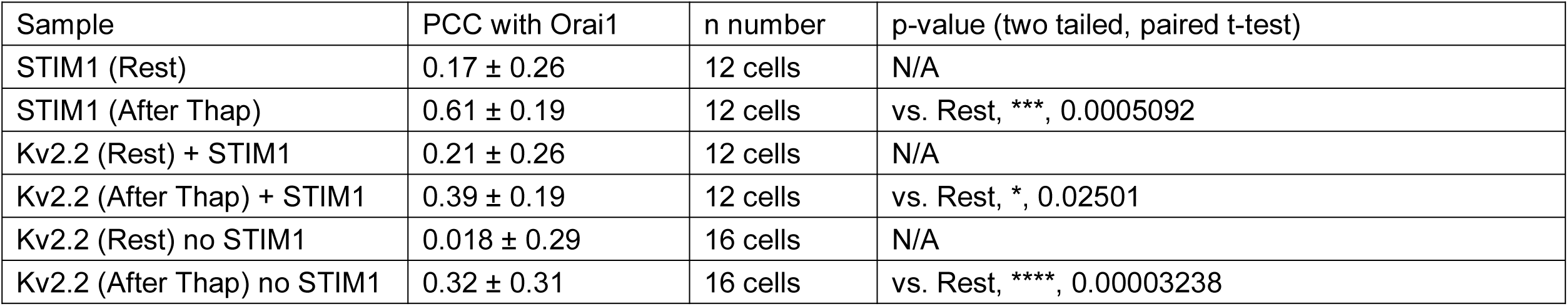
Store depletion results in increased colocalization of Kv2.2 and Orai1.

**Figure 9-Table 1.**
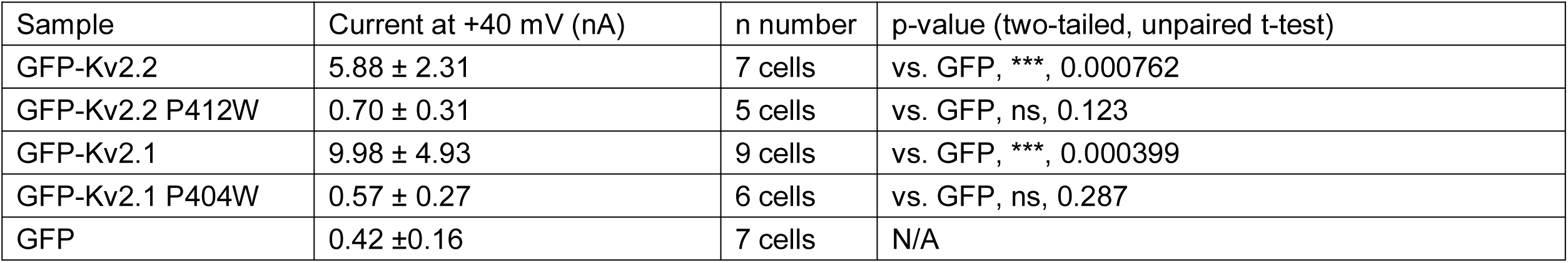
Whole cell current levels of conducting and nonconducting Kv2 channels.

**Figure 9-Table 2.**
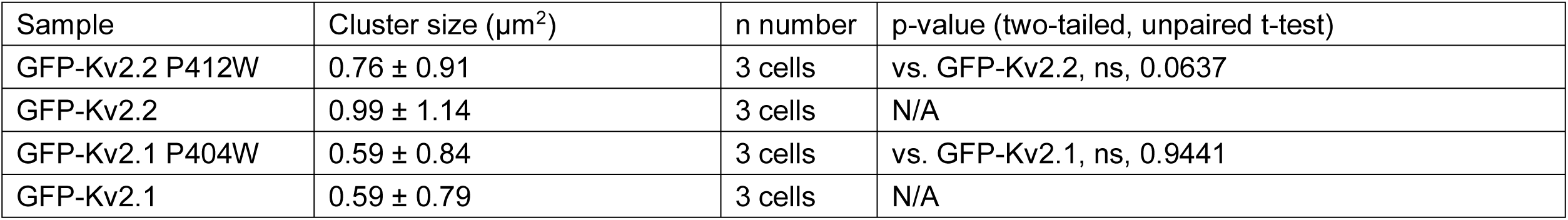
Clustering of conducting and nonconducting Kv2 channels.

**Figure 9-Table 3.**
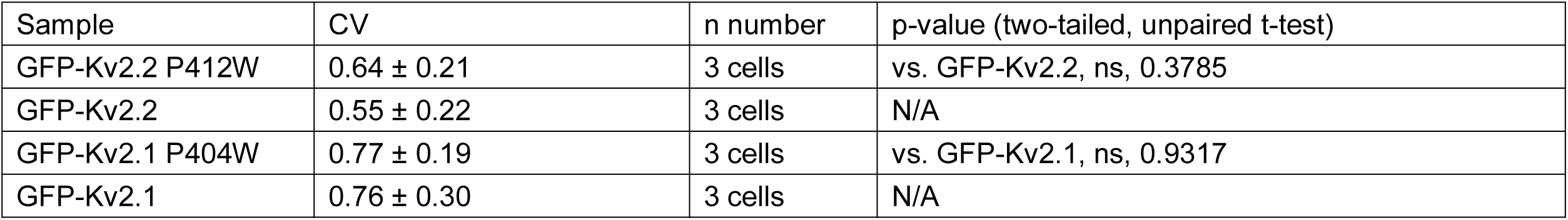
Coefficient of variation of conducting and nonconducting Kv2 channels.

**Figure 9-Table 4.**
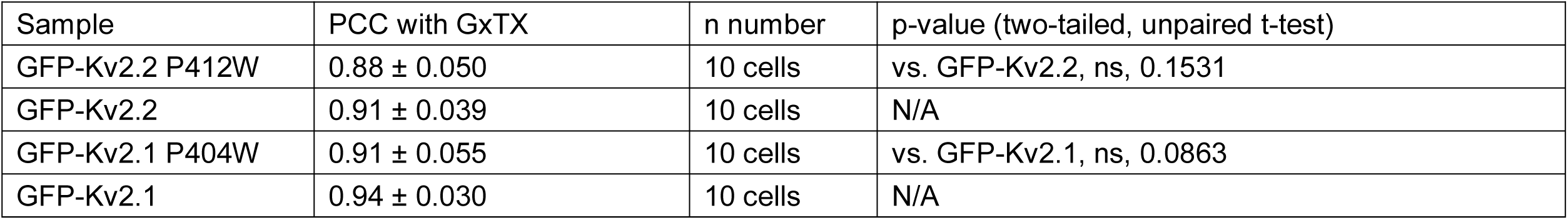
Cell surface expression of conducting and nonconducting Kv2 channels.

**Figure 10-Table 1.**
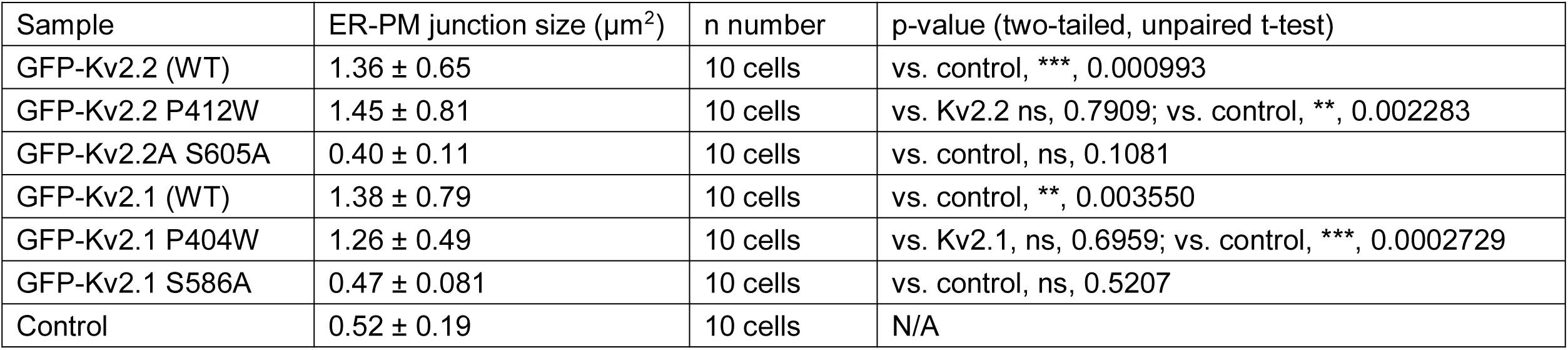
Impact of Kv2 channel isoforms on ER-PM junction size.

**Figure 10-Table 2.**
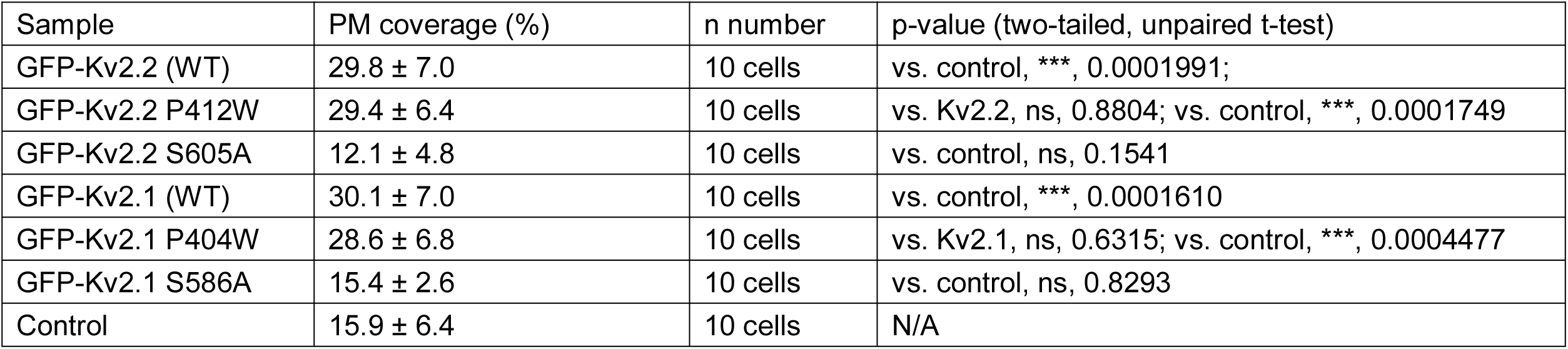
Impact of Kv2 channel isoforms on PM occupancy by ER-PM junctions.

**Figure 10-Table 3.**
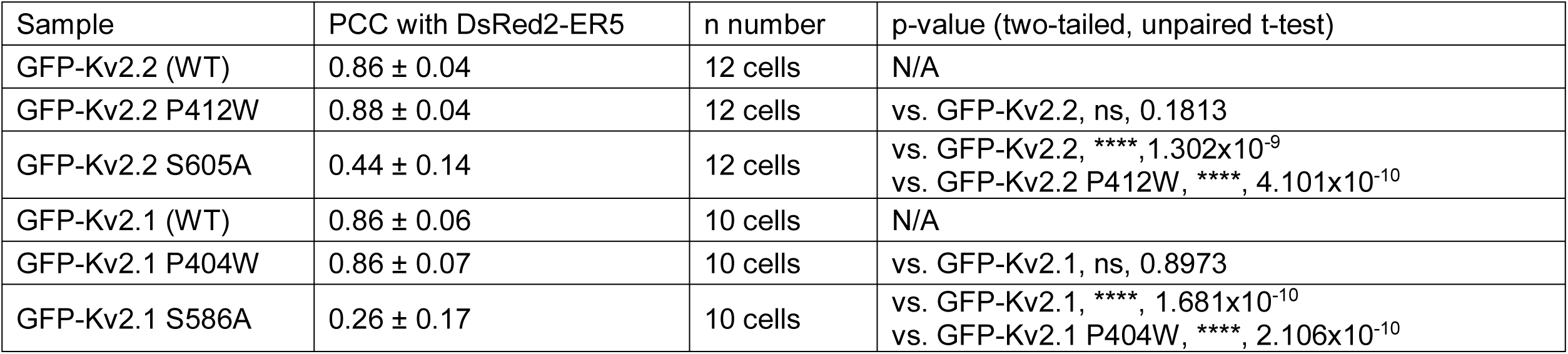
Colocalization of Kv2 channel isoforms with near-PM ER.

**Figure 10-Table 4.**
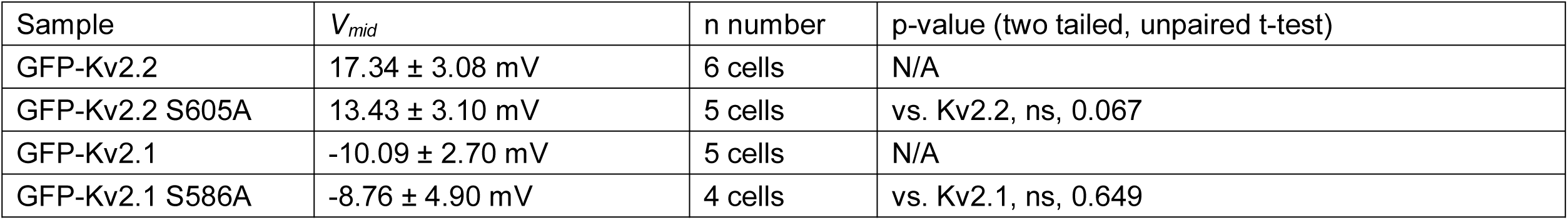
Midpoint of voltage activation of Kv2 channel isoforms.

**Figure 10-Table 5.**
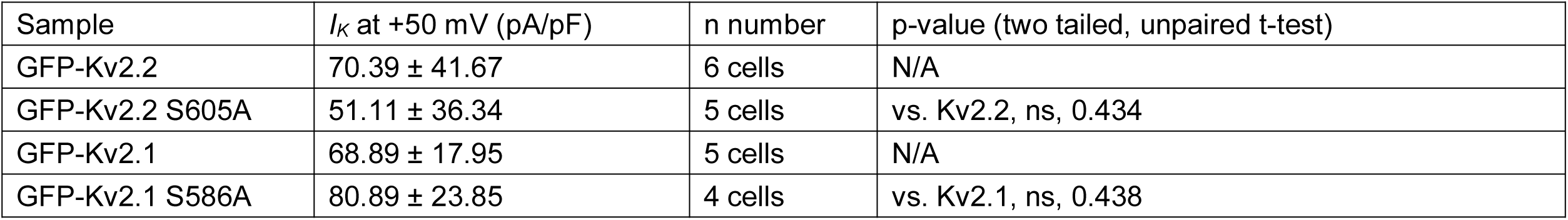
Normalized whole cell current levels of Kv2 channel isoforms.

**Figure 10-Table 6.**
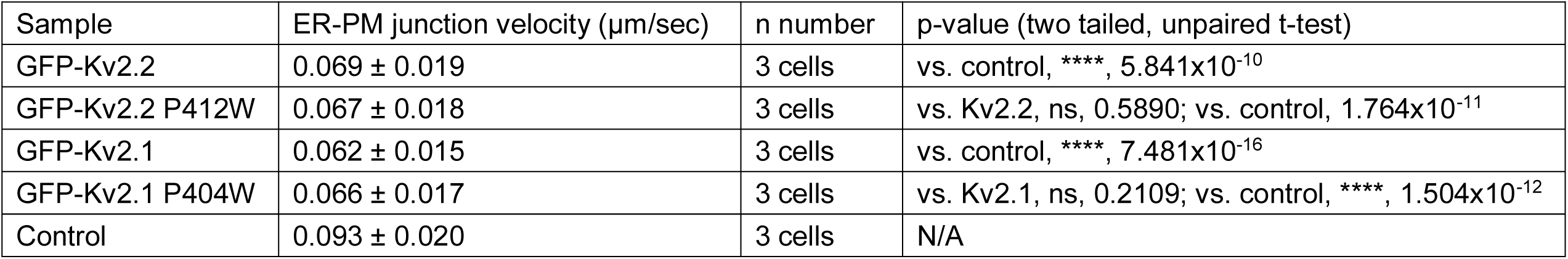
Impact of Kv2 channel isoforms on mobility of near-PM ER.

**Figure 11-Table 1.**
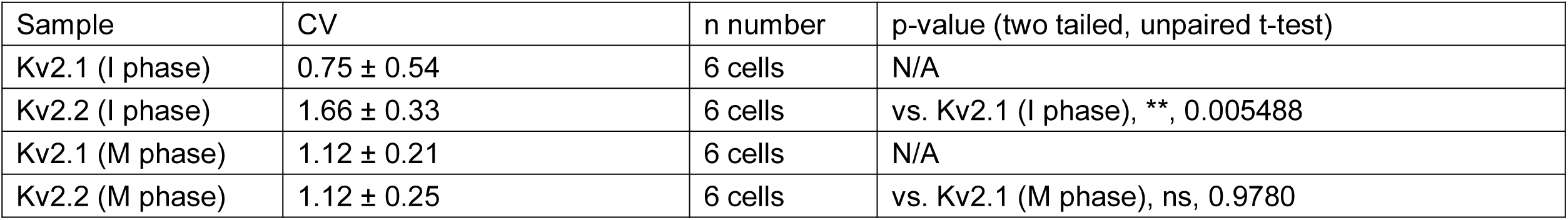
Effects of cell cycle on Kv2 channel clustering.

**Figure 11-Table 2.**
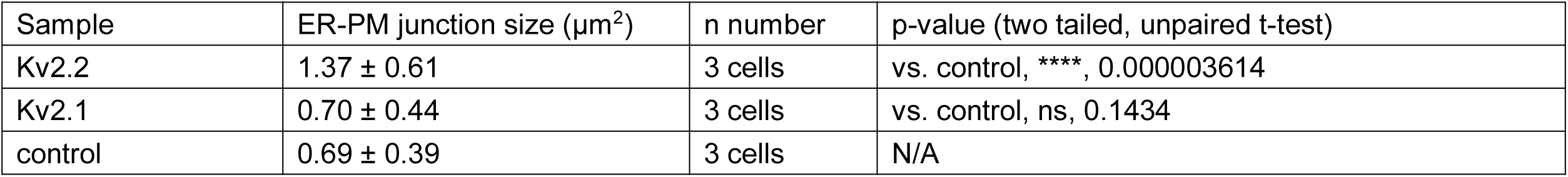
EPJ size in interphase cells.

**Figure 12-Table 1.**
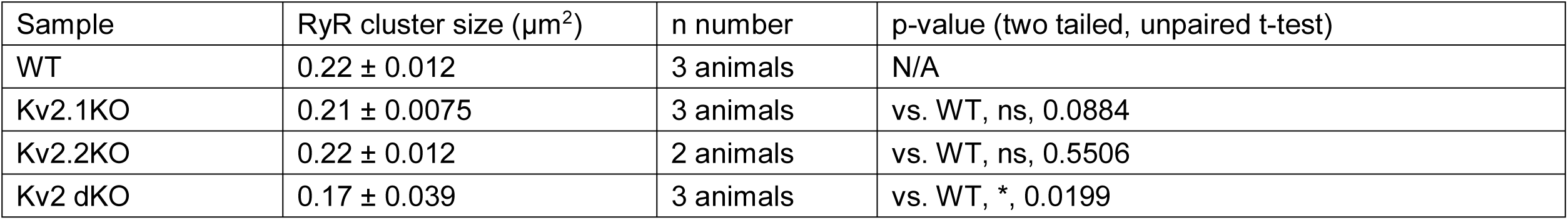
Reduced RyR cluster size in CA1 pyramidal neurons in Kv2 dKO mice.

## Acknowledgments

We thank Drs. Marina Besprozvannaya, Eamonn Dickson, Karl Murray, Jodi Nunnari, and Nicholas Vierra for helpful advice and critical discussions. We also thank Dr. Nicholas Vierra for critical reading of this manuscript. We acknowledge Yongam Lee and Steve Wiler for expert molecular biology technical assistance. We also thank Dr. Michael Paddy at the UC Davis MCB Imaging Facility for expert advice on imaging. We thank Grace Or for help in preparation of cultured hippocampal neurons. GxTX was synthesized at the Molecular Foundry of the Lawrence Berkeley National Laboratory under U. S. Department of Energy Contract DE- AC02-05CH11231. This research was funded by NIH R01 NS042225 and NIH U01 NS090581 to J. S. Trimmer, and NIH R01 NS096317 to J. T. Sack. M. Kirmiz was supported by NIH T32 GM0007377. P. Thapa was supported by American Heart Association postdoctoral fellowship 17POST33670698.

## References

1. Henne WM, Liou J & Emr SD (2015) Molecular mechanisms of inter-organelle ER-PM contact sites. Curr Opin Cell Biol 35:123–130. DOI: 10.1016/j.ceb.2015.05.001.

2. Gallo A, Vannier C & Galli T (2016) Endoplasmic reticulum-plasma membrane associations:structures and functions. Annu Rev Cell Dev Biol 32:279–301. DOI: 10.1146/annurev-cellbio-111315-125024.

3. Saheki Y & De Camilli P (2017) Endoplasmic reticulum-plasma membrane contact sites. Annu Rev Biochem 86:659–684. DOI: 10.1146/annurev-biochem-061516-044932.

4. Chang CL, Chen YJ & Liou J (2017) ER-plasma membrane junctions: Why and how do we study them? Biochim Biophys Acta 1864:1494–1506. DOI: 10.1016/j.bbamcr.2017.05.018.

5. Dickson EJ (2017) Endoplasmic reticulum-plasma membrane contacts regulate cellular excitability. Adv Exp Med Biol 997:95–109. DOI: 10.1007/978-981-10-4567-7_7.

6. Balla T (2017) Ca(2+) and lipid signals hold hands at endoplasmic reticulum-plasma membrane contact sites. J Physiol. DOI: 10.1113/jp274957.

7. Saheki Y & De Camilli P (2017) The extended-synaptotagmins. Biochim Biophys Acta 1864:1490–1493. DOI: 10.1016/j.bbamcr.2017.03.013.

8. Takeshima H, Hoshijima M & Song LS (2015) Ca(2)(+) microdomains organized by junctophilins. Cell Calcium 58:349–356. DOI: 10.1016/j.ceca.2015.01.007.

9. Prakriya M & Lewis RS (2015) Store-operated calcium channels. Physiological reviews 95:1383–1436. DOI: 10.1152/physrev.00020.2014.

10. Carrasco S & Meyer T (2011) STIM proteins and the endoplasmic reticulum-plasma membrane junctions. Annu Rev Biochem 80:973–1000. DOI: 10.1146/annurev-biochem-061609-165311.

11. Nishi M, Sakagami H, Komazaki S, Kondo H & Takeshima H (2003) Coexpression of junctophilin type 3 and type 4 in brain. Brain Res Mol Brain Res 118:102–110.

12. Min SW, Chang WP & Sudhof TC (2007) E-Syts, a family of membranous Ca2+-sensor proteins with multiple C2 domains. Proc Natl Acad Sci U S A 104:3823–3828. DOI: 10.1073/pnas.0611725104.

13. Moccia F, Zuccolo E, Soda T, Tanzi F, Guerra G, Mapelli L, Lodola F & D’Angelo E (2015) Stim and Orai proteins in neuronal Ca(2+) signaling and excitability. Front Cell Neurosci 9:153. DOI: 10.3389/fncel.2015.00153.

14. Rosenbluth J (1962) Subsurface cisterns and their relationship to the neuronal plasma membrane. J Cell Biol 13:405–421.

15. Henkart M, Landis DM & Reese TS (1976) Similarity of junctions between plasma membranes and endoplasmic reticulum in muscle and neurons. J Cell Biol 70:338–347.

16. Wu Y, Whiteus C, Xu CS, Hayworth KJ, Weinberg RJ, Hess HF & De Camilli P (2017) Contacts between the endoplasmic reticulum and other membranes in neurons. Proc Natl Acad Sci U S A 114:E4859–E4867. DOI: 10.1073/pnas.1701078114.

17. Trimmer JS (2015) Subcellular localization of K+ channels in mammalian brain neurons: remarkable precision in the midst of extraordinary complexity. Neuron 85:238–256. DOI: 10.1016/j.neuron.2014.12.042.

18. Johnston J, Griffin SJ, Baker C, Skrzypiec A, Chernova T & Forsythe ID (2008) Initial segment Kv2.2 channels mediate a slow delayed rectifier and maintain high frequency action potential firing in medial nucleus of the trapezoid body neurons. J Physiol 586:3493–3509. DOI: jphysiol.2008.153734 [pii]10.1113/jphysiol.2008.153734.

19. Guan D, Tkatch T, Surmeier DJ, Armstrong WE & Foehring RC (2007) Kv2 subunits underlie slowly inactivating potassium current in rat neocortical pyramidal neurons. J Physiol 581:941–960.

20. Du J, Haak LL, Phillips-Tansey E, Russell JT & McBain CJ (2000) Frequency-dependent regulation of rat hippocampal somato-dendritic excitability by the K+ channel subunit Kv2.1. J Physiol 522:19–31. DOI: PHY_0063[pii].

21. Malin SA & Nerbonne JM (2002) Delayed rectifier K+ currents, IK, are encoded by Kv2 alpha-subunits and regulate tonic firing in mammalian sympathetic neurons. J Neurosci 22:10094–10105.

22. Liu PW & Bean BP (2014) Kv2 channel regulation of action potential repolarization and firing patterns in superior cervical ganglion neurons and hippocampal CA1 pyramidal neurons. J Neurosci 34:4991–5002. DOI: 10.1523/JNEUROSCI.1925-13.2014.

23. Palacio S, Chevaleyre V, Brann DH, Murray KD, Piskorowski RA & Trimmer JS (2017) Heterogeneity in Kv2 channel expression shapes action potential characteristics and firing patterns in CA1 versus CA2 hippocampal pyramidal neurons. eNeuro 4. DOI: 10.1523/ENEURO.0267-17.2017.

24. Honigsperger C, Nigro MJ & Storm JF (2017) Physiological roles of Kv2 channels in entorhinal cortex layer II stellate cells revealed by Guangxitoxin-1E. J Physiol 595:739–757. DOI: 10.1113/jp273024.

25. Pathak D, Guan D & Foehring RC (2016) Roles of specific Kv channel types in repolarization of the action potential in genetically identified subclasses of pyramidal neurons in mouse neocortex. J Neurophysiol 115:2317–2329. DOI: 10.1152/jn.01028.2015.

26. Torkamani A, Bersell K, Jorge BS, Bjork RL, Jr., Friedman JR, Bloss CS, Cohen J, Gupta S, Naidu S, Vanoye CG, George AL, Jr. & Kearney JA (2014) De novo KCNB1 mutations in epileptic encephalopathy. Ann Neurol 76:529–540. DOI: 10.1002/ana.24263.

27. Saitsu H, Akita T, Tohyama J, Goldberg-Stern H, Kobayashi Y, Cohen R, Kato M, Ohba C, Miyatake S, Tsurusaki Y, Nakashima M, Miyake N, Fukuda A & Matsumoto N (2015) De novo KCNB1 mutations in infantile epilepsy inhibit repetitive neuronal firing. Sci Rep 5:15199. DOI: 10.1038/srep15199.

28. Thiffault I, Speca DJ, Austin DC, Cobb MM, Eum KS, Safina NP, Grote L, Farrow EG, Miller N, Soden S, Kingsmore SF, Trimmer JS, Saunders CJ & Sack JT (2015) A novel epileptic encephalopathy mutation in KCNB1 disrupts Kv2.1 ion selectivity, expression, and localization. J Gen Physiol 146:399–410. DOI: 10.1085/jgp.201511444.

29. de Kovel CG, Brilstra EH, van Kempen MJ, Van’t Slot R, Nijman IJ, Afawi Z, De Jonghe P, Djemie T, Guerrini R, Hardies K, Helbig I, Hendrickx R, Kanaan M, Kramer U, Lehesjoki AE, Lemke JR, Marini C, Mei D, Moller RS, Pendziwiat M, Stamberger H, Suls A, Weckhuysen S, Euro ERESC & Koeleman BP (2016) Targeted sequencing of 351 candidate genes for epileptic encephalopathy in a large cohort of patients. Mol Genet Genomic Med 4:568–580. DOI: 10.1002/mgg3.235.

30. Jacobson DA, Kuznetsov A, Lopez JP, Kash S, Ammala CE & Philipson LH (2007) Kv2.1 ablation alters glucose-induced islet electrical activity, enhancing insulin secretion. Cell Metab 6:229–235.

31. Li XN, Herrington J, Petrov A, Ge L, Eiermann G, Xiong Y, Jensen MV, Hohmeier HE, Newgard CB, Garcia ML, Wagner M, Zhang BB, Thornberry NA, Howard AD, Kaczorowski GJ & Zhou YP (2013) The role of voltage-gated potassium channels Kv2.1 and Kv2.2 in the regulation of insulin and somatostatin release from pancreatic islets. J Pharmacol Exp Ther 344:407–416. DOI: 10.1124/jpet.112.199083.

32. Patel AJ, Lazdunski M & Honore E (1997) Kv2.1/Kv9.3, a novel ATP-dependent delayed930 rectifier K+ channel in oxygen-sensitive pulmonary artery myocytes. EMBO J 16:6615–6625.

33. Schmalz F, Kinsella J, Koh SD, Vogalis F, Schneider A, Flynn ER, Kenyon JL & Horowitz B (1998) Molecular identification of a component of delayed rectifier current in gastrointestinal smooth muscles. Am J Physiol 274:G901–911.

34. Trimmer JS (1991) Immunological identification and characterization of a delayed rectifier K+ channel polypeptide in rat brain. Proc Natl Acad Sci U S A 88:10764–10768.

35. Scannevin RH, Murakoshi H, Rhodes KJ & Trimmer JS (1996) Identification of a cytoplasmic domain important in the polarized expression and clustering of the Kv2.1 K+ channel. J Cell Biol 135:1619–1632.

36. Murakoshi H & Trimmer JS (1999) Identification of the Kv2.1 K+ channel as a major component of the delayed rectifier K+ current in rat hippocampal neurons. J Neurosci 19:1728–1735.

37. Lim ST, Antonucci DE, Scannevin RH & Trimmer JS (2000) A novel targeting signal for proximal clustering of the Kv2.1 K+ channel in hippocampal neurons. Neuron 25:385–397. DOI: S0896-6273(00)80902-2 [pii].

38. Mandikian D, Bocksteins E, Parajuli LK, Bishop HI, Cerda O, Shigemoto R & Trimmer JS (2014) Cell type-specific spatial and functional coupling between mammalian brain Kv2.1 K(+) channels and ryanodine receptors. J Comp Neurol 522:3555–3574. DOI: 10.1002/cne.23641.

39. Bishop HI, Guan D, Bocksteins E, Parajuli LK, Murray KD, Cobb MM, Misonou H, Zito K, Foehring RC & Trimmer JS (2015) Distinct cell- and layer-specific expression patterns and independent regulation of Kv2 channel subtypes in cortical pyramidal neurons. J Neurosci 35:14922–14942. DOI: 10.1523/JNEUROSCI.1897-15.2015.

40. Du J, Tao-Cheng JH, Zerfas P & McBain CJ (1998) The K+ channel, Kv2.1, is apposed to astrocytic processes and is associated with inhibitory postsynaptic membranes in hippocampal and cortical principal neurons and inhibitory interneurons. Neuroscience 84:37–48.

41. Bishop HI, Cobb MM, Kirmiz M, Parajuli LK, Mandikian D, Philp AM, Melnik M, Kuja-Panula J, Rauvala H, Shigemoto R, Murray KD & Trimmer JS (2018) Kv2 ion channels determine the expression and localization of the associated AMIGO-1 cell adhesion molecule in adult brain neurons. Front Mol Neurosci 11:1. DOI: 10.3389/fnmol.2018.00001.

42. Kihira Y, Hermanstyne TO & Misonou H (2010) Formation of heteromeric Kv2 channels in mammalian brain neurons. J Biol Chem 285:15048–15055. DOI: M109.074260 [pii]10.1074/jbc.M109.074260.

43. O’Connell KM & Tamkun MM (2005) Targeting of voltage-gated potassium channel isoforms to distinct cell surface microdomains. Journal of cell science 118:2155–2166.

44. O’Connell KM, Rolig AS, Whitesell JD & Tamkun MM (2006) Kv2.1 potassium channels are retained within dynamic cell surface microdomains that are defined by a perimeter fence. J Neurosci 26:9609–9618.

45. Mohapatra DP & Trimmer JS (2006) The Kv2.1 C terminus can autonomously transfer Kv2.1-like phosphorylation-dependent localization, voltage-dependent gating, and muscarinic modulation to diverse Kv channels. J Neurosci 26:685–695. DOI: 26/2/685 [pii]10.1523/JNEUROSCI.4620-05.2006.

46. Tamkun MM, O’Connell K M & Rolig AS (2007) A cytoskeletal-based perimeter fence selectively corrals a sub-population of cell surface Kv2.1 channels. Journal of cell science 120:2413–2423.

47. Cobb MM, Austin DC, Sack JT & Trimmer JS (2015) Cell cycle-dependent changes in localization and phosphorylation of the plasma membrane Kv2.1 K+ channel impact endoplasmic reticulum membrane contact sites in COS-1 cells. J Biol Chem 290:29189–29201. DOI: 10.1074/jbc.M115.690198.

48. Antonucci DE, Lim ST, Vassanelli S & Trimmer JS (2001) Dynamic localization and clustering of dendritic Kv2.1 voltage-dependent potassium channels in developing hippocampal neurons. Neuroscience 108:69–81.

49. Franzini-Armstrong C & Jorgensen AO (1994) Structure and development of E-C coupling units in skeletal muscle. Annu Rev Physiol 56:509–534. DOI: 10.1146/annurev.ph.56.030194.002453.

50. Sun XH, Protasi F, Takahashi M, Takeshima H, Ferguson DG & Franzini-Armstrong C (1995) Molecular architecture of membranes involved in excitation-contraction coupling of cardiac muscle. J Cell Biol 129:659–671.

51. Fox PD, Haberkorn CJ, Akin EJ, Seel PJ, Krapf D & Tamkun MM (2015) Induction of stable ER-plasma-membrane junctions by Kv2.1 potassium channels. Journal of cell science 128:2096–2105. DOI: 10.1242/jcs.166009.

52. Misonou H, Mohapatra DP, Park EW, Leung V, Zhen D, Misonou K, Anderson AE & Trimmer JS (2004) Regulation of ion channel localization and phosphorylation by neuronal activity. Nat Neurosci 7:711–718. DOI: 10.1038/nn1260nn1260 [pii].

53. Misonou H, Mohapatra DP, Menegola M & Trimmer JS (2005) Calcium- and metabolic state-dependent modulation of the voltage-dependent Kv2.1 channel regulates neuronal excitability in response to ischemia. J Neurosci 25:11184–11193. DOI: 25/48/11184 [pii]10.1523/JNEUROSCI.3370-05.2005.

54. Cerda O & Trimmer JS (2011) Activity-dependent phosphorylation of neuronal Kv2.1 potassium channels by CDK5. J Biol Chem 286:28738–28748. DOI: 10.1074/jbc.M111.251942.

55. Dong WH, Chen JC, He YL, Xu JJ & Mei YA (2013) Resveratrol inhibits K(v)2.2 currents through the estrogen receptor GPR30-mediated PKC pathway. Am J Physiol Cell Physiol 305:C547–557. DOI: 10.1152/ajpcell.00146.2013.

56. Baver SB, Hope K, Guyot S, Bjorbaek C, Kaczorowski C & O’Connell KM (2014) Leptin modulates the intrinsic excitability of AgRP/NPY neurons in the arcuate nucleus of the hypothalamus. J Neurosci 34:5486–5496. DOI: 10.1523/JNEUROSCI.4861-12.2014.

57. Hwang PM, Glatt CE, Bredt DS, Yellen G & Snyder SH (1992) A novel K+ channel with unique localizations in mammalian brain: molecular cloning and characterization. Neuron 8:473–481.

58. Hwang PM, Fotuhi M, Bredt DS, Cunningham AM & Snyder SH (1993) Contrasting immunohistochemical localizations in rat brain of two novel K+ channels of the Shab subfamily. J Neurosci 13:1569–1576.

59. Hwang PM, Cunningham AM, Peng YW & Snyder SH (1993) CDRK and DRK1 K+ channels have contrasting localizations in sensory systems. Neuroscience 55:613–620.

60. Zurek N, Sparks L & Voeltz G (2011) Reticulon short hairpin transmembrane domains are used to shape ER tubules. Traffic 12:28–41. DOI: 10.1111/j.1600-0854.2010.01134.x.

61. Day RN & Davidson MW (2009) The fluorescent protein palette: tools for cellular imaging. Chem Soc Rev 38:2887–2921. DOI: 10.1039/b901966a.

62. Besprozvannaya M, Dickson E, Li H, Ginburg KS, Bers DM, Auwerx J & Nunnari J (2018) GRAM domain proteins specialize functionally distinct ER-PM contact sites in human cells. eLife 7. DOI: 10.7554/eLife.31019.

63. Herrington J, Zhou YP, Bugianesi RM, Dulski PM, Feng Y, Warren VA, Smith MM, Kohler MG, Garsky VM, Sanchez M, Wagner M, Raphaelli K, Banerjee P, Ahaghotu C, Wunderler D, Priest BT, Mehl JT, Garcia ML, McManus OB, Kaczorowski GJ & Slaughter RS (2006) Blockers of the delayed-rectifier potassium current in pancreatic beta-cells enhance glucose-dependent insulin secretion. Diabetes 55:1034–1042.

64. Tilley DC, Eum KS, Fletcher-Taylor S, Austin DC, Dupre C, Patron LA, Garcia RL, Lam K, Yarov-Yarovoy V, Cohen BE & Sack JT (2014) Chemoselective tarantula toxins report voltage activation of wild-type ion channels in live cells. Proc Natl Acad Sci U S A 111:E4789–4796. DOI: 10.1073/pnas.1406876111.

65. Jensen CS, Watanabe S, Stas JI, Klaphaak J, Yamane A, Schmitt N, Olesen SP, Trimmer JS, Rasmussen HB & Misonou H (2017) Trafficking of Kv2.1 channels to the axon initial segment by a novel nonconventional secretory pathway. J Neurosci 37:11523–11536. DOI: 10.1523/JNEUROSCI.3510-16.2017.

66. Sanchez-Ponce D, DeFelipe J, Garrido JJ & Munoz A (2012) Developmental expression of Kv potassium channels at the axon initial segment of cultured hippocampal neurons. PLoS One 7:e48557. DOI: 10.1371/journal.pone.0048557.

67. Leterrier C (2016) The Axon Initial Segment, 50 years later: A nexus for neuronal organization and function. Curr Top Membr 77:185–233. DOI: 10.1016/bs.ctm.2015.10.005.

68. Sanchez-Ponce D, DeFelipe J, Garrido JJ & Munoz A (2011) In vitro maturation of the cisternal organelle in the hippocampal neuron’s axon initial segment. Mol Cell Neurosci 48:104–116. DOI: 10.1016/j.mcn.2011.06.010.

69. King AN, Manning CF & Trimmer JS (2014) A unique ion channel clustering domain on the axon initial segment of mammalian neurons. J Comp Neurol 522:2594–2608. DOI: 10.1002/cne.23551.

70. Schluter A, Del Turco D, Deller T, Gutzmann A, Schultz C & Engelhardt M (2017) Structural plasticity of synaptopodin in the axon initial segment during visual cortex development. Cereb Cortex 27:4662–4675. DOI: 10.1093/cercor/bhx208.

71. Spector I, Shochet NR, Kashman Y & Groweiss A (1983) Latrunculins: novel marine toxins that disrupt microfilament organization in cultured cells. Science 219:493–495.

72. Williams RT, Manji SS, Parker NJ, Hancock MS, Van Stekelenburg L, Eid JP, Senior PV, Kazenwadel JS, Shandala T, Saint R, Smith PJ & Dziadek MA (2001) Identification and characterization of the STIM (stromal interaction molecule) gene family: coding for a novel class of transmembrane proteins. Biochem J 357:673–685.

73. Soboloff J, Spassova MA, Hewavitharana T, He LP, Xu W, Johnstone LS, Dziadek MA & Gill DL (2006) STIM2 is an inhibitor of STIM1-mediated store-operated Ca2+ Entry. Curr Biol 16:1465–1470. DOI: 10.1016/j.cub.2006.05.051.

74. Brandman O, Liou J, Park WS & Meyer T (2007) STIM2 is a feedback regulator that stabilizes basal cytosolic and endoplasmic reticulum Ca2+ levels. Cell 131:1327–1339. DOI: 10.1016/j.cell.2007.11.039.

75. Shalygin A, Skopin A, Kalinina V, Zimina O, Glushankova L, Mozhayeva GN & Kaznacheyeva E (2015) STIM1 and STIM2 proteins differently regulate endogenous store-operated channels in HEK293 cells. J Biol Chem 290:4717–4727. DOI: 10.1074/jbc.M114.601856.

76. Inoue T, Heo WD, Grimley JS, Wandless TJ & Meyer T (2005) An inducible translocation strategy to rapidly activate and inhibit small GTPase signaling pathways. Nat Methods 2:415–418. DOI: 10.1038/nmeth763.

77. Lee HC, Wang JM & Swartz KJ (2003) Interaction between extracellular Hanatoxin and the resting conformation of the voltage-sensor paddle in Kv channels. Neuron 40:527–536. DOI: S0896627303006366 [pii].

78. Wu MM, Covington ED & Lewis RS (2014) Single-molecule analysis of diffusion and trapping of STIM1 and Orai1 at endoplasmic reticulum-plasma membrane junctions. Molecular biology of the cell 25:3672–3685. DOI: 10.1091/mbc.E14-06-1107.

79. VanDongen AM, Frech GC, Drewe JA, Joho RH & Brown AM (1990) Alteration and restoration of K+ channel function by deletions at the N- and C-termini. Neuron 5:433–443.

80. Misonou H, Menegola M, Mohapatra DP, Guy LK, Park KS & Trimmer JS (2006) Bidirectional activity-dependent regulation of neuronal ion channel phosphorylation. J Neurosci 26:13505–13514. DOI: 26/52/13505 [pii]10.1523/JNEUROSCI.3970-06.2006.

81. Misonou H, Thompson SM & Cai X (2008) Dynamic regulation of the Kv2.1 voltage-gated potassium channel during brain ischemia through neuroglial interaction. J Neurosci 28:8529–8538. DOI: 28/34/8529 [pii]10.1523/JNEUROSCI.1417-08.2008.

82. Romer SH, Deardorff AS & Fyffe RE (2016) Activity-dependent redistribution of Kv2.1 ion channels on rat spinal motoneurons. Physiol Rep 4. DOI: 10.14814/phy2.13039.

83. Speca DJ, Ogata G, Mandikian D, Bishop HI, Wiler SW, Eum K, Wenzel HJ, Doisy ET, Matt L, Campi KL, Golub MS, Nerbonne JM, Hell JW, Trainor BC, Sack JT, Schwartzkroin PA & Trimmer JS (2014) Deletion of the Kv2.1 delayed rectifier potassium channel leads to neuronal and behavioral hyperexcitability. Genes Brain Behav 13:394–408. DOI: 10.1111/gbb.12120.

84. Hermanstyne TO, Subedi K, Le WW, Hoffman GE, Meredith AL, Mong JA & Misonou H (2013) Kv2.2: a novel molecular target to study the role of basal forebrain GABAergic neurons in the sleep-wake cycle. Sleep 36:1839–1848. DOI: 10.5665/sleep.3212.

85. Hermanstyne TO, Kihira Y, Misono K, Deitchler A, Yanagawa Y & Misonou H (2010) Immunolocalization of the voltage-gated potassium channel Kv2.2 in GABAergic neurons in the basal forebrain of rats and mice. J Comp Neurol 518:4298–4310. DOI: 10.1002/cne.22457.

86. Guan D, Armstrong WE & Foehring RC (2013) Kv2 channels regulate firing rate in pyramidal neurons from rat sensorimotor cortex. J Physiol 591:4807–4825. DOI: 10.1113/jphysiol.2013.257253.

87. Kimm T, Khaliq ZM & Bean BP (2015) Differential regulation of action potential shape and burst-frequency firing by BK and Kv2 channels in substantia nigra dopaminergic neurons. J Neurosci 35:16404–16417. DOI: 10.1523/JNEUROSCI.5291-14.2015.

88. Benndorf K, Koopmann R, Lorra C & Pongs O (1994) Gating and conductance properties of a human delayed rectifier K+ channel expressed in frog oocytes. J Physiol 477 (Pt 1):1–14.

89. O’Connell KM, Loftus R & Tamkun MM (2010) Localization-dependent activity of the Kv2.1 delayed-rectifier K+ channel. Proc Natl Acad Sci U S A 107:12351–12356. DOI: 1003028107 [pii]10.1073/pnas.1003028107.

90. Fox PD, Loftus RJ & Tamkun MM (2013) Regulation of Kv2.1 K(+) conductance by cell surface channel density. J Neurosci 33:1259–1270. DOI: 10.1523/JNEUROSCI.3008-12.2013.

91. Kaczmarek LK (2006) Non-conducting functions of voltage-gated ion channels. Nat Rev Neurosci 7:761–771. DOI: 10.1038/nrn1988.

92. Dai XQ, Manning Fox JE, Chikvashvili D, Casimir M, Plummer G, Hajmrle C, Spigelman AF, Kin T, Singer-Lahat D, Kang Y, Shapiro AM, Gaisano HY, Lotan I & Macdonald PE (2012) The voltage-dependent potassium channel subunit Kv2.1 regulates insulin secretion from rodent and human islets independently of its electrical function. Diabetologia 55:1709–1720. DOI: 10.1007/s00125-012-2512-6.

93. Fu J, Dai X, Plummer G, Suzuki K, Bautista A, Githaka JM, Senior L, Jensen M, Greitzer-Antes D, Manning Fox JE, Gaisano HY, Newgard CB, Touret N & MacDonald PE (2017) Kv2.1 clustering contributes to insulin exocytosis and rescues human beta-cell dysfunction. Diabetes 66:1890–1900. DOI: 10.2337/db16-1170.

94. Li L, Pan ZF, Huang X, Wu BW, Li T, Kang MX, Ge RS, Hu XY, Zhang YH, Ge LJ, Zhu DY, Wu YL & Lou YJ (2016) Junctophilin 3 expresses in pancreatic beta cells and is required for glucose-stimulated insulin secretion. Cell Death Dis 7:e2275. DOI: 10.1038/cddis.2016.179.

95. Lees JA, Messa M, Sun EW, Wheeler H, Torta F, Wenk MR, De Camilli P & Reinisch KM (2017) Lipid transport by TMEM24 at ER-plasma membrane contacts regulates pulsatile insulin secretion. Science 355. DOI: 10.1126/science.aah6171.

96. Marini C, Romoli M, Parrini E, Costa C, Mei D, Mari F, Parmeggiani L, Procopio E, Metitieri T, Cellini E, Virdò S, De Vita D, Gentile M, Prontera P, Calabresi P & Guerrini R (2017) Clinical features and outcome of 6 new patients carrying de novo KCNB1 gene mutations. Neurology Genetics 3. DOI: 10.1212/nxg.0000000000000206.

97. Murakoshi H, Shi G, Scannevin RH & Trimmer JS (1997) Phosphorylation of the Kv2.1 K+ channel alters voltage-dependent activation. Mol Pharmacol 52:821–828.

98. Park KS, Mohapatra DP, Misonou H & Trimmer JS (2006) Graded regulation of the Kv2.1 potassium channel by variable phosphorylation. Science 313:976–979. DOI: 313/5789/976 [pii]10.1126/science.1124254.

99. Ikematsu N, Dallas ML, Ross FA, Lewis RW, Rafferty JN, David JA, Suman R, Peers C, Hardie DG & Evans AM (2011) Phosphorylation of the voltage-gated potassium channel Kv2.1 by AMP-activated protein kinase regulates membrane excitability. Proc Natl Acad Sci U S A 108:18132–18137. DOI: 10.1073/pnas.1106201108.

100. Frazzini V, Guarnieri S, Bomba M, Navarra R, Morabito C, Mariggio MA & Sensi SL (2016) Altered Kv2.1 functioning promotes increased excitability in hippocampal neurons of an Alzheimer’s disease mouse model. Cell Death Dis 7:e2100. DOI: 10.1038/cddis.2016.18.

101. Muennich EA & Fyffe RE (2004) Focal aggregation of voltage-gated, Kv2.1 subunit-containing, potassium channels at synaptic sites in rat spinal motoneurones. J Physiol 554:673–685. DOI: 10.1113/jphysiol.2003.056192.

102. Sharma N, D’Arcangelo G, Kleinlaus A, Halegoua S & Trimmer JS (1993) Nerve growth factor regulates the abundance and distribution of K+ channels in PC12 cells. J Cell Biol 123:1835–1843.

103. Schulien AJ, Justice JA, Di Maio R, Wills ZP, Shah NH & Aizenman E (2016) Zn(2+) - induced Ca(2+) release via ryanodine receptors triggers calcineurin-dependent redistribution of cortical neuronal Kv2.1 K(+) channels. J Physiol 594:2647–2659. DOI: 10.1113/JP272117.

104. Shah NH, Schulien AJ, Clemens K, Aizenman TD, Hageman TM, Wills ZP & Aizenman E (2014) Cyclin e1 regulates Kv2.1 channel phosphorylation and localization in neuronal ischemia. J Neurosci 34:4326–4331. DOI: 10.1523/JNEUROSCI.5184-13.2014.

105. Dunn KW, Kamocka MM & McDonald JH (2011) A practical guide to evaluating colocalization in biological microscopy. Am J Physiol Cell Physiol 300:C723–742. DOI: 10.1152/ajpcell.00462.2010.

106. Misonou H, Mohapatra DP & Trimmer JS (2005) Kv2.1: a voltage-gated K+ channel critical to dynamic control of neuronal excitability. Neurotoxicology 26:743–752. DOI: S0161-813X(05)00048-3 [pii]10.1016/j.neuro.2005.02.003.

107. Hartzell CA, Jankowska KI, Burkhardt JK & Lewis RS (2016) Calcium influx through CRAC channels controls actin organization and dynamics at the immune synapse. eLife 5. DOI: 10.7554/eLife.14850.

108. Hsieh TS, Chen YJ, Chang CL, Lee WR & Liou J (2017) Cortical actin contributes to spatial organization of ER-PM junctions. Molecular biology of the cell 28:3171–3180. DOI: 10.1091/mbc.E17-06-0377.

109. Manning CF, Bundros AM & Trimmer JS (2012) Benefits and pitfalls of secondary antibodies: why choosing the right secondary is of primary importance. PLoS One 7:e38313. DOI: 10.1371/journal.pone.0038313.

110. Bernsen J (1986) Dynamic thresholding of gray-level images. Proc. 8th Int. Conf. on Pattern Recognition, pp 1251–1255.

111. Dickson EJ, Jensen JB, Vivas O, Kruse M, Traynor-Kaplan AE & Hille B (2016) Dynamic formation of ER-PM junctions presents a lipid phosphatase to regulate phosphoinositides. J Cell Biol 213:33–48. DOI: 10.1083/jcb.201508106.

112. Ikeda SR & Korn SJ (1995) Influence of permeating ions on potassium channel block by external tetraethylammonium. J Physiol 486 (Pt 2):267–272.

113. Immke D, Wood M, Kiss L & Korn SJ (1999) Potassium-dependent changes in the conformation of the Kv2.1 potassium channel pore. J Gen Physiol 113:819–836.

114. Immke D & Korn SJ (2000) Ion-Ion interactions at the selectivity filter. Evidence from K(+)- dependent modulation of tetraethylammonium efficacy in Kv2.1 potassium channels. J Gen Physiol 115:509–518.

115. Sack JT, Aldrich RW & Gilly WF (2004) A gastropod toxin selectively slows early transitions in the Shaker K channel’s activation pathway. J Gen Physiol 123:685–696. DOI: 10.1085/jgp.200409047.

116. Thielicke W & Stamhuis EJ (2014) Towards user-friendly, affordable and accurate digital particle image velocimetry in MATLAB. J Open Res Software 2:e30. DOI: doi.org/10.5334/jors.bl.

